# Thalamus–cortex interactions drive cell type-specific cortical development in human pluripotent stem cell-derived assembloids

**DOI:** 10.1101/2025.10.11.681772

**Authors:** Masatoshi Nishimura, Shota Adachi, Tomoki Kodera, Akinori Y. Sato, Ryosuke F. Takeuchi, Fumitaka Osakada

## Abstract

The thalamus plays a pivotal role in the development and function of neural circuits in the cerebral cortex. However, how thalamus–cortex interactions influence human cortical development remains unknown primarily because of the inaccessibility of the human embryonic brain. Here, we demonstrate thalamus-dependent gene expression, circuit organization, and neural activity during corticogenesis using human thalamocortical assembloids (hThCAs). Human cortical and thalamic organoids (hCOs and hThOs) derived from induced pluripotent stem cells exhibited region-specific gene expression and spontaneous neuronal activity. Upon the fusion of these organoids, hThCAs reconstructed reciprocal thalamus–cortex axonal projections and synaptic connections. Transcriptomic analysis revealed thalamus-dependent acceleration of cortical maturation, with upregulation of programs linked to axon development, subplate/cortical plate identity, and activity-regulated genes. Histological analysis showed expanded progenitor pools and increased deep-layer neurons within hThCAs. Wide-field Ca2+ imaging demonstrated that wave-like activity originated in the thalamic region and propagated to the cortical region. Furthermore, two-photon Ca2+ imaging of cortical neurons revealed that synchronous activity emerged exclusively in pyramidal tract (PT) and corticothalamic (CT) neurons, whereas intratelencephalic (IT) neurons remain asynchronous, highlighting cell type–specific circuit integration within hThCAs. These synchronized events were absent in isolated hCOs or in cortico–cortical assembloids, underscoring the specificity of thalamic input. Our findings suggest that diffusible thalamic cues broadly enhance progenitor expansion, while long-range thalamic input organizes cell type–specific synchronous activity. This study demonstrates the thalamus-dependent acquisition of mature cortical phenotypes in a cell type-specific manner in hThCAs, establishing developmental mechanisms linking regional interactions and cell type–specific circuit specification. Thus, hThCAs provide a tractable human platform for dissecting human-specific developmental processes and modeling neurodevelopmental disorders with disrupted thalamocortical communication at the molecular, cellular, and circuit levels.

**Graphical Abstract:** 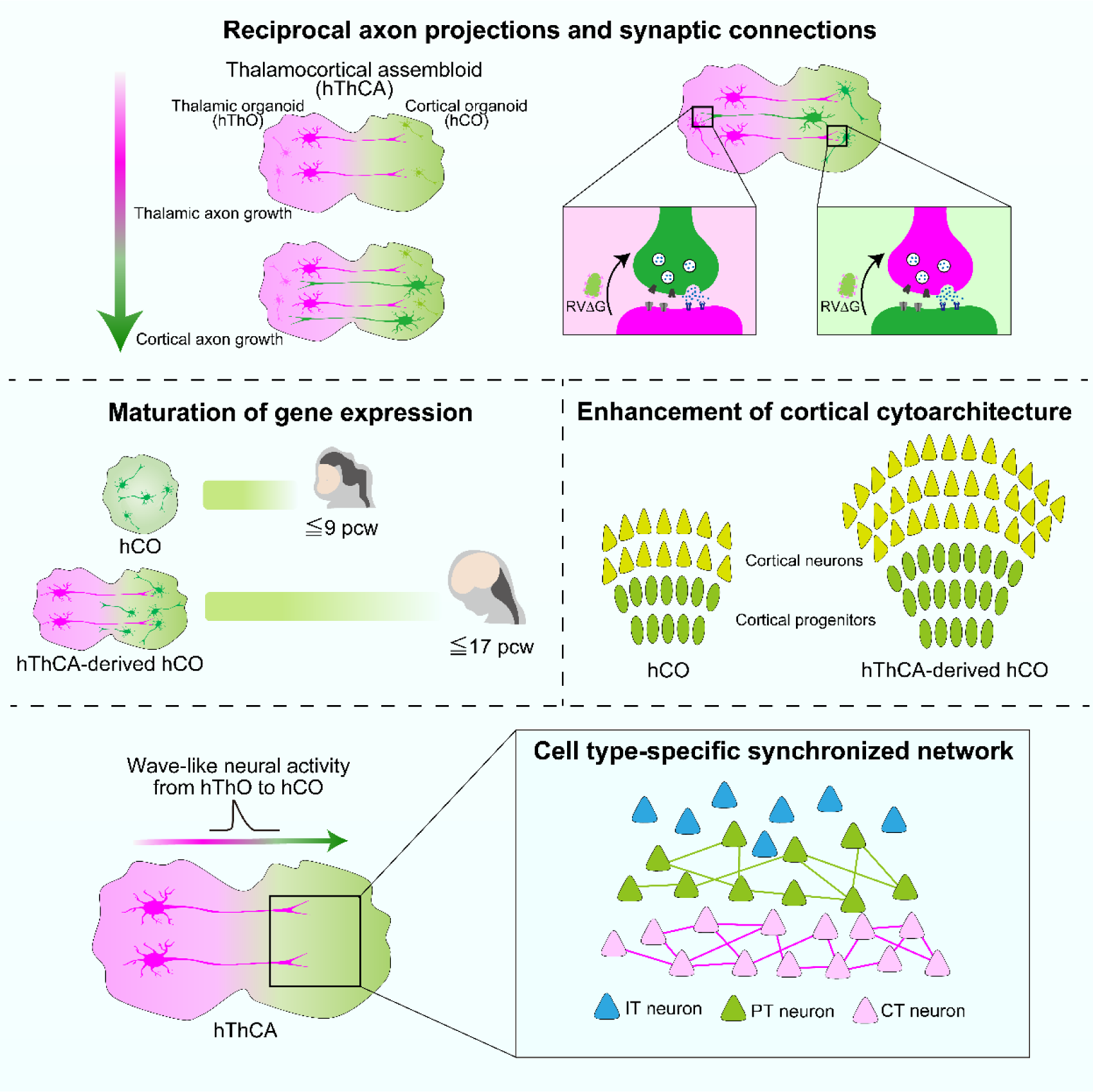

**Highlights:**

1. hThCAs recapitulate the developmental sequence of reciprocal projections with synapses.
2. hThCA cortical transcriptomes resemble the late fetal human cortex.
3. Thalamic paracrine signals promote progenitor expansion and cortical cytoarchitecture.
4. Wave-like activity originates in the thalamus and propagates across the cortex.
5. Thalamic input drives cell type–specific synchronization and circuit integration.

## Introduction

The cerebral cortex holds remarkable abilities, such as sensory perception, motor control, and cognition (1, 2). These complex functions arise from interactions between intracortical and subcortical regions via cell type-specific neural circuits (3). The cerebral cortex contains three primary subtypes of excitatory neurons: intratelencephalic (IT), pyramidal tract (PT), and corticothalamic (CT) neurons (4, 5). IT neurons project to intracortical areas and the striatum, whereas PT and CT neurons project to subcortical areas, including the thalamus. Among subcortical structures, the thalamus plays a crucial role as a relay hub, integrating sensory and motor signals and modulating cortical processing (5–7). Notably, thalamus–cortex interactions begin early in development (8). The thalamus innervates newborn cortical neurons, thereby shaping cortical organization during development (9). In rodent models, thalamic inputs influence area patterning in the cortex during development, such as the establishment of boundaries between primary and higher-order cortices and the acquisition of modality-specific identities in cortical neurons (10, 11). Furthermore, synchronous calcium waves originating in the thalamus propagate to the cortex, shaping the cortical area size and topographic somatosensory map (12, 13). These findings highlight the essential role of thalamus– cortex interactions in cortical development. However, because of the limited availability of functional human tissues, how thalamus–cortex interactions influence the development and organization of the human cortex remain poorly understood. Therefore, establishing models that recapitulate thalamus–cortex interactions in human development is essential to advancing our understanding of human cortical development and function.

Neural organoids derived from pluripotent stem cells offer a powerful *in vitro* system for studying the development of the human brain, which is otherwise difficult to access, and for testing hypotheses by enabling the manipulation of human tissues *in vitro* (14–18). While cortical organoids can recapitulate some aspects of cortical structure and function, such as neurogenesis, neural activity, and synaptogenesis (19–23), they lack interregional interactions that are essential for brain development and function. A newly developed approach that employs the assembly of multiple region-specific organoids, or neural assembloids, allows the reconstruction of interregional neural circuits in the human brain *in vitro* (24–27). Xiang *et al.* (28) generated thalamocortical assembloids by fusing cortical and thalamic organoids, thereby forming long-distance reciprocal axonal projections. More recently, Kim *et al.* (29) and Patton *et al.* (30) demonstrated long-range functional synaptic connections and synaptic plasticity, respectively, within the thalamocortical assembloids. While these thalamocortical assembloids partially recapitulate *in vivo* thalamus–cortex interactions, the specific role of thalamic inputs on human cortical circuit development at the cell type level remains unknown.

In this study, we explored thalamus-dependent mechanisms of human cortical circuit development using thalamocortical assembloids (hThCAs). We generated human cortical organoids (hCOs) and human thalamic organoids (hThOs) from human induced pluripotent stem cells (hiPSCs), each with region-specific gene expression and functional neurons. Upon fusion of hCOs and hThOs, hThCAs exhibited thalamus-dependent changes in gene expression, progenitor pools, circuit organization, and cortical activity patterns. Notably, calcium signals originating in hThOs propagated to hCOs within the hThCAs, and synchronous activity emerged exclusively in cortical neuron subtypes projecting to the thalamus. Our findings demonstrate the thalamus-dependent development of cell type-specific cortical circuits in hThCAs, highlighting cell type-specific and regional interactions as developmental mechanisms for the specialization and diversification of cortical circuits. Our human assembloid model provides a powerful platform for dissecting mechanisms underlying human cortical development, modeling neurodevelopmental disorders, and advancing drug discovery.

## Results

### Reciprocal axonal projections and synaptic connections are reconstructed in human thalamocortical assembloids

To reconstruct interactions between the thalamus and the cerebral cortex, we first generated region-specific human cortical organoids (hCOs) and thalamic organoids (hThOs) from hiPSCs using previously reported protocols (Figs. S1A and S1B) (23, 28). We characterized these induced organoids by evaluating their gene expression profiles with qPCR analysis. The hCOs showed higher expressions of the cortical markers *FOXG1*, *EMX1*, *EMX2*, *TBR1*, and *BCL11B* than the hThOs (Fig. S1C). Conversely, significantly higher expression of the diencephalic marker *OTX2* and thalamic markers *TCF7L2*, *DBX1*, *GBX2*, *OLIG3*, and *SLC17A6* was observed in the hThOs than in the hCOs (Fig. S1D). Immunohistochemistry further demonstrated the specific expression of TCF7L2 in hThOs, while FOXG1 was specifically expressed in hCOs (Figs. S1E and S1F). These results demonstrate the establishment of a cortical or thalamic fate within each organoid. The functionality of neurons in hCOs and hThOs was assessed by two-photon Ca2+ imaging with the genetically encoded Ca2+ indicator jGCaMP7f (31). jGCaMP7f was introduced to neurons in each region-specific organoid using AAVDJ-CaMKIIa-jGCaMP7f. Fluorescence signal changes in jGCaMP7f were observed in both organoids (Figs. S1G and S1H), indicating spontaneous neural activities in hCOs and hThOs.

We then utilized an assembloid approach to explore thalamus–cortex interaction *in vitro*. To distinguish each region within assembloids, two distinct lines of hiPSCs expressing either mCherry or EYFP were generated with the CRISPR-Cas9 genome editing system (Figs. S2A–S2C). hCOs and hThOs were generated from the EYFP- and mCherry-expressing hiPSC lines, respectively. The hCOs and hThOs were fused to form thalamocortical assembloids (hThCAs) (Fig. 1A). The hThCAs exclusively expressed the cortical marker FOXG1 on the EYFP+ hCO side and the thalamic marker TCF7L2 on the mCherry+ hThO side (Fig. 1B), indicating that the hThCAs retained the corresponding regional identities of the organoids.

**Figure 1.**
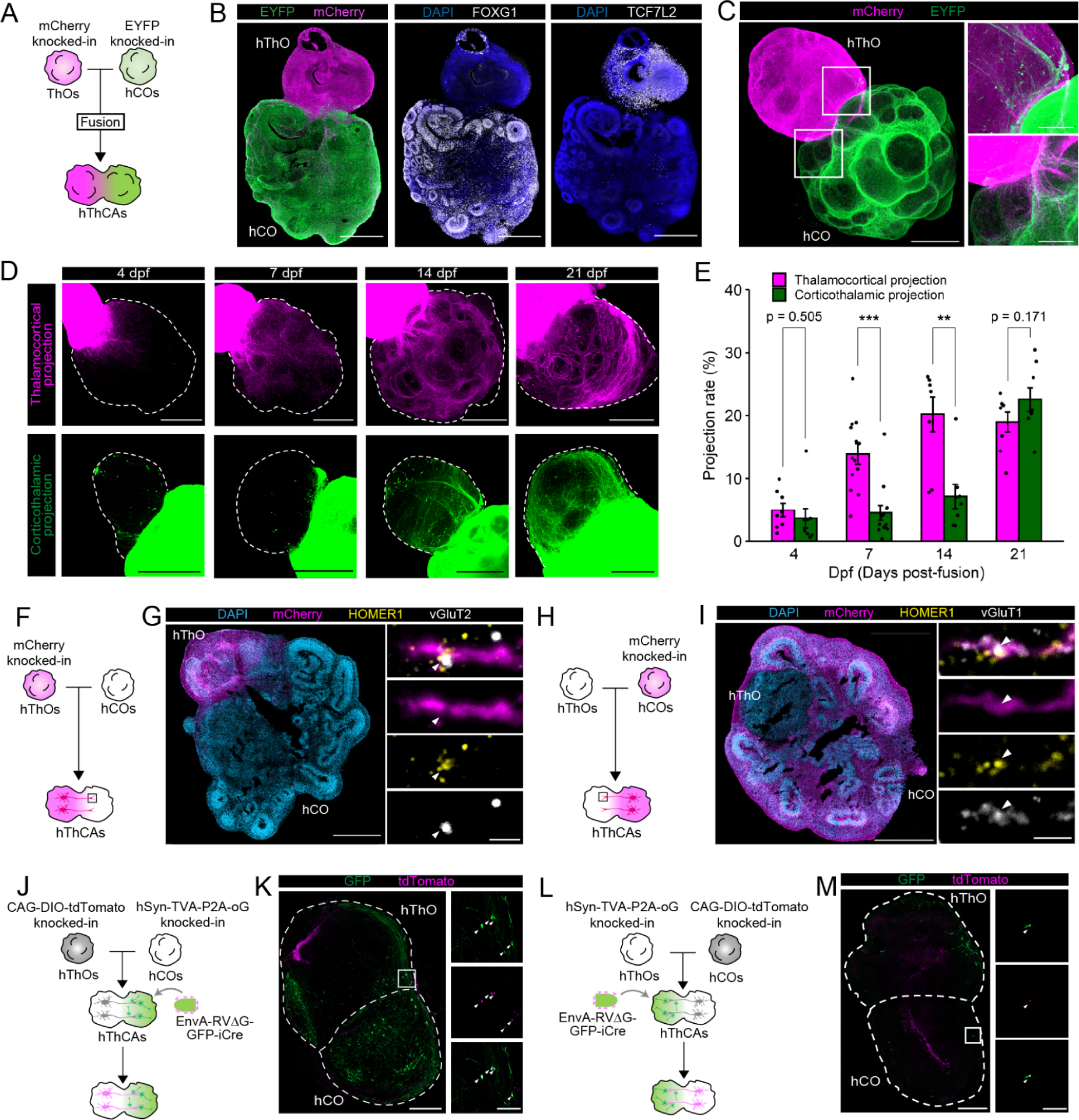
Reciprocal axonal projections and synaptic connections in hThCAs. (A) Method for generating hThCAs by fusing hThOs and hCOs. (B) Immunostaining for the cortical marker FOXG1, thalamic marker TCF7L2, and fluorescent proteins in hThCAs at 14 days post fusion (dpf). Scale bars: 500 µm. (C) Three-dimensional immunostaining of hThCAs at 14 dpf. Scale bars represent 1,000 µm (left) and 200 µm (right). (D) Reciprocal axonal projections between the cortex and thalamus within hThCAs at 4, 7, 14, and 21 dpf. Scale bar: 500 µm. (E) Quantification of axonal projections within hThCAs on 4, 7, 14, and 21 dpf. Data are expressed as mean ± SEM: n = 8 hThCAs at 4, 14, and 21 dpf, n = 13 hThCAs at 7 dpf. \*\**p* < 0.01, \*\*\**p* < 0.001, Welch’s *t*-test. (F) Generation of hThCAs from mCherry-expressing hThOs and unlabeled hCOs. (G) Immunostaining for mCherry, the presynaptic marker at thalamic axon terminals vGluT2, and the postsynaptic marker HOMER1 in the hThOs axons extending to hCOs within hThCAs at 52 dpf. Scale bars represent 1,000 µm (left) and 2 µm (right). White arrowheads, colocalization of vGluT2 and HOMER1 on mCherry+ hThOs axon. (H) Generation of hThCAs from unlabeled hThOs and mCherry-expressing hCOs. (I) Immunostaining for mCherry, the presynaptic marker at cortical axon terminals vGluT1, and the postsynaptic marker HOMER1 in hCOs axons extending to hThOs in hThCAs at 52 dpf. Scale bars represent 1,000 µm (left) and 2 µm (right). White arrowheads, colocalization of vGluT1 and HOMER1 on mCherry+ hThOs axon. (J) Method for detecting long-range thalamocortical synaptic connections in hThCAs using transsynaptic rabies tracing and KI-iPS cell lines. (K) Transsynaptic labeling of thalamocortical inputs from hThOs to hCOs within hThCAs at 42 dpf. Scale bars represent 500 µm (left) and 100 µm (right). (L) Method for detecting long-range corticothalamic synaptic connections in hThCAs using transsynaptic rabies tracing and KI-iPS cell lines. (M) Transsynaptic labeling of corticothalamic inputs from hCOs to hThOs within hThCAs at 21 dpf. Scale bars represent 500 µm (left) and 100 µm (right).

We investigated the formation of reciprocal thalamus–cortex projections within the hThCAs by performing tissue clearing using the CUBIC method (32) and 3D immunostaining for EYFP and mCherry. Fourteen days post-fusion (dpf), EYFP+ axons extended from hCOs toward hThOs, while mCherry+ axons extended from hThOs toward hCOs (Fig. 1C). The mCherry+ thalamocortical axons preferentially targeted cortical plate-like regions rather than ventricular zone-like regions (Figs. S2D–S2F), signifying interactions between thalamic and cortical neurons. In addition, mCherry+ thalamocortical axons were broadly distributed across proximal, distal, and deep cortical compartments (Figs. S2G and S2H). Notably, axonal projections from hThOs to hCOs formed earlier than those from hCOs to hThOs (Figs. 1D and 1E), which is consistent with the developmental sequence of the *in vivo* pattern where the development of feedforward thalamocortical projections preceded that of feedback corticothalamic projections in primates (8). We also generated cortico-cortical assembloids (hCCAs) as a control by fusing two hCOs and found that cell migration occurred more frequently in hCCAs than in hThCAs (Figs. S2I–S2K). Thus, the axon projection-biased interaction was a unique phenotype in the hThCAs. These results demonstrate the recapitulation of the development of *in vivo* thalamus–cortex interactions by hThCAs.

To identify reciprocal synaptic connections between the hThOs and hCOs within the hThCAs, we conducted an immunohistochemical analysis of synaptic proteins. In the mCherry- hCOs fused with mCherry+ hThOs, vGluT2, the presynaptic marker at thalamic axon terminals, and postsynaptic marker HOMER1 were colocalized on mCherry+ thalamic axons (Figs. 1F and 1G). Similarly, vGluT1, the presynaptic marker at cortical axon terminals, and postsynaptic marker HOMER1 were colocalized on the mCherry+ cortical axons that extended into the hThOs within the hThCAs (Figs. 1H and 1I).

We further investigated the long-range synaptic connections between hThOs and hCOs in hThCAs by RVΔG transsynaptic tracing. Using the CRISPR-Cas9 system, we first generated two independent iPSC lines: one with TVA and RVoG controlled by the human synapsin promoter (hiPSCs-TVA-oG) and another carrying a Cre-dependent DIO-tdTomato cassette at the AAVS1 locus (hiPSCs-DIO-tdTomato) (Figs. S3A–S3F). To verify the formation of thalamocortical synapses, hThOs and hCOs were generated from hiPSCs- DIO-tdTomato and hiPSCs-TVA-oG, respectively, and fused to create hThCAs. The hThCAs were infected with EnvA-RVΔG-GFP-iCre (Fig. 1J). Fourteen days after EnvA-RVΔG-GFP-iCre infection, we observed GFP+ cells on the hiPSCs-TVA-oG-derived hCOs side and GFP+ tdTomato+ cells on the hiPSCs-DIO-tdTomato-derived hThOs side of the hThCAs (Fig. 1K). To investigate corticothalamic synapse formation in hThCAs, we generated hThOs from hiPSCs-TVA-oG and hCOs from hiPSCs-DIO-tdTomato and created hThCAs (Fig. 1L). Following EnvA-RVΔG-GFP-iCre infection, GFP+ cells were observed in the hThOs side and GFP+tdTomato+ cells in the hCOs side of the hThCAs (Fig. 1M). Collectively, these anatomical and transsynaptic investigations suggest that both thalamocortical and corticothalamic reciprocal axonal projections and excitatory synapses were formed within the hThCAs.

### Thalamus–cortex interactions enhance the maturation of cortical gene expression in thalamocortical assembloids

To reveal the effects of thalamic inputs on cortical gene expression patterns, we conducted a comprehensive transcriptomic analysis. hThCAs were generated by fusing mCherry+ hCOs and mCherry- hThOs. At approximately day 70, following dissociation, sorted mCherry+ cells were subjected to bulk RNA sequencing (Fig. 2A). The transcriptomic profiles of mCherry+ hThCA-derived hCOs were compared with those of mCherry+ hCOs cultured alone. To assess the developmental relevance of isolated hCOs and hThCA-derived hCOs to the human brain, we compared the transcriptomes of hCOs and hThCA-derived hCOs with those of region-specific fetal brains obtained from the BrainSpan dataset (33). Both hCOs and hThCA-derived hCOs were highly correlated with the cortex (Fig. 2B) and exhibited no changes in cortical areal identity (Fig. S2A). However, Uniform Manifold Approximation and Projection (UMAP) revealed a distinct separation between the hCOs and hThCA-derived hCOs (Fig. 2C), suggesting changes in gene expression patterns driven by thalamic inputs. Analysis of differentially expressed genes between the hCOs and the hThCA-derived hCOs identified 414 upregulated genes and 776 downregulated genes in hThCA-derived hCOs compared to hCOs (Fig. 2D). Gene ontology (GO) analysis of the upregulated genes in the hThCA-derived hCOs revealed enrichment in categories related to axon development and synaptic signaling (Fig. 2E), whereas downregulated genes were associated with sensory organ and nephron development (Fig. S4B). The genes associated with the cortical plate and subplate, regions that receive thalamic input during early development (34), were upregulated in hThCA-derived hCOs (Fig. 2F). While most primate-specific genes (35) showed comparable expression levels between hCOs and hThCA-derived hCOs (Fig. S4C), the expression levels of *OSTN* and its receptor *NPR3* were higher in hThCA-derived hCOs (Fig. 2G). *OSTN* encodes OSTEOCRIN, a primate-specific activity-dependent secreted factor expressed in cortical neurons driven by thalamic inputs that regulates dendritic growth (36). These results suggest that functional inputs from hThOs to hCOs drive activity-dependent gene expression, leading to the formation of mature cortical circuits within hThCAs.

**Figure 2.**
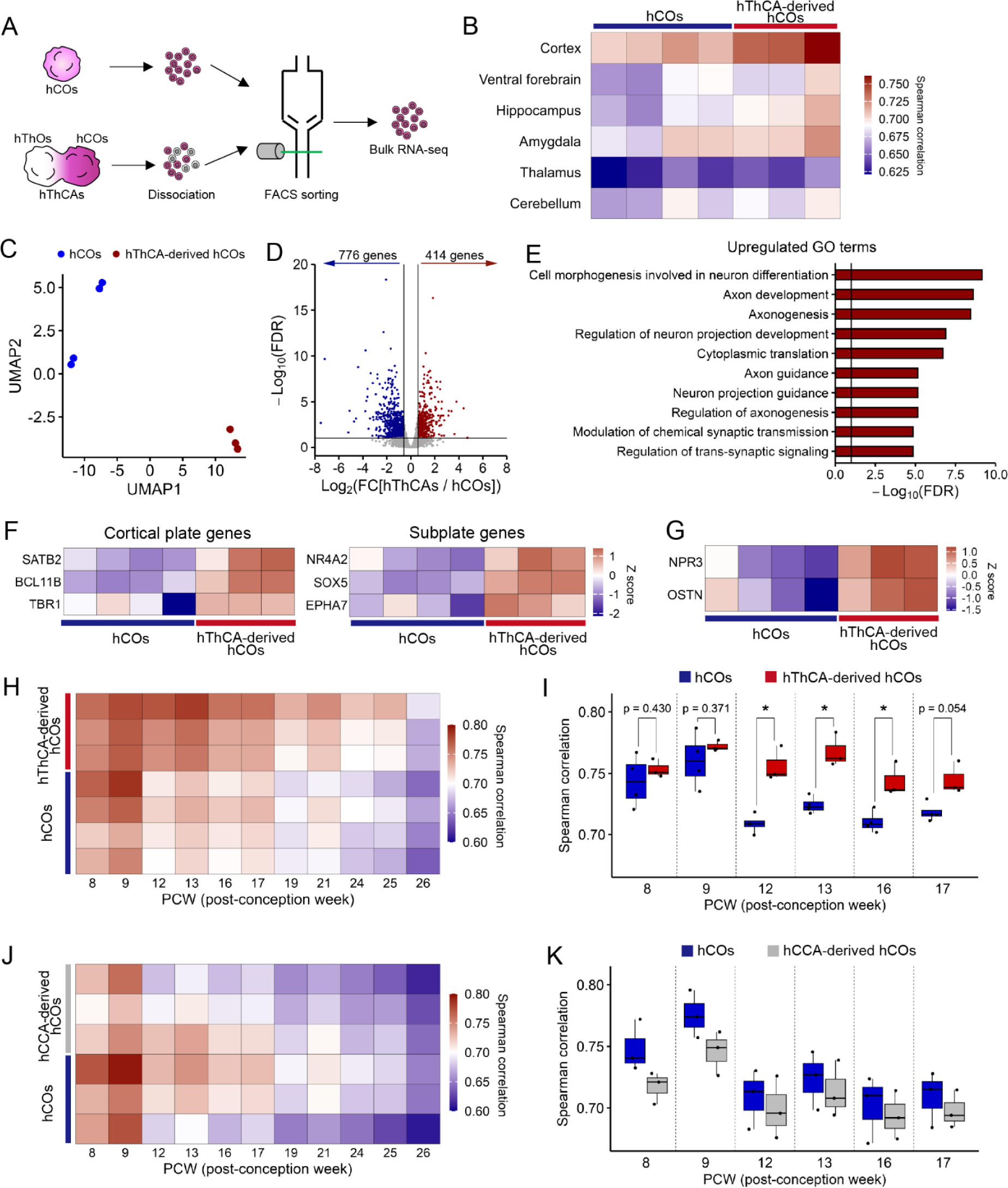
Gene expression patterns of isolated hCOs and hThCA-derived hCOs. (A) Bulk RNA sequencing method for hCOs and hThCA-derived hCOs. (B) Heatmap showing the Spearman correlations between the transcriptomes of region-specific human fetal brains and those of hCOs or hThCA-derived hCOs. n = 4 hCOs, n = 3 hThCA- derived hCOs. (C) Uniform Manifold Approximation and Projection (UMAP) for the transcriptomes of hCOs and hThCA-derived hCOs. n = 4 hCOs, n = 3 hThCA-derived hCOs. (D) Volcano plot showing the differentially expressed genes between hCOs and hThCA-derived hCOs. Vertical lines represent the fold-change = 1.5 or −1.5; horizontal lines represent the false discovery rate (−log10FDR) = 1. (E) Top 10 gene ontology terms upregulated in hThCA-derived hCOs. Vertical line represents −log10FDR = 1. (F) Heatmaps showing the expression of cortical plate genes (left) and subplate genes (right) in hCOs and hThCA-derived hCOs. n = 4 hCOs, n = 3 hThCA-derived hCOs. (G) Heatmap showing the expression of *OSTN* and *NPR3* in hCOs and hThCA-derived hCOs. n = 4 hCOs, n = 3 hThCA-derived hCOs. (H) Heatmap showing the Spearman correlations between the transcriptomes of human fetal brain and those of hCOs or hThCA-derived hCOs. n = 4 hCOs, n = 3 hThCA-derived hCOs. (I) Box plot showing the Spearman correlations between the transcriptomes of human fetal brain and those of hCOs or hThCA-derived hCOs. n = 4 hCOs, n = 3 hThCA-derived hCOs. \**p* < 0.05, Welch’s *t*-test. (J) Heatmap showing the Spearman correlations between the transcriptomes of human fetal brain and those of hCOs or hCCA-derived hCOs. n = 4 hCOs, n = 3 hThCA-derived hCOs. (K) Box plot showing the Spearman correlations between the transcriptomes of human fetal brain and those of hCOs or hCCA-derived hCOs. n = 3 hCOs, n = 3 hCCA-derived hCOs.

To further assess maturation stages of hCOs and hThCA-derived hCOs, we compared the transcriptomes of hCOs and hThCA-derived hCOs with those of the fetal cerebral cortex at different developmental stages. Isolated hCOs were highly correlated with the human fetal cortex at 8–9 post-conception weeks (pcw), whereas hThCA-derived hCOs exhibited high similarity to the human fetal cortex up to 17 pcw (Figs. 2H and 2I). To determine whether the maturation of hThCA-derived hCOs is specifically dependent on thalamus–cortex interactions, we generated cortico-cortical assembloids (hCCAs) by fusing two hCOs and analyzed the transcriptomes of hCCA-derived hCOs (Fig. S4D). The gene expression patterns of hCCA-derived hCOs also correlated with the fetal cerebral cortex (Fig. S4E). However, UMAP did not reveal a distinct separation between hCOs and hCCA-derived hCOs (Fig. S4F). Furthermore, the correlation between hCCA-derived hCOs and the fetal cerebral cortex were limited to the 8–9 pcw stage, like isolated hCOs (Figs. 2J and 2K). Therefore, thalamic inputs, not fusion *per se*, are essential for driving the maturation of cortical gene expression in hCOs.

### Thalamocortical assembloids exhibit increased progenitor pools and cortical neurons

Thalamus–cortex interactions promote the proliferation of cortical precursors and enhance the generation of cortical neurons during early development (37). To investigate these effects in the hThCAs, we examined the cellular compositions of isolated hCOs and hThCA-derived hCOs. Immunostaining revealed the formation of neural rosettes, in which PAX6+ progenitor cells were radially organized outside the N-CADHERIN+ apical membrane, in both hCOs and hThCA-derived hCOs (Fig. 3A). However, hThCA-derived hCOs exhibited a significantly greater number of neural rosettes compared to hCOs (Fig. 3B). Additionally, the thickness and area of individual rosettes were significantly increased in hThCA-derived hCOs (Figs. 3C and 3D). In contrast, the enlargement of neural rosettes was not observed in hCCA-derived hCOs (Figs. S5A–S5D), suggesting that cortical progenitor expansion is a thalamus-specific effect rather than a consequence of fusion. We then evaluated the abundance of proliferative and intermediate progenitors in hCOs and hThCA-derived hCOs. Although the proportion of PAX6+ progenitors remained comparable between hCOs and hThCA-derived hCOs, the fractions of Ki67+ proliferating cells and TBR2+ intermediate progenitors in hThCA-derived hCOs were significantly higher than those in hCOs (Figs. 3E and 3F). Thus, thalamus–cortex interactions promote the proliferation of cortical precursors within hThCAs.

**Figure 3.**
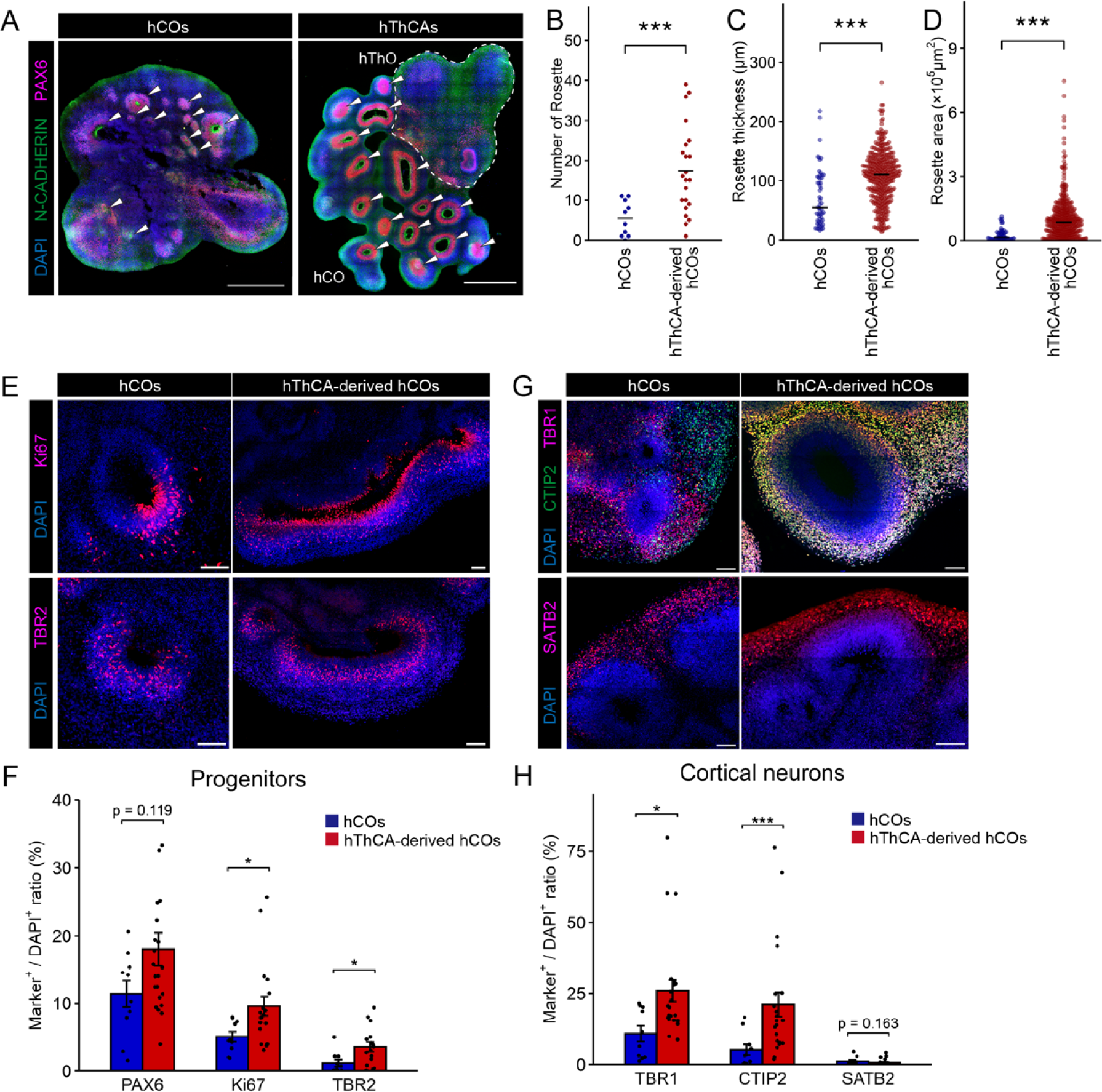
Cytoarchitecture in isolated hCOs and hThCA-derived hCOs. (A) Immunostaining for N-CADHERIN and PAX6 in hCOs and hThCAs on day 80. Scale bars represent 1,000 µm. White arrowheads represent neural rosettes. (B) Quantification of the number of rosettes in hCOs and hThCA-derived hCOs. n = 10 hCOs, n =22 hThCA-derived hCOs. \*\*\**p* < 0.001, Welch’s *t*-test. (C) Quantification of the thickness of each rosette in hCOs and hThCA-derived hCOs. n = 55 rosettes (hCOs), n = 384 rosettes (hThCA-derived hCOs). \*\*\**p* < 0.001, Mann–Whitney U test. (D) Quantification of the area of each rosette in hCOs and hThCA-derived hCOs. n = 55 rosettes (hCOs), n = 384 rosettes (hThCA-derived hCOs). \*\*\**p* < 0.001, Mann–Whitney U test. (E) Immunostaining for Ki67 or TBR2 in hCOs and hThCA-derived hCOs on day 80. Scale bars represent 100 µm. (F) Quantification of PAX6, Ki67, and TBR2 expression in hCOs and hThCA-derived hCOs. Data are expressed as mean ± SEM. n = 10 hCOs, n = 22, 19, or 16 (PAX6, Ki67, or TBR2) hThCA- derived hCOs. \**p* < 0.05, Mann–Whitney U test. (G) Immunostaining for TBR1 and CTIP2 or SATB2 in hCOs and hThCA-derived hCOs on day 80. Scale bars represent 100 µm. (H) Quantification of TBR1, CTIP2, and SATB2 expression in hCOs and hThCA-derived hCOs. Data are expressed as mean ± SEM. n = 10 hCOs, n = 22 hThCA-derived hCOs. \**p* < 0.05, \*\*\**p* < 0.001, Mann–Whitney U test.

We next examined whether this progenitor expansion was mediated by direct tissue contact or indirect interactions via soluble factors. Coculture hCOs with hThOs in the same dish without physical fusion significantly increased the number and area of neural rosettes in hCOs (Figs. S5A–S5D), suggesting that non-contact, paracrine signaling from the thalamus contributed to progenitor pool expansion.

The expansion of the progenitor pool in hThCA-derived hCOs prompted an investigation of potential differences in the abundance of cortical neurons between hCOs and hThCA-derived hCOs. Immunostaining for cortical neuron markers revealed that the proportions of TBR1+ and CTIP2+ deep-layer neurons were significantly increased in hThCA-derived hCOs compared to hCOs, hCCAs, and cocultured hCOs (Figs. 3G, 3H, S5E, and S5F). Although SATB2+ upper-layer neurons were also observed in hThCA-derived hCOs, there was no observable difference in their abundance between hCOs and hThCA-derived hCOs (Figs. 3G and 3H). Collectively, these results indicate that thalamus–cortex interactions promote progenitor proliferation and enhance the generation of deep-layer cortical neurons via diffusible cues.

### Cortical neurons exhibit synchronous activity only when fused with thalamic organoids

In the developing brain, thalamic waves originate in the thalamus and propagate to the cortex, regulating cortical development (12). To determine whether similar thalamocortical functional interactions occur in hThCAs, we examined neuronal activity in hThCAs comprising hCOs and mCherry+ hThOs. The hThCAs were infected with RVΔG-GCaMP6s and subjected to wide-field Ca2+ imaging at 5 days post-infection (Figs. 4A and 4B). GCaMP6s fluorescence signals generated in hThOs propagated to hCOs and elicited synchronous cortical activity within the hThCAs (Figs. 4C and 4D), suggesting that hThCAs recapitulate thalamus-to-cortex propagation observed *in vivo*.

**Figure 4.**
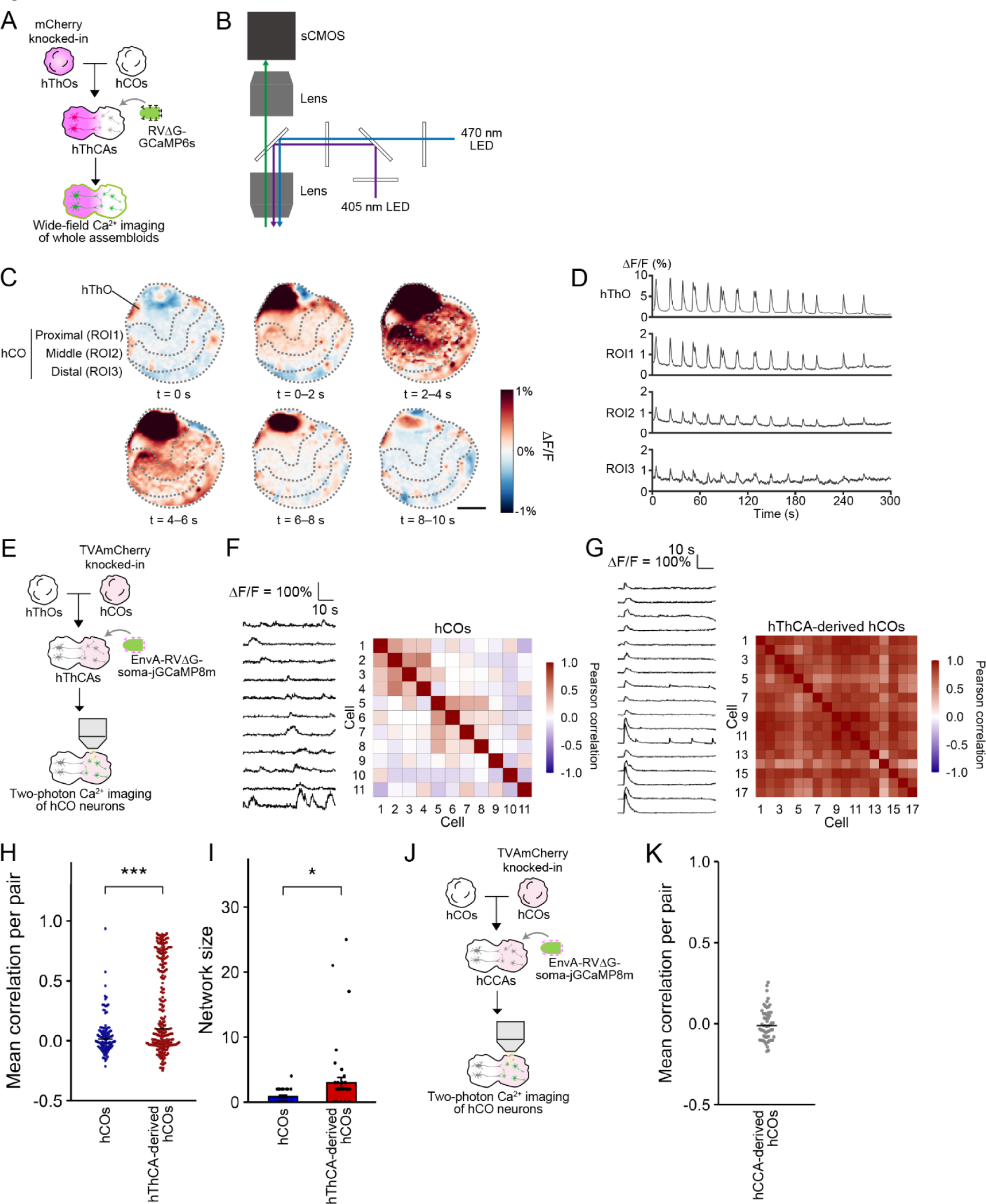
Thalamus-dependent synchronous neural activity within hThCAs. (A) Method of the wide-field Ca2+ imaging for hThCAs. (B) Schematic of the wide-field imaging system. (C) GCaMP6s fluorescence changes in hThCA. The gray dashed box represents each region. Each image represents the mean of 80 frames (2 s). Scale bar represents 1,000 µm. (D) Representative time-series traces of GCaMP6s signal changes in each region in hThCA. (E) Method for two-photon Ca2+ imaging of cortical neurons in hThCAs. (F) Representative time-series traces of jGCaMP8m signal changes (left) and heatmap showing correlation coefficients of activity between all pairs of simultaneously recorded neurons in hCOs (right). (G) Representative time-series traces of synchronous jGCaMP8m signal changes (left) and heatmap showing correlation coefficients of activity between all pairs of simultaneously recorded neurons in hThCAs (right). (H) Quantification of mean correlation coefficients of activity between all pairs of simultaneously recorded neurons from hCOs and hThCAs. Blue and red dots, correlation of each cell pair; black line, median. n = 96 pairs (hCOs), n = 233 pairs (hThCA-derived hCOs). \*\*\**p* < 0.001, Mann–Whitney U test. (I) Quantification of the network synchronization size in hCOs and hThCAs. Data are expressed as mean ± SEM. n = 28 network (hCOs), n = 42 network (hThCA-derived hCOs). \**p* < 0.05, Mann–Whitney U test. (J) Method for two-photon Ca2+ imaging of cortical neurons in hCCAs. (K) Quantification of mean correlation coefficients of activity between all pairs of simultaneously recorded neurons from hCCAs. n = 96 pairs.

To further investigate cortical neural activity at the cellular level, we performed two-photon Ca2+ imaging for hThCAs. We first established a hiPSC line expressing TVA, a receptor for the viral envelope EnvA (38), under the human synapsin promoter using the CRISPR-Cas9 system (hiPSCs-TVA) (Figs. S6A–S6C). hCOs were generated from these hiPSCs-TVA and subsequently fused with TVA-negative hThOs to generate hThCAs. Cortical neurons within the hThCAs were selectively labeled with jGCaMP8m (39) by infection of the hThCAs with EnvA-pseudotyped rabies virus vectors encoding soma-jGCaMP8m (EnvA-RVΔG-soma-jGCaMP8m) (40) (Fig. S6D). Both hThCAs and hCOs were infected with EnvA-RVΔG-soma-jGCaMP8m and subjected to two-photon Ca2+ imaging 5 days post-infection (Fig. 4E). While both isolated hCOs and hThCA-derived hCOs exhibited spontaneous neural activity, only hThCA-derived hCOs showed synchronous neural activity (Figs. 4F and 4G). To quantify and compare the activity synchronization of cortical neurons between hCOs and hThCAs, we calculated the mean correlation coefficients between cells (27). We found significantly increased activity synchronization of neuron pairs in hThCAs compared to isolated hCOs (Fig. 4H). In addition, the network size, defined as the number of synchronized cells, was significantly larger in hThCAs than in hCOs (Fig. 4I). To determine whether this synchronous neural activity was specifically dependent on thalamus–cortex interactions, we developed hCCAs as a control by fusing normal hCOs and hiPSCs-TVA-derived hCOs. Following EnvA-RVΔG-soma-jGCaMP8m infection, we performed two-photon Ca2+ imaging of hCCAs (Fig. 4J). In contrast to hThCAs, cortical neurons in hCCAs exhibited asynchronous neural activity, indicating that synchronous neural activity observed in hThCAs is thalamus-dependent (Fig. 4K).

To determine the role of hThOs in cortical activity within hThCAs, we selectively suppressed synaptic transmissions from the thalamus to the cortex using the tetanus toxin light chain (TeLC) (41, 42). We generated TVA-expressing ThOs, fused them with hCOs to form hThCAs, and infected the hThCAs with EnvA-RVΔG-TeLC-DsRed (Fig. S7A). The TVA-expressing thalamic cells expressed TeLC (Fig. S7B), resulting in the suppression of synaptic vesicle exocytosis within the hThCAs. The TeLC expression did not markedly alter the structure or cell composition of hThCA-derived hCOs (Fig. S7C–S7G). Two-photon Ca2+ imaging of the cortical compartment within the hThCAs revealed that suppression of synaptic transmissions from the thalamus reduced activity synchronization of neuron pairs and tended to decrease coactive ensembles in the cortex (Fig. S7H–S7K). Taken together, these results indicate that thalamic activity drives wave-like propagation into the cortex and that thalamus-cortex synaptic interactions are necessary for the generation of correlated cortical activity with a large network in hThCAs.

### Cell type-specific synchronous neural activity emerges in thalamocortical assembloids

The cerebral cortex comprises a heterogeneous cell population of excitatory neurons, which can be classified into three types: intratelencephalic (IT), pyramidal tract (PT), and corticothalamic (CT) neurons (Fig. S8A). To determine whether these excitatory neuron subtypes develop in our system, we generated AAVs expressing EGFP or SYFP2 under control of the validated subtype-selective enhancers mscRE16, mscRE4, and mscRE10 for IT, PT, and CT neurons (AAV9-IT/PT/CTenhancer-EGFP or SYFP2), respectively (43) (Fig. S8B). Fourteen days after AAVs infections, fluorescence signals of EGFP or SYFP2 in hCOs were detected in hCOs (Figs. S8C and S8D). To validate the cell-type specificity of the enhancer constructs used to label IT, PT, and CT populations, we labeled cortical neurons in hCOs and hThCA-derived hCOs with GFP driven by each subtype-specific enhancer, isolated GFP⁺ cells using a cell sorter, and subjected the purified populations to RNA seqencing (Fig. S8E). The dimensional reduction by UMAP revealed distinct separation among IT-, PT-, and CT-labeled populations in hThCA-derived hCOs, whereas each subtype in isolated hCOs was not well separated into distinct clusters (Fig. S8F and S8G), suggesting that thalamic interactions sharpen cell type identity and specification in the cortex. These results indicate that these AAV-enhancers labeled molecularly distinct subtypes of cortical projection neurons in hThCAs.

To identify which excitatory neuron subtypes contribute to synchronous neural activity in hThCAs, we performed two-photon Ca2+ imaging of IT, PT, and CT neurons. We generated AAVs that encode the TVA gene under the control of subtype-specific enhancers (AAV9-IT/PT/ITenhancer-TVA) (Fig. 5A). Following AAV infection, hThCAs were subsequently infected with EnvA-RVΔG-soma-jGCaMP8m for targeted expression of the calcium indicator in cortical neuron subtypes (Fig. 5B). Two-photon Ca2+ imaging revealed that IT neurons exhibited asynchronous activity, whereas PT and CT neurons showed significantly high synchronous activity in hThCAs (Figs 5C–E). Furthermore, the mean correlation coefficients between cells for each subtype revealed significantly increased activity synchronization of PT and CT neurons in hThCAs (Fig. 5F). In contrast, cell type-specific two-photon Ca2+ imaging of hCOs and hCCAs revealed no synchronous activity in any of the three cortical neuron subtypes, indicating that synchronized dynamics require thalamic interaction (Figs. 5G, 5H, and Figs. S9A–S9C). Taken together, these results suggest that thalamic interactions refine projection neuron subtype identity and cell type-specific cortical circuits in hThCAs, with PT and CT neurons displaying synchronized activity.

**Figure 5.**
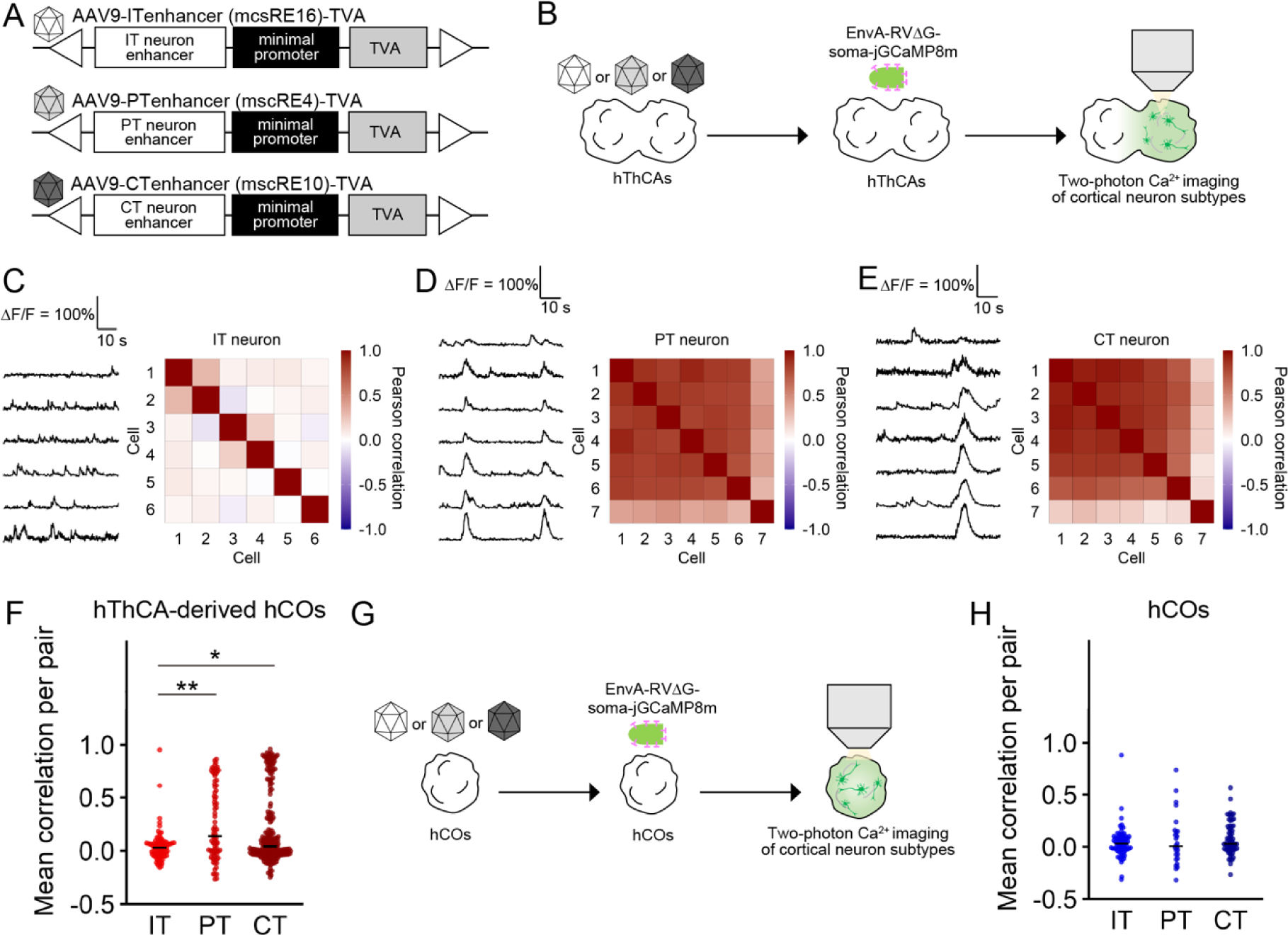
Cell type-specific two-photon Ca^2+^ imaging of isolated hCOs and hThCA-derived hCOs. (A) AAVs for expressing TVA under the control of mscRE16, mscRE4, and mscRE10 enhancers specific for intratelencephalic (IT), pyramidal tract (PT), and corticothalamic (CT) neurons, respectively. (B) Method for cell type-specific two-photon Ca2+ imaging of hThCA-derived hCOs. (C) Representative time-series traces of jGCaMP8m signal changes (left) and heatmap showing the correlation coefficients of activity between all pairs of simultaneously recorded IT neurons in hThCA-derived hCOs (right). (D) Representative time-series traces of jGCaMP8m signal changes (left) and heatmap showing correlation coefficients of activity between all pairs of simultaneously recorded PT neurons in hThCA-derived hCOs (right). (E) Representative time-series traces of jGCaMP8m signal changes (left) and heatmap showing correlation coefficients of activity between all pairs of simultaneously recorded CT neurons in hThCA-derived hCOs (right). (F) Quantification of the mean correlation coefficients of the activity between all pairs of simultaneously recorded IT, PT, or CT neurons from hThCA-derived hCOs. n = 87 pairs (IT), n = 81 pairs (PT), and n = 223 pairs (CT). \**p* < 0.05, \*\**p* < 0.01, Dunn test. (G) Method for cell type-specific two-photon Ca2+ imaging of hCOs. (H) Quantification of the mean correlation coefficients of the activity between all pairs of simultaneously recorded IT, PT, or CT neurons from hCOs. n = 79 pairs (IT), n = 26 pairs (PT), and n = 65 pairs (CT). *p* = 0.332, Kruskal-Wallis rank sum test.

## Discussion

In this study, we addressed the influence of thalamus–cortex interactions on human cortical development using hiPSC-derived hThCAs. hThCAs formed *in vivo-*like reciprocal axonal projections and long-range synaptic connections between hCOs and hThOs. Transcriptomic analysis revealed that hCOs within the hThCAs acquired characteristics of late-stage cortical identity. In addition, hThCAs showed abundant cortical progenitors and neurons, indicating that thalamic interactions contribute to cortical cytoarchitecture. Wide-field and two-photon Ca2+ imaging demonstrated the thalamus-dependent emergence of synchronous cortical activity patterns in specific cell types in hThCAs. Thus, we conclude that thalamic inputs drive cortical circuit specification and maturation in a cell type-specific manner during development. These findings suggest that regional interactions orchestrate the acquisition of mature phenotypes via cell type-specific interactions, leading to the development of functional neural circuits. This interplay between regional and cell type-specific interactions may constitute developmental mechanisms underlying the specialization and diversification of the brain.

### Recapitulation of *in vivo* thalamus–cortex interactions in hThCAs

In the developing primate brain, newborn thalamic neurons initiate axonal projections toward the cortex at the onset of the neurogenesis of cortical layer VI neurons (before 7 pcw in humans) (8). These thalamic projections innervate the cortical subplate approximately at the peak of layer VI neurogenesis (8.5–11 pcw in humans) (8). Subsequently, cortical neurons send feedback projections to the thalamus, establishing a thalamocortical loop between the developing thalamus and cortex (44). In our study, we fused hThOs with hCOs on days 22–24, when TBR1+ cortical layer VI neurons begin to develop in the hCOs. Axonal projections from the hThOs to the hCOs began to develop 4 days post-fusion, specifically extending to cortical neurons while excluding progenitors. Notably, following the peak of the development of feedforward projections from the hThOs to the hCOs, the development of feedback projections from the hCOs to the hThOs significantly increased. Transcriptomic analysis further revealed that hCOs within hThCAs exhibited maturation signatures corresponding to the embryonic cortex at 12–17 pcw, a developmental stage characterized by significant thalamic input (8), which is consistent with the developmental stage at which the cortex receives thalamic inputs. Thus, thalamocortical interactions in hThCAs occur in a spatiotemporal pattern that closely recapitulates *in vivo* brain development.

In addition to the spatiotemporal dynamics of interactions, the precise targeting of thalamic axons during early brain development is governed by coordinated molecular mechanisms (9, 45). Various guidance molecules secreted from the cortex and ganglionic eminences are pivotal in directing the long-distance journey of thalamic axons (45–47). In our study, GO analysis of upregulated genes in hThCAs revealed enrichment in categories related to “axon guidance” and “neuron projection guidance”. Thus, similar molecular mechanisms driving axonal projections *in vivo* may actively govern axonal projections and connectivity in hThCAs. Manipulation of the genes identified in this study warrant further investigation into the detailed molecular mechanisms underlying human thalamus–cortex interactions.

Despite this molecular evidence, the structural organization of thalamocortical projections in hThCAs remains incomplete. Thalamic axons reached cortical compartments, but lacked fasciculated bundles and appeared randomly distributed. The enrichment of axon-guidance pathways in our transcriptomic data indicates that hThCA-derived hCOs possess intrinsic pathfinding machinery, but extrinsic guidance cues may be missing. In the developing brain, ganglionic eminence (GE)-derived interneurons form a “corridor” that serves as a scaffold for thalamocortical axon (TCA) navigation (46). The absence of GE-derived signals in our current model likely causes the lack of axonal fasciculation observed in hThCAs. To more faithfully model the thalamocortical pathway, future studies should generate tripartite assembloids that include GE organoids positioned between hThOs and hCOs. Such a configuration would enable the examination of corridor formation, TCA fasciculation, and bundle organization under controlled conditions, providing a quantitative and mechanistic framework to investigate corridor-dependent TCA guidance in human brain development.

### Thalamus-dependent cortical development in hThCAs

In the developing brain, cortical arealization is initially defined by spatial gradient expression patterns of specific transcription factors, such as SP8 and COUP-TF1, along the anterior-posterior and medial-lateral axes (48, 49). This patterning occurs independently of the thalamus and is subsequently refined by thalamic input (48–50). Although RNA sequencing analyses revealed the upregulation of genes associated with the subplate and cortical plate and the maturation toward later developmental stages in hThCA-derived hCOs, both hCOs and hThCA-derived hCOs did not show high transcriptional similarity to multiple cortical areas and exhibited no changes in areal identity between hCOs and hThCA-derived hCOs. This finding is consistent with the previous study demonstrating that cortical excitatory neurons in cortical organoids possessed area-specific gene expression, but lacked spatial areal organization inside cortical organoids (51). Taken together, these results support the notion that thalamic input alone is insufficient to establish spatial arealization. Thus, to organize cortical regions in cortical organoids, upstream spatial patterning of these transcription factors is likely required before the cerebral cortex receives thalamic inputs.

In addition to areal specification, thalamic inputs influence cortical growth and cytoarchitecture. Previous studies demonstrated that secreted factors from thalamic axons promote progenitor proliferation, neurogenesis, cell survival, and dendritic growth in the cortex (37, 52). In our study, immunohistochemical analysis revealed expanded progenitor pools and increased cortical neuron abundance in hThCA-derived hCOs. To elucidate the mechanisms of thalamic influence on cortical development, we distinguished between indirect paracrine signaling and direct synaptic interactions. Coculture of hCOs and hThOs in the same dish without physical fusion increased neural rosette formation in the hCOs, suggesting that non-contact, paracrine signaling from the thalamus are sufficient to drive progenitor pool expansion. This finding aligns with prior *ex vivo* studies showing that thalamic axon terminals release growth factors that enhance cortical progenitor proliferation (37). In contrast, selective suppression of thalamic synaptic transmission using TeLC did not negatively alter progenitor pool size, suggesting that synaptic activity is not necessary for progenitor expansion.

At later stages, thalamic inputs orchestrate the emergence of functional cortical circuits through synchronous activity. In mouse models, highly synchronized spontaneous activity originates as calcium waves from the developing thalamus and propagates to the cortex (11, 12). Similarly, wide-field Ca2+ imaging demonstrated wave-like synchronous activity transmitted from hThOs to hCOs, and two-photon Ca2+ imaging revealed that synchronous neural activity in cortical neurons emerged within hThCAs. These synchronous events were evident among cortical neuron subtypes projecting to the thalamus. Importantly, these activity patterns did not occur in isolated hCOs or cortico-cortical assembloids (hCCAs), suggesting the specificity of thalamic influence on synchronous cortical activity. Furthermore, TeLC-mediated suppression of thalamic synaptic transmission reduced mean pairwise correlations and decreased coactive ensemble size, demonstrating that synaptic thalamic input is essential for cortical network organization. Consistently, our transcriptomic analyses identified upregulation of activity-regulated, thalamus-responsive genes (e.g., OSTN, NPR3) (36) in hThCA-derived hCOs, linking thalamus-driven synchrony to activity-dependent transcription and neuronal maturation.

Taken together, these findings suggest dual, temporally coordinated mechanisms by which thalamic influences shape human cortical development. Paracrine thalamic signals expand progenitor pools and enhance cortical architecture maturation, while synaptic thalamic interactions subsequently organize cell type-specific circuit formation and activity-dependent gene expression in the cortex. Further studies will be required to dissect these mechanisms, providing deeper insights into how thalamic signals orchestrate cortical development at the molecular, cellular, and circuit levels.

### Cell type-specific formation of cortical circuits in hThCAs

Cortical functions depend on the cortico-thalamo-cortical loop (CTC loop), a fundamental circuit composed of distinct neuron subtypes. At least two types of CTC loops have been identified, namely, the simple loop and complex loop (5). In the simple loop, monosynaptic connections are formed directly between thalamic neurons and PT or CT neurons. In contrast, the complex loop consists of IT-mediated polysynaptic connections formed between thalamic neurons and PT or CT neurons. These CTC loops have been implicated in cortical connectivity and function, as well as in the pathophysiology of psychiatric disorders (53, 54). Therefore, modeling cell type-specific human thalamus–cortex interactions is essential for elucidating mechanisms of cortical function and psychiatric disorders.

In our hThCAs, synchronized events arise selectively in PT and CT neurons, whereas IT neurons remain asynchronous. In isolated hCOs, none of the three classes exhibit synchrony. These findings suggest that thalamic drive is required to initiate ensemble synchrony in the developing cortex and that the simple loop is the primary substrate for early-stage cortical synchronization. The absence of IT neuron synchrony further implies that the complex loop via IT networks is weak or not yet established under our current conditions. Considering the temporal sequence of neuronal birth and maturation (55), it is possible that early thalamic inputs first couple to the corticofugal circuits comprising PT and CT neurons in hThCAs, while IT-mediated complex-loop dynamics emerge later as intracortical connectivity matures. Further investigations, such as cell type-specific rabies viral tracing, would uncover the formation of cell type-specific CTC loops in hThCAs, providing cell type-level insights into the developmental assembly of human thalamocortical circuits.

### Perspective

The present study provides constructive understandings of human brain development and neurodevelopmental disorders, including autism spectrum disorder, intellectual disability, and epilepsy, using human assembloids derived from pluripotent stem cells. Our *in vitro* human assembloid model enables the investigation of the molecular, cellular, and circuit mechanisms as well as human-specific mechanisms underlying human cortical development mediated by thalamic interactions. Disruptions of thalamus–cortex interactions are implicated in sensory and cognitive deficits observed in neurodevelopmental disorders (56–58). Our model offers a powerful platform to investigate pathophysiological processes and mechanisms caused by genetic and environmental factors during human cortical circuit formation, thereby advancing both our understanding of human brain development and neurodevelopmental disorders and the development of therapeutic interventions for these disorders.

## Methods

All experiments in this study were approved by Nagoya University and in accordance with the Guidelines of Nagoya University.

### Human induced pluripotent stem cell culture

The human induced pluripotent stem cell (hiPSC) lines (clone: 1383D6) were provided by Dr. Masato Nakagawa and Dr. Shinya Yamanaka of the Center for iPS Cell Research, Kyoto University. The human iPS cells were maintained on iMatrix 511 (Nippi Inc., Tokyo, Japan)-coated culture dishes in StemFit AK02N (Ajinomoto, Tokyo, Japan), as described previously (23, 59). For passaging, when hiPSCs grew to sub-confluency, hiPS cells were treated with accutase (Nacalai Tesque, Kyoto, Japan), dissociated into single cells, and replated at a density of 1.0 × 103 cells/cm2 onto iMatrix 511-coated dishes in AK02N in the presence of 10 μM Y-27632 (Wako, Osaka, Japan) for 24 h. The medium was changed to a fresh one without Y-27632 on the next day of plating and every day thereafter. hiPSCs were maintained in a humidified atmosphere of 5% CO2 at 37°C.

### Generation of human cortical organoids and human thalamic organoids

Human cortical organoids (hCOs) were generated according to a previously published protocol (23). Briefly, single-cell suspensions of undifferentiated hiPSCs were plated into a low-adherent 96-well V-bottom plate (10,000 cells/well) in Dulbecco’s Modified Eagle’s Medium (DMEM)/Ham’s F12 + KnockOut Serum Replacement (KSR) medium (DMEM/F12, 15% v/v KSR, 1% v/v Minimum Essential Medium Non-Essential Amino Acid [MEM-NEAA], 1% v/v L-glutamine, and 100 µM β-mercaptoethanol) supplemented with 100 nM LDN-193189, 500 nM A-83-01, 2 µM IWR-1 *endo*, and 50 µM Y27632. Half-medium changes were performed every other day. Y27632 was removed on day 4. After 10 days, The organoids were transferred to a spinning culture (70 rpm) in a low-adherent 6-cm dish and maintained in DMEM/F12+Neurobasal medium (1:1 mixture of DMEM/F12 and Neurobasal medium, 0.5% v/v N2 supplement, 1% v/v B27 supplement minus vitamin A, 0.5% v/v MEM-NEAA, 1% v/v L-glutamine, 0.025% v/v insulin, 50 µM β-mercaptoethanol, 10 µg/mL ciprofloxacin hydrochloride monohydrate, and 1% v/v penicillin/streptomycin) supplemented with 20 ng/mL basic fibroblast growth factor (bFGF). The medium was changed every other day until day 18. From day 18, organoids were maintained in DMEM/F12+Neurobasal medium supplemented with 20 ng/mL brain-derived neurotrophic factor (BDNF) and 200 µM ascorbic acid, with medium changes every other day thereafter.

Human thalamic organoids (hThOs) were generated according to a previously published protocol (28) with some modifications. Briefly, single-cell suspensions of undifferentiated hiPSCs were plated into a low-adherent 96-well V-bottom plate (10,000 cells/well) in DMEM/F12+KSR medium (DMEM/F12, 15% v/v KSR, 1% v/v MEM-NEAA, 1% v/v L-glutamine, and 100 µM β-mercaptoethanol) supplemented with 100 nM LDN-193189, 500 nM A-83-01, 4 µg/mL insulin, and 50 µM Y27632. Half-medium changes were performed every other day. Y27632 was removed on day 4. After 8 days, organoids were transferred to a spinning culture (70 rpm) in a low-adherent 6-cm dish and maintained in DMEM/F12+N2+B27 medium (DMEM/F12, 0.15% w/v D[+]-glucose, 100 µM β-mercaptoethanol, 1% v/v N2 supplement, and 2% v/v B27 supplement minus vitamin A) supplemented with 30 ng/mL BMP7 and 1 µM PD325901. The medium was changed every other day until day 16. From day 16, organoids were maintained in DMEM/F12+Neurobasal medium supplemented with 20 ng/mL BDNF and 200 µM ascorbic acid, with medium changes every other day thereafter.

### Generation of human assembloids

Human assembloids were generated using the same strategy according to previously published protocols (60, 61). Briefly, for human thalamocortical assembloids (hThCAs), hCOs and hThOs were collected on days 22–24 and fused by mixing them in a 1.5 mL low-binding tube for 3 days. The 1.5 mL low-binding tube was filled with 1 mL DMEM/F12+Neurobasal medium supplemented with 20 ng/mL BDNF and 200 µM ascorbic acid. The medium was gently changed 2 days post-fusion (dpf). On 3 dpf, the assembloids were transferred to a low-adherent 6-well plate or 6-cm dish in spinning culture (70 rpm). The same procedure was performed for cortico-cortical assembloids (hCCAs).

### Generation of Knocked-in iPSC lines using the CRISPR-Cas9 system

Knocked-in hiPSC lines were generated as described previously (61) with some modifications. The CRISPR-Cas9 system was used to knock-in a specific gene at the AAVS1 locus of hiPSCs. Guide RNA (gRNA) plasmids were constructed by assembling AflII-digested pCAG-mCherry-gRNA (Addgene, plasmid 87110) with gRNA sequences using NEBuilder HiFi DNA Assembly (NEB). Donor DNA plasmids were constructed by assembling BamHI and HindIII-digested pUC19 vectors with left and right homology arm (HA) fragments, and specific gene fragments (CAG-mCherry, CAG-EYFP, hSyn-TVA-P2A-oG, CAG-DIO-tdTomato, hSyn-TVA, or hSyn-TVA-mCherry). Left and right HAs were amplified by polymerase chain reaction (PCR) from the genome extracted from hiPSCs. Other fragments were obtained by PCR amplification from appropriate plasmids. hiPSCs (1.0 × 106 cells, 100 mm dish) were transfected with 2.5 µg gRNA expression vector, 2.5 µg Cas9 expression vector (pCAG-1BPNLS-Cas9-1BPNLS-2A-GFP; Addgene, plasmid 87109), and 5.0 µg donor plasmid using 20 μL Lipofectamine Stem (Thermo Fisher Scientific) according to the manufacturer’s protocol. To obtain two knocked-in hiPSC lines (CAG-mCherry and CAG-EYFP), the hiPSCs were dissociated into single cells 5 days after transfection and replated into a 100-mm dish at 2.0 × 104 cells. Five days after replating, single hiPSC colonies were manually selected based on mCherry or EYFP expression. To obtain other knocked-in hiPSC lines (hSyn-TVA-P2A-oG, CAG-DIO-tdTomato, or hSyn-TVA-mCherry), the hiPSCs were treated with 500 µg/mL G418 sulfate 2 days after transfection. Five days after G418 treatment, the hiPSCs were dissociated into single cells, all of which were replated into a 100-mm dish. Approximately 7 days after replating, single hiPSC colonies were manually selected. These hiPSC clones were genotyped by PCR with the primers listed in Table S1. Knocked-in hiPSCs were subsequently expanded and stocked.

### Quantitative reverse transcription PCR

Gene expression levels were evaluated by quantitative PCR (qPCR) as described previously (23, 59, 61). Total RNA was isolated using the FavorPrep Tissue Total RNA Purification Mini Kit. Subsequently, 500 ng of the total RNA was reverse-transcribed using the PrimeScript RT Master Mix (Takara). The synthesized cDNA was analyzed by RT-qPCR using the TB Green Fast qPCR Mix (Takara) on a LightCycler system (Roche). The primers used in the present study are listed in Table S2.

### Sectioning, immunostaining, and histological analysis

Organoids and assembloids were fixed with 4% paraformaldehyde (PFA) in phosphate-buffered saline (PBS) at 4°C overnight and washed three times with PBS at room temperature. Fixed samples were transferred into 15% sucrose in PBS for two overnight incubations, followed by 30% sucrose in PBS, as described previously (61). Organoids/assembloids were transferred to cryomold (Sakura Finetek) and embedded in a 2:1 mixture of optimal cutting temperature compound (Sakura Finetek) and 30% sucrose solution. Blocks were cut into 20-to 40-µm sections using a cryostat (CM3050S, Leica).

Immunostaining was performed as previously described (23, 61, 62). The organoid sections were reacted with Blocking One (Nacalai Tesque) for 1 h at room temperature and then with primary antibodies overnight at 4°C. The sections were washed with PBS three times and incubated with secondary antibodies for 90 minutes at room temperature (22-25°C). The primary antibodies used and their working dilutions were listed in Table S3. Cell nuclei were counterstained with 4′,6-diamidino-2-phenylindole (DAPI; 1.25 µg/mL).

Specimens were imaged with a confocal microscope (LSM800, Zeiss) and a fluorescence macroscope (Thunder, Leica). Histological data was analyzed by Fiji (v2.9.0, NIH). Neural rosettes were identified as radially organized cell clusters based on DAPI staining and the expression of PAX6 (radial progenitors) and N-CADHERIN (apical membranes). For quantification, the marker-positive ratio was calculated as the marker-positive area divided by the DAPI-positive area.

### Two-photon calcium imaging

Two-photon calcium imaging was performed as previously described (23, 61). The lower half of the organoids and assembloids was embedded in 4% low-melting-point agarose filled with oxygenated artificial cerebrospinal fluid (ACSF) consisting of 125 mM NaCl, 2.5 mM KCl, 1.25 mM NaH2PO4•H2O, 25 mM NaHCO3, 25 mM D-glucose, 2 mM CaCl2, and 1 mM MgCl2. Imaging was performed using a two-photon fluorescent microscope equipped with GaAsP-type non-descanned detector and resonant scanner (A1R- MP+, Nikon)(63). A 920-nm pulse laser (InSight DeepSee+, Spectra-Physics) was used to excite jGCaMP7f and jGCaMP8m. Time-lapse images were acquired using a 25× water immersion lens (NA: 1.1, Nikon) at a resolution of 512 × 512 pixels at 30.0 Hz. For the pharmaceutical approach, 50 µM CNQX and MK-801 were added in the ACSF during the imaging period and washed three times with ACSF.

MATLAB (Mathworks, R2022b) was used for all image processing and statistical quantification. In the first step of image processing of the time-lapse data, cross-correlation–based rigid image registration (64) was performed to correct the image displacement caused by motion artifacts. Changes in fluorescence intensity in individual cells was quantified. Regions of interest of individual cells were semi-manually defined using the “Cell Magic Wand” plugin in ImageJ (65). To reduce high-frequency artifact effects caused by electrical noise, fluorescence data of jGCaMP7f and jGCaMP8m were smoothed using Savitzky–Golay (filter width = 301 frames) and sliding moving average (filter width = 7 frames) filters, respectively. The normalized fluorescence signal change (ΔF/F0) was computed to evaluate the activity of each neuron. The baseline of each cell (F0) was computed using a low-pass percentile filter (10 percentile; cut-off frequency = 1/15 Hz). We determined the active cell based on three criteria calculated from ΔF/F. First, the skewness of the ΔF/F distribution during recording had to be >1.0. Second, the standard deviation of ΔF/F had to be >0.04. Third, at least one calcium event had to be detected during the recording period. The calcium event in jGCaMP8m was determined as the peak with “MinPeakHeight = 0.3,” “MinPeakDistance = 15,” and “MinPeakWidth = 7.5” using the MATLAB findpeaks function. The mean correlation was calculated as the average correlation coefficient of all active cell pairs for each imaging site. Network size was defined as the number of nodes in the correlation network for each imaging site. In the network, each node represents a cell, and each edge represents a correlation coefficient greater than 0.4.

For hThO silencing in hThCAs, hThCAs that expressed TVA on the hThO side were infected with EnvA-RVΔG-TeLC-DsRedX. Twenty-nine to fifty days after the EnvA-RVΔG delivery, the hThCAs were then infected with RVΔG-soma-jGCaMP8m. Two-photon Ca2+ imaging was performed for the hCO side in the hThCAs two to three days after the RVΔG infection.

### Wide-field calcium imaging

Wide-field imaging from assembloids was performed using a custom-built macroscope with tandem lenses (Plan Apo 1x, WD = 61.5 mm, Leica) and a sCMOS camera (ORCA-Fusion, Hamamatsu Photonics) (66). Assembloids were generated by fusing mCherry+ hThOs and mCherry- hCOs, infected with RVΔG-GCMP6s, and embedded in slide glass with a hole filled with ACSF consisting of 125 mM NaCl, 2.5 mM KCl, 1.25 mM NaH2PO4•H2O, 25 mM NaHCO3, 25 mM D-glucose, 2 mM CaCl2, and 1 mM MgCl2, and covered with a cover glass. The excitation light was provided by alternating blue (M470L4, Thorlabs) and violet (M405L3, Thorlabs) LEDs at 20 Hz each.

Image processing was performed using MATLAB (MathWorks, R2022b). After motion correction, images were binned to half resolution to reduce data size. To correct for artifacts and photobleaching, the signal excited at 405 nm was subtracted from the signal excited at 470 nm. The relative fluorescence change (ΔF/F) was then computed using the 50th percentile of the entire time series as the baseline.

### Tissue clearing

The assembloids were subjected to the CUBIC method (67), followed by immunohistochemistry. Briefly, assembloids were fixed with 4% PFA in PBS at 4°C overnight. The next day, assembloids were washed with PBS three times and incubated with CUBIC Reagent-1 diluted 2-fold with H2O for 3–18 h and then with Reagent-1 overnight at 37°C. Assembloids were washed with PBS three times for 90 minutes and then stained with anti-GFP (1:1,000, chicken, Abcam) and anti-mCherry (1:1,000, rat, Thermo Fisher Scientific) antibodies in PBS containing 0.1% Triton X-100 and 5% Blocking One for 2 days at 37°C. The assembloids were subsequently washed with PBS three times for 90 min and incubated with secondary antibodies for 2 days at 37°C. For refractive index matching, assembloids were incubated with CUBIC Reagent-2 diluted 2-fold for 3–18 h at room temperature and then with Reagent-2 overnight at room temperature.

Confocal imaging of assembloids was performed using a 10× objective lens at a depth of approximately 0–300 μm. The projection rate was defined as the percentage of EYFP coverage in hThOs and of mCherry coverage in hCOs with Fiji (ImageJ, v2.9.0, NIH), as previously described (27).

### Adeno-associated virus production

Adeno-associated virus (AAV) vectors were generated in HEK293T cells, as described previously (68). Briefly, AAVDJ was produced by HEK293T cells transfected with pHelper, the AAVDJ rep/cap vector, and the genomic vector pAAV-CaMKIIa-jGCaMP7f. AAV9 was produced by HEK293T cells transfected with the pDP9 vector and the genomic vector pAAV-3×(core)mscRE4-minCMV-SYFP2 (43), pAAV-mscRE10-minBGpromoter-EGFP (43), pAAV-mscRE16-minBGpromoter-EGFP (43), pAAV-3×(core)mscRE4-minCMV-TVA (this study), pAAV-mscRE10-minBGpromoter-TVA (this study), or pAAV-mscRE16-minBGpromoter-TVA (this study). AAVDJ was produced by transfecting HEK293T cells with pHelper, the AAVDJ rep/cap vector, and the genomic vector pAAV-3×(core)mscRE4-minCMV-TVA, pAAV-mscRE10-minBGpromoter-TVA, or pAAV-mscRE16-minBGpromoter-TVA. The transfected cells were harvested 3 days after transfection and lysed via freeze-and-thaw cycles for purification. After centrifugation, the supernatant was loaded onto gradients (15%, 25%, 40%, and 58%) of iodixanol OptiPrep (Serumwerk Bernburg). After centrifugation at 16,000 ×*g* at 4°C for 4 h, 200 µL of the 40% iodixanol fraction was collected and used for infection experiments. AAV genomic titers were quantified by qPCR. The titers of AAVs were listed in Table S4. Virus aliquots were stored at −80°C until use.

### G-deleted rabies virus production

EnvA-pseudotyped G-deleted rabies viral (RVΔG) vectors were produced as previously described (40, 68, 69). Briefly, RVΔGs were recovered by transfecting B7GG cells with pcDNA-SAD-B19N, pcDNA-SAD-B19P, pcDNA-SAD-B19L, pcDNA-SAD-B19G, and the rabies viral genome vector pSAD-B-19ΔG-GFP, pSAD-B19ΔG-GFP-iCre, pSAD-B19ΔG-GCaMP6s, pSAD-B19ΔG-soma-jGCaMP8m, pSAD-B19ΔG-DsRedX, or pSAD-B19ΔG-TeLC-DsRedX using Lipofectamine 2000 (Thermo Fisher Scientific). For the pseudotyping of RVΔG with EnvA, BHK-EnvA cells were infected with unpseudotyped RVΔG. The virus-containing medium was concentrated by two rounds of ultracentrifugation (XE-90, Beckman-Coulter). The infectious titers in HEK-TVA cells were then determined. HEK293T cells were used to check for contamination with unpseudotyped RVΔG. Virus aliquots were stored at −80°C until use. The titer of RV was listed in Table S4.

### Organoid and assembloid dissociation

Organoid and assembloid dissociation was performed according to a previously published protocol (70) with some modifications. Briefly, organoids were collected and washed once with PBS and incubated with TrypLE Select (Thermo Fisher Scientific) diluted 2-fold with 1 mM EDTA solution for 30 min at 37°C. Organoids and assembloids were dissected into small aggregates by pipetting and then incubated for another 15 min at 37°C to complete dissociation. After stopping the enzymatic reaction with FACS buffer, 2% v/v fetal bovine serum with PBS, the dissociated cells were centrifuged at 200 ×*g* for 5 min. The cell pellet was resuspended in the FACS buffer and filtered through a 40-µm cell strainer. The prepared sample was immediately used for the cell sorting experiment.

For subtype validation, organoids and assembloids were dissociated with Papain Dissociation system (Worthington Biochemical Corporation) according to manufacturer’s instructions. All solutions used were oxygenated with 95% O2 and 5% CO2 for 5–10 min before use. Organoids and assembloids were collected, washed once with EBSS, dissected into small pieces, and then incubated in oxygenated papain solution added with DNase for 1 h at 37°C, with gentle shaking every 20 min. Gentle pipetting was then performed to complete dissociation and the dissociated cells were centrifuged at 300 ×*g* for 5 min. The cell pellet was resuspended in the EBSS containing albumin-ovomucoid inhibitor and DNase, and the dissociated cells were centrifuged at 70 ×*g* for 6 min. The cell pellet was resuspended in the FACS buffer and filtered through a 40-µm cell strainer. The prepared sample was immediately used for the cell sorting experiment.

### Rabies virus tracing

For retrograde transsynaptic tracing in assembloids, hThCAs that expressed the TVA and oG on one side were infected with EnvA-RVΔG-GFP-iCre. Seven to 14 days after rabies viral vector infection, hThCAs were fixed with 4% PFA in PBS at 4°C overnight and processed for immunostaining.

### Bulk RNA sequencing

hThCAs, hCCAs, and hCOs were collected for bulk RNA sequencing. The hThCAs and hCCAs expressing mCherry on the hCO side (one hCOs in hCCAs) and the mCherry+ hCOs were dissociated and subjected to a cell sorter (CytoFLEX SRT, Beckman) to collect the mCherry+ cortical cell population. For enhancer validation, hThCAs and hCOs were infected with AAVDJs that encode TVA under the control of each subtype enhancer. Seven to fifteen days after AAV delivery, the hThCAs and hCOs were infected with EnvA-RVΔG-GFP to label TVA-expressing cortical cells. Three days after rabies infection, hCOs and hThCAs were dissociated and subjected to a cell sorter to collect GFP+ cortical cell populations.

Total RNA was isolated using the FavorPrep Tissue Total RNA Purification Mini Kit. cDNA libraries were prepared using the NEBNext Poly(A) mRNA Magnetic Isolation Module and NEBNext Ultra II RNA Library Prep Kit for Illumina and sequenced using the NextSeq550 (Illumina) to obtain 81-bp single-end reads. The estimated counts were generated with the salmon package (v1.10.3) (71) using the transcript index from GRCh38. The low-expression genes were filtered out using filterByExpr, and normalization of trimmed mean of M-values (TMM) was performed using edgeR (v4.2.1). For enhancer validation, the batch effect was removed with combat treatment (19). UMAP was performed using the TMM normalized read counts (counts per million). Differentially expressed genes between hThCA-derived hCOs and isolated hCOs were identified by false discovery rate (FDR) < 0.1 using the likelihood ratio test and 1.5 fold change. Gene ontology (GO) enrichment analysis was performed using the ClusterProfiler package (v4.12.2). FDR for each GO term was calculated using the Benjamini–Hochberg method. GO terms with FDR < 0.1 were defined as statistically significant. Spearman correlation with BrainSpan data was calculated for all combinations of the organoid or BrainSpan data matching the labeled conditions using only genes common to both experiments.

### Quantification and statistical analysis

Values are expressed as mean ± standard error of the mean (SEM) unless otherwise stated. For statistical analyses, data were pooled from at least two independent sets of experiments. Statistical analyses were performed using R (v4.4.0). Details on statistical analyses are also found in each figure legend. Probability values <5% were considered significant. The time window of organoid and assembloid for each analysis was described in Table S5.

## Acknowledgments

We thank members of the Osakada Laboratory for their valuable discussions; Mr. Hanada, Mr. Nishimura, Mr. Kano, Mr. Kato, and Mr. Kobayashi (Research Equipment Development Group, Technical Center, Nagoya University) for manufacturing instruments; and Dr. Nomoto and Ms. Akama (Center for Gene Research, Nagoya University) for assistance with bulk RNA sequencing. This work was supported by the Grants-in-Aid from the Japan Society for the Promotion of Science (F.O.; 20K21476, 21H05168, and 22H02771), PRESTO and CREST from the Japan Science and Technology Agency (F.O.; JPMJPR14F6 and JPMJCR1851), CREST and Brain/MINDS 2.0 from Japan Agency for Medical Research and Development (F.O.; JP23gm1510011 and JP24wm0625110), and SRF Foundation (F.O.). M.N. is a recipient of Nagoya University Interdisciplinary Frontier Fellowship.

---

**Figure S1.**
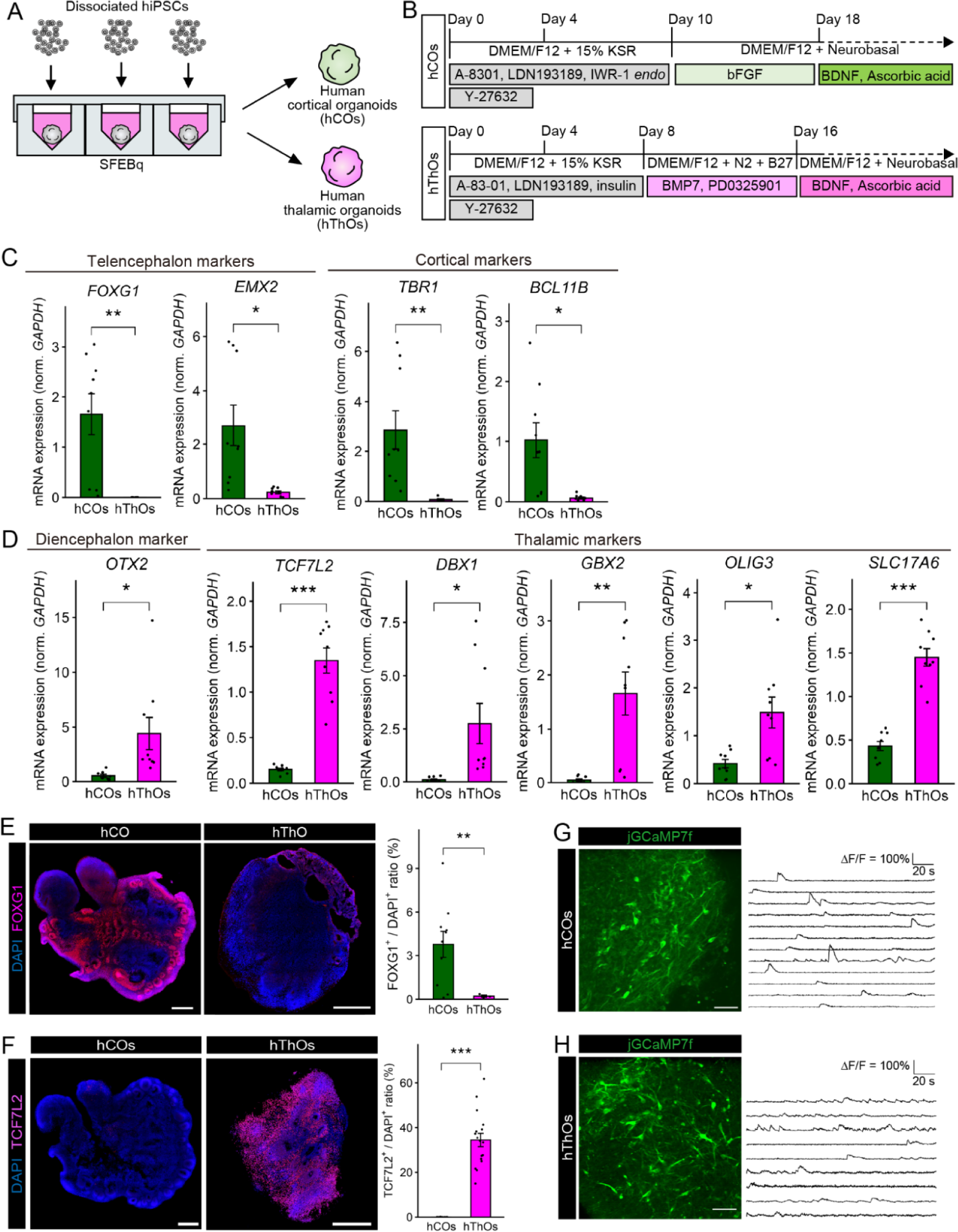
Characterization of hCOs and hThOs. (A) Schematic of generating region-specific organoids from hiPSCs. (B) Protocol of the stepwise treatment for generating hCOs and hThOs from hiPSCs. (C) qPCR analysis of telencephalic and cortical markers in hCOs and hThOs on day 42. Data are expressed as mean ± SEM. n = 9 hCOs, n = 9 hThOs, \**p* < 0.05, \*\**p* < 0.01, \*\*\**p* < 0.001, Welch’s *t*-test. (D) qPCR analysis of diencephalic and thalamic markers in hCOs and hThOs on day 42. Data are expressed as mean ± SEM. n = 9 hCOs and n = 9 hThOs, \**p* < 0.05, \*\**p* < 0.01, \*\*\**p* < 0.001, Welch’s *t*-test. (E) Immunostaining for FOXG1 (left) and quantification of FOXG1 expression in hCOs and hThOs (right) on day 42. Data are expressed as mean ± SEM: n = 13 hCOs, n = 3 hThOs. Scale bar represents 500 µm. \*\**p* < 0.01, Welch’s *t*-test. (F) Immunostaining for TCF7L2 (left) and quantification of TCF7L2 expression in hCOs and hThOs (right) on day 42. Data are expressed as mean ± SEM: n = 11 hCOs, n = 17 hThOs. Scale bar represents 500 µm. \*\*\**p* < 0.001, Welch’s *t*-test. (G) Representative images and time-series traces of jGCaMP7f signal changes in hCOs on days 50. Scale bar represents 50 µm. (H) Representative images and time-series traces of jGCaMP7f signal changes in hThOs on days 50. Scale bar represents 50 µm.

**Figure S2.**
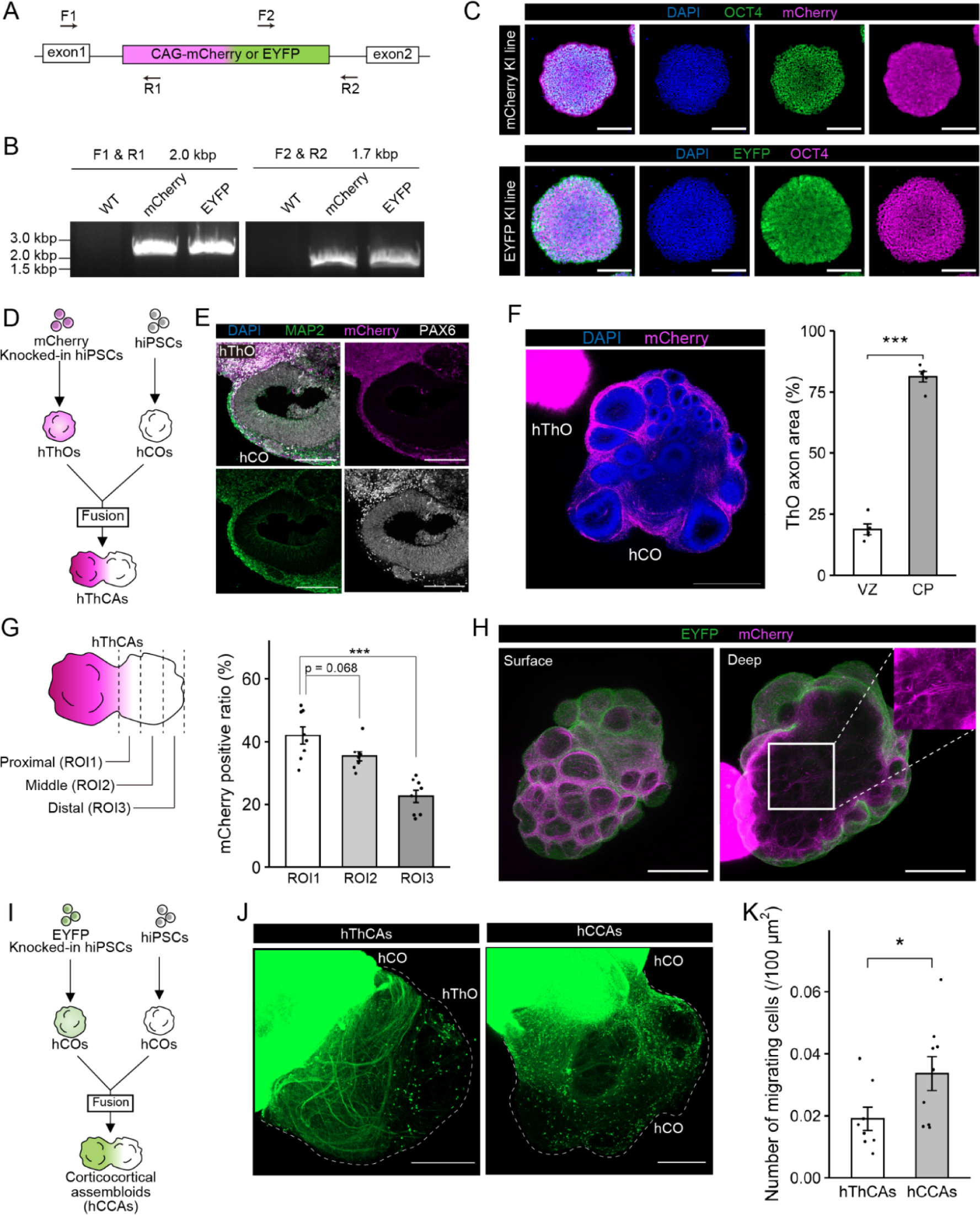
Characterization of reciprocal projections within hThCAs. (A) Primers for genotyping mCherry or EYFP knocked-in iPSC lines. Arrows, PCR primers. (B) Electrophoresis images of the PCR products for mCherry-KI or EYFP-KI iPSC lines. (C) Immunostaining for the pluripotent marker OCT4 and fluorescent proteins in mCherry-KI or EYFP-KI hiPSC lines. Scale bars represent 200 µm. (D) Generation of hThCAs from mCherry-expressing hThOs and unlabeled hCOs. (E) Immunostaining for mCherry, the neuronal marker MAP2, and the progenitor marker PAX6 in hThCAs at 14 dpf. Scale bars represent 200 µm. (F) Three-dimensional immunostaining for mCherry (left) and quantification of the location of mCherry+ axons within the hCOs of hThCAs at 14 dpf (right). Scale bar represents 500 µm. Data are expressed as mean ± SEM: n = 5 organoids. \*\*\**p* < 0.001, Welch’s *t*-test. (G) Schematic of each region and quantification of the proportion of mCherry+ thalamic axons in each region. Data are expressed as ± SEM: n = 8 hThCAs. \*\*\**p* < 0.001, Dunnett’s test. (H) Representative image of hThCAs at 14 dpf stained for EYFP and mCherry. Scale bars represent 500 µm. (I) Generation of human corticocortical assembloids (hCCAs) from EYFP-expressing and unlabeled hCOs. (J) EYFP-expressing axonal projections from hCOs to the other side within hThCAs and hCCAs at 21 dpf. Scale bars represent 500 µm. (K) Quantification of EYFP+ migrating cells from hCOs to the other side within hThCAs or hCCAs. Data are expressed as ± SEM: n = 8 hThCAs, n = 9 hCCAs. \**p* < 0.05, Welch’s *t*-test.

**Figure S3.**
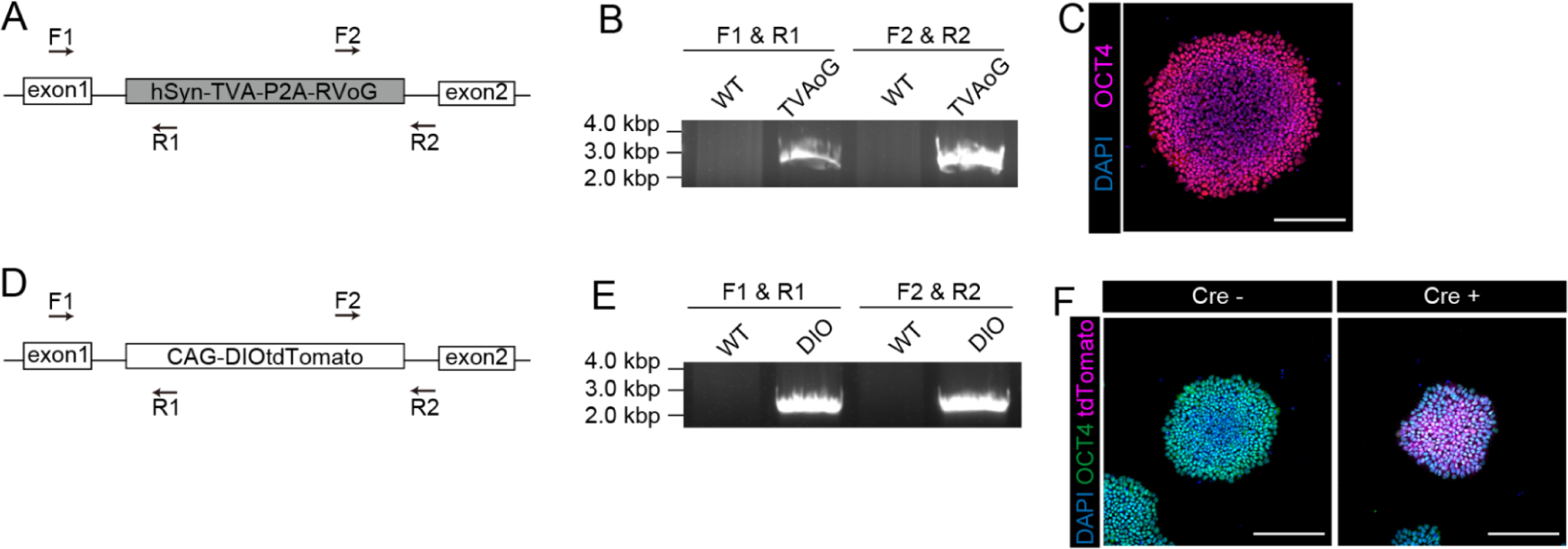
Generation of hiPSCs-TVA-oG and hiPSCs-DIO-tdTomato knocked-in lines. (A) Primers for genotyping the hiPSCs-TVA-P2A-oG line. Arrows, PCR primers. (B) Electrophoresis images of the PCR products for the hiPSCs-TVA-P2A-oG line. (C) Immunostaining for the pluripotent marker OCT4 in the hiPSCs-TVA-P2A-oG line. Scale bars represent 200 µm. (D) Primers for genotyping the hiPSCs-DIO-tdTomato line. Arrows, PCR primers. (E) Electrophoresis images of the PCR products for the knocked-in hiPSC lines. (F) Immunostaining for the pluripotent marker OCT4 and tdTomato in the hiPSCs-DIO- tdTomato line with or without Cre. Scale bars represent 200 µm.

**Figure S4.**
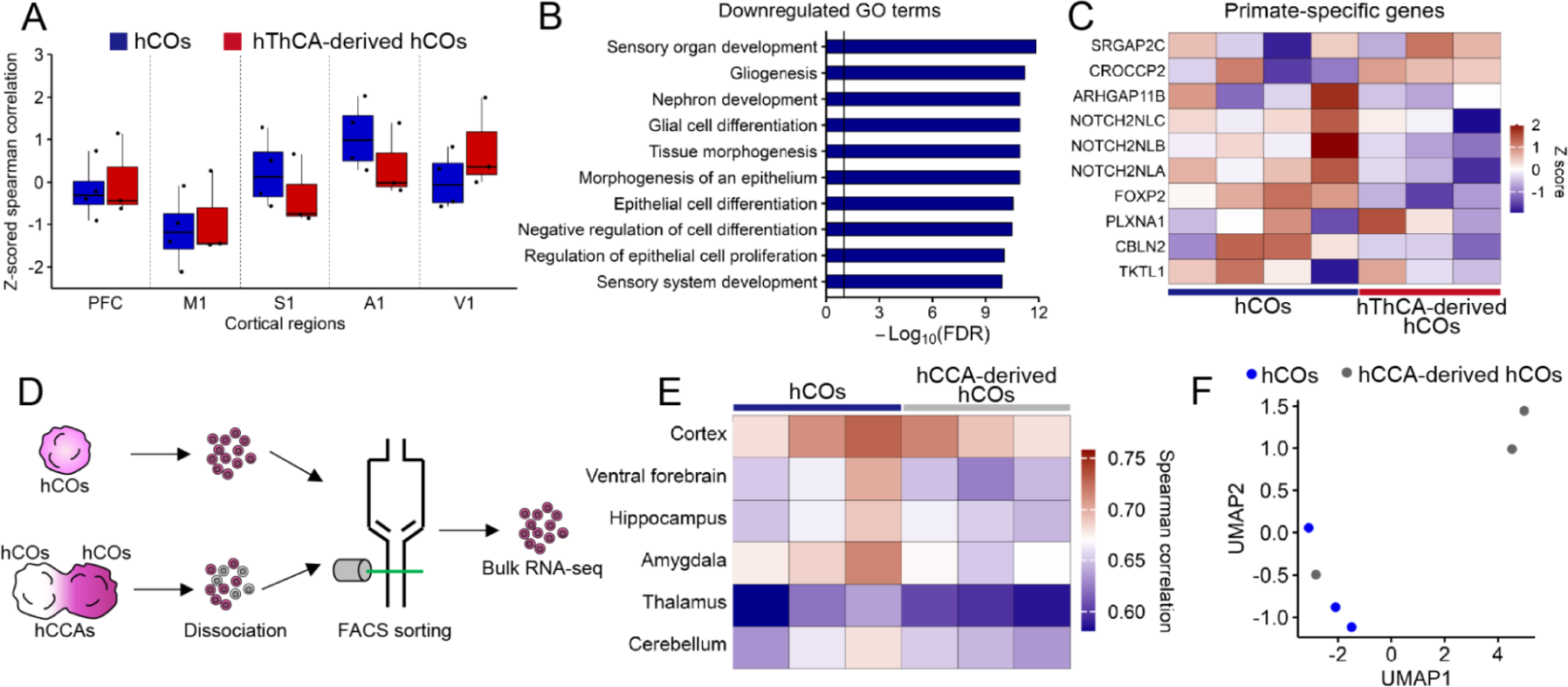
Characterization of transcriptomes in hCOs and assembloid-derived hCOs. (A) Z-scored spearman correlation between the transcriptomes of each region of human fetal cortex (12–17 pcw) and those of hCOs or hThCA-derived hCOs. n= 4 hCOs, n = 3 hThCA-derived hCOs. (B) Top 10 gene ontology terms downregulated in the hThCA-derived hCOs. Vertical line represents −log10FDR = 1. (C) Heatmap showing the expression of primate-specific genes in the hCOs and hThCA- derived hCOs. n= 4 hCOs, n = 3 hThCA-derived hCOs. (D) Bulk RNA sequencing method for hCOs and human corticocortical assembloids (hCCA)- derived hCOs. (E) Heatmap showing the Spearman correlations between the transcriptomes of region-specific human fetal brains and those of hCOs or hCCA-derived hCOs. n= 3 hCOs, n = 3 hCCA-derived hCOs. (F) UMAP for the transcriptomes of hCOs and hCCA-derived hCOs. n= 3 hCOs, n = 3 hCCA- derived hCOs.

**Figure S5.**
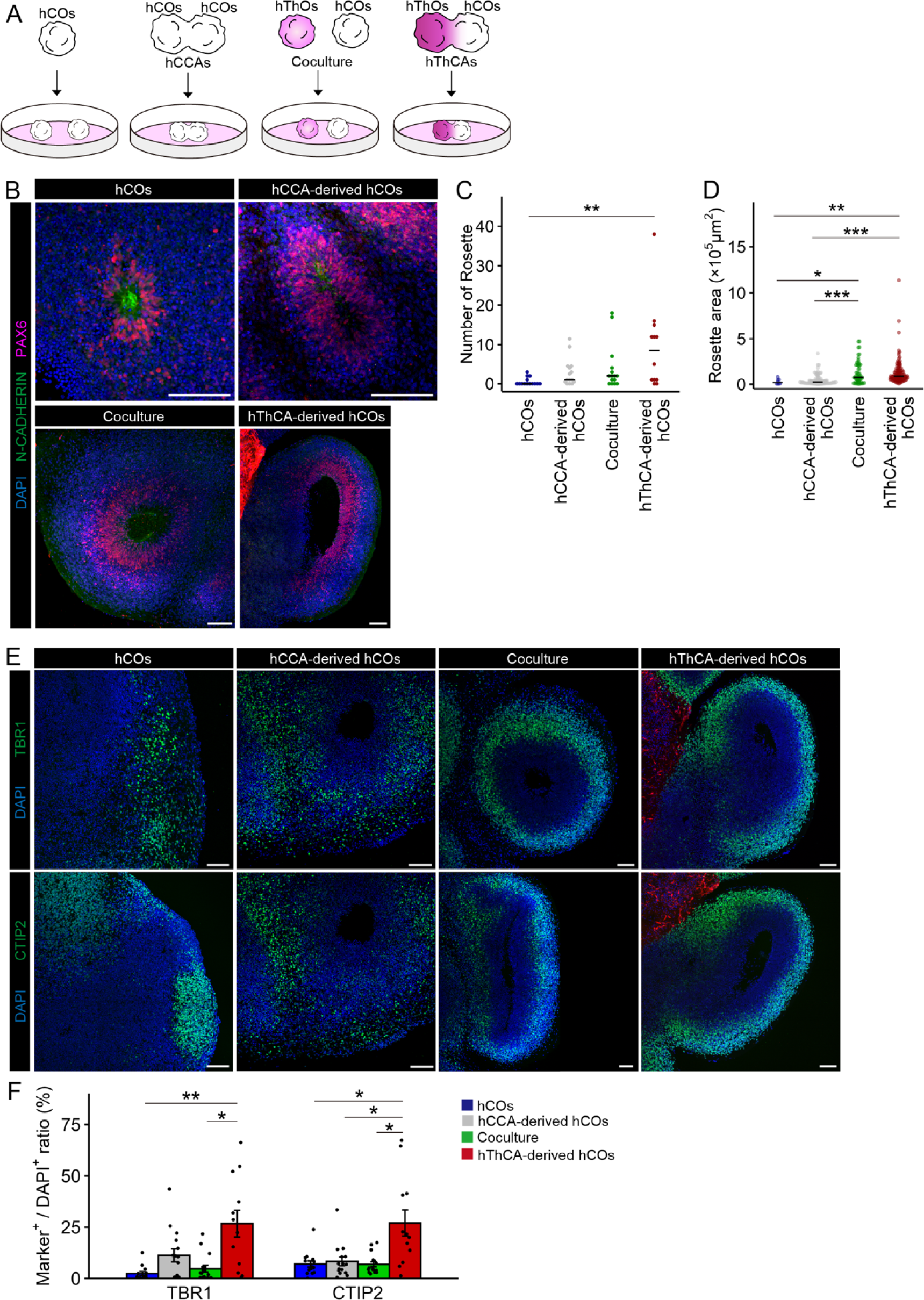
Cytoarchitecture in isolated hCOs, hCCA-derived hCOs, cocultured hCOs, and hThCA- derived hCOs. (A) Schematic of each culture condition. (B) Immunostaining for N-CADHERIN and PAX6 in each hCOs on day 80. Scale bars represent 100 µm. (C) Quantification of the number of rosettes in each hCOs. n = 14 hCOs, n = 15 hCCA-derived hCOs, n = 15 Coculture, and n = 12 hThCAs-derived hCOs. \**p* < 0.05, Dunn test. (D) Quantification of the area of each rosette in each hCOs. n = 8 rosettes (hCOs), n = 87 rosettes (hCCA-derived hCOs), n = 59 rosettes (Coculture), and n = 113 rosettes (hThCAs-derived hCOs). \**p* < 0.05, \*\**p* < 0.01, \*\*\**p* < 0.001, Dunn test. (E) Immunostaining for TBR1 and CTIP2 in each hCOs on day 80. Scale bars represent 100 µm. (F) Quantification of TBR1 and CTIP2 expression in each hCOs. Data are expressed as mean ± SEM. n = 14 hCOs, n = 15 hCCA-derived hCOs, n = 16 Coculture, and n = 12 hThCAs- derived hCOs. \**p* < 0.05, \*\**p* < 0.01, Dunn test.

**Figure S6.**
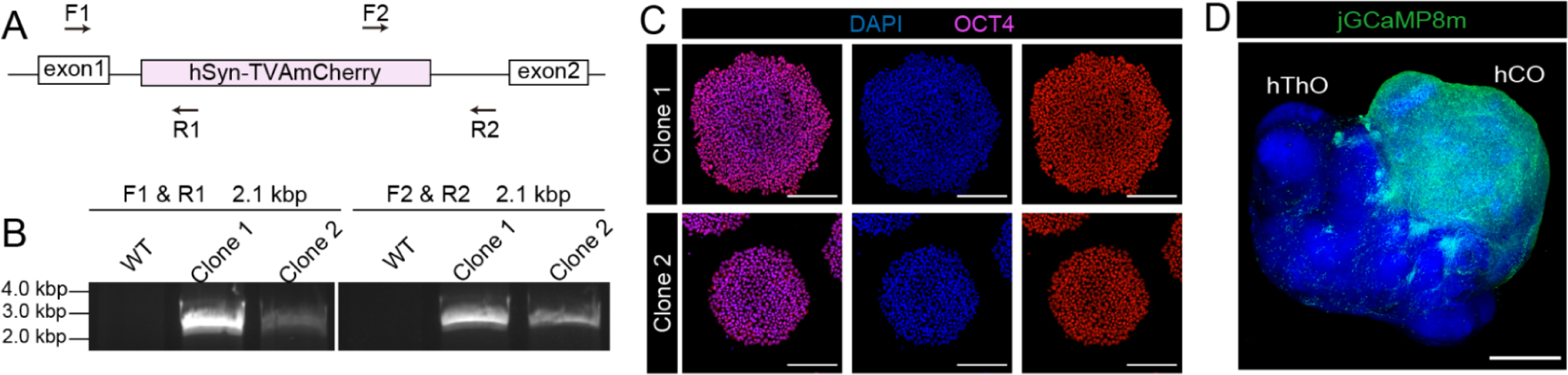
Generation of the hiPSCs-TVAmCherry line. (A) Primers for genotyping the hiPSCs-TVAmCherry line. Arrows, PCR primers. (B) Electrophoresis images of the PCR products for the knocked-in hiPSC lines. (C) Immunostaining for the pluripotent marker OCT4 in the hiPSCs-TVAmCherry line. Scale bars represent 200 µm. (D) Restricted jGCaMP8m expression in cortical neurons of the hThCAs on day 54. Scale bar represents 1,000 µm.

**Figure S7.**
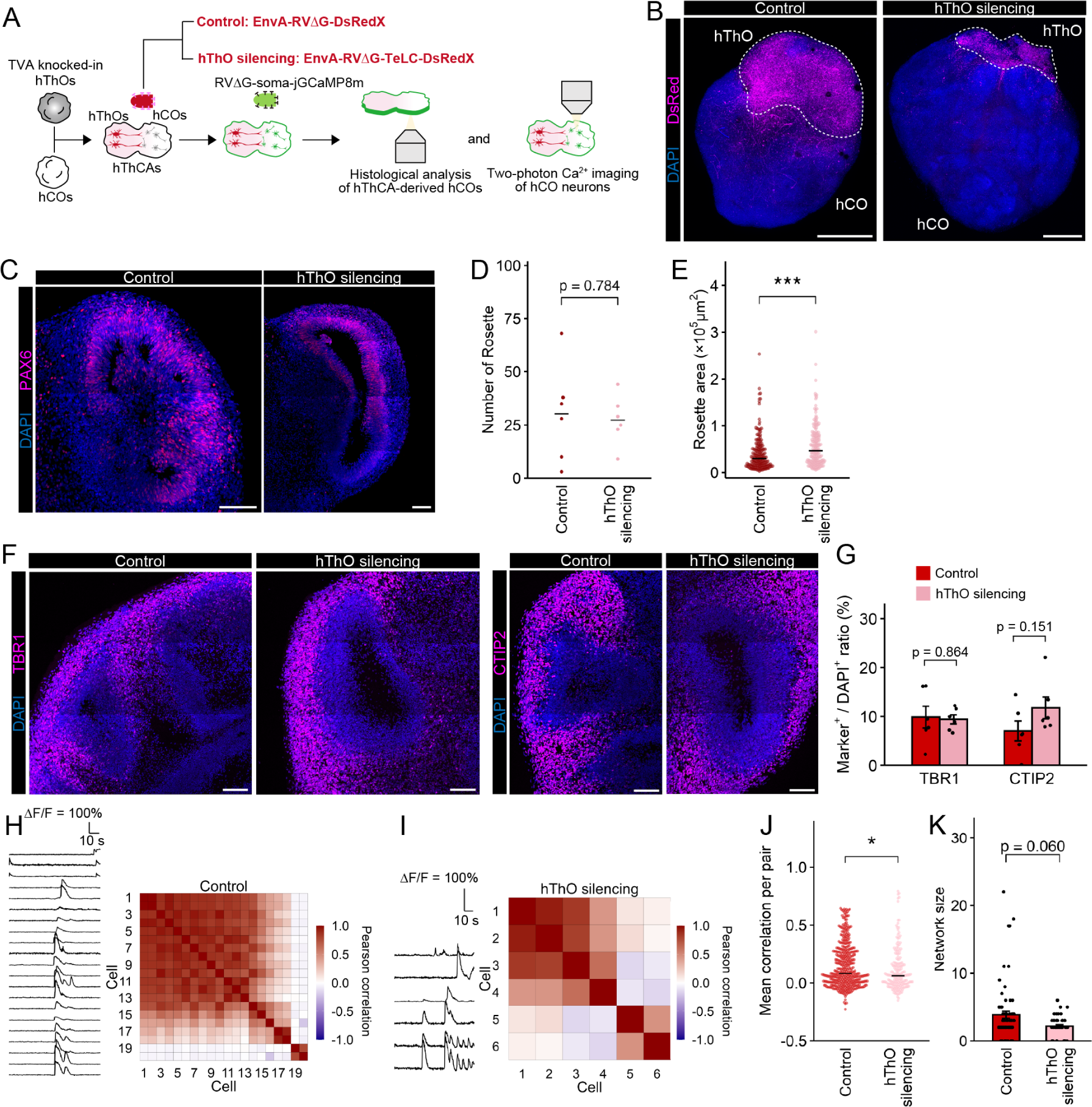
Silencing of hThCA-derived hThOs activity. (A) Experimental design for silencing of hThCA-derived hThOs activity. (B) Restricted DsRed expression in thalamic neurons of the hThCAs on day 101. Scale bars represent 1,000 µm. (C) Immunostaining for PAX6 in hThCAs in control and hThO silencing conditions on day 83. Scale bars represent 100 µm. (D) Quantification of the number of rosettes in hThCA-derived hCOs in each condition. n = 6 hThCA-derived hCOs (Control and hThO silencing). Welch’s *t*-test. (E) Quantification of the area of each rosette in hThCA-derived hCOs in each condition. n = 183 rosettes (Control), n = 164 rosettes (hThO silencing). \*\*\**p* < 0.001, Mann–Whitney U test. (F) Immunostaining for TBR1 and CTIP2 in hThCAs in each condition on day 83. Scale bars represent 100 µm. (G) Quantification of TBR1 and CTIP2 expression hThCA-derived hCOs in each condition. Data are expressed as mean ± SEM. n = 6 hThCA-derived hCOs (Control and hThO silencing). Welch’s *t*-test. (H) Representative time-series traces of jGCaMP8m signal changes (left) and heatmap showing correlation coefficients of activity between all pairs of simultaneously recorded neurons (right) in hThCA-derived hCOs in the control condition. (I) Representative time-series traces of jGCaMP8m signal changes (left) and heatmap showing correlation coefficients of activity between all pairs of simultaneously recorded neurons (right) in hThCA-derived hCOs in the hThO silencing condition. (J) Quantification of mean correlation coefficients of activity between all pairs of simultaneously recorded neurons from hThCA-derived hCOs in each condition. Red dots, correlation of each cell pair; black line, median. n = 389 pairs (Control), n = 179 pairs (hThO silencing), \**p* < 0.05, Mann–Whitney U test. (K) Quantification of the network synchronization size in hThCA-derived hCOs. Mann– Whitney U test. Data are expressed as mean ± SEM. n = 69 network (Control), n = 41 network (hThO silencing). Mann–Whitney U test.

**Figure S8.**
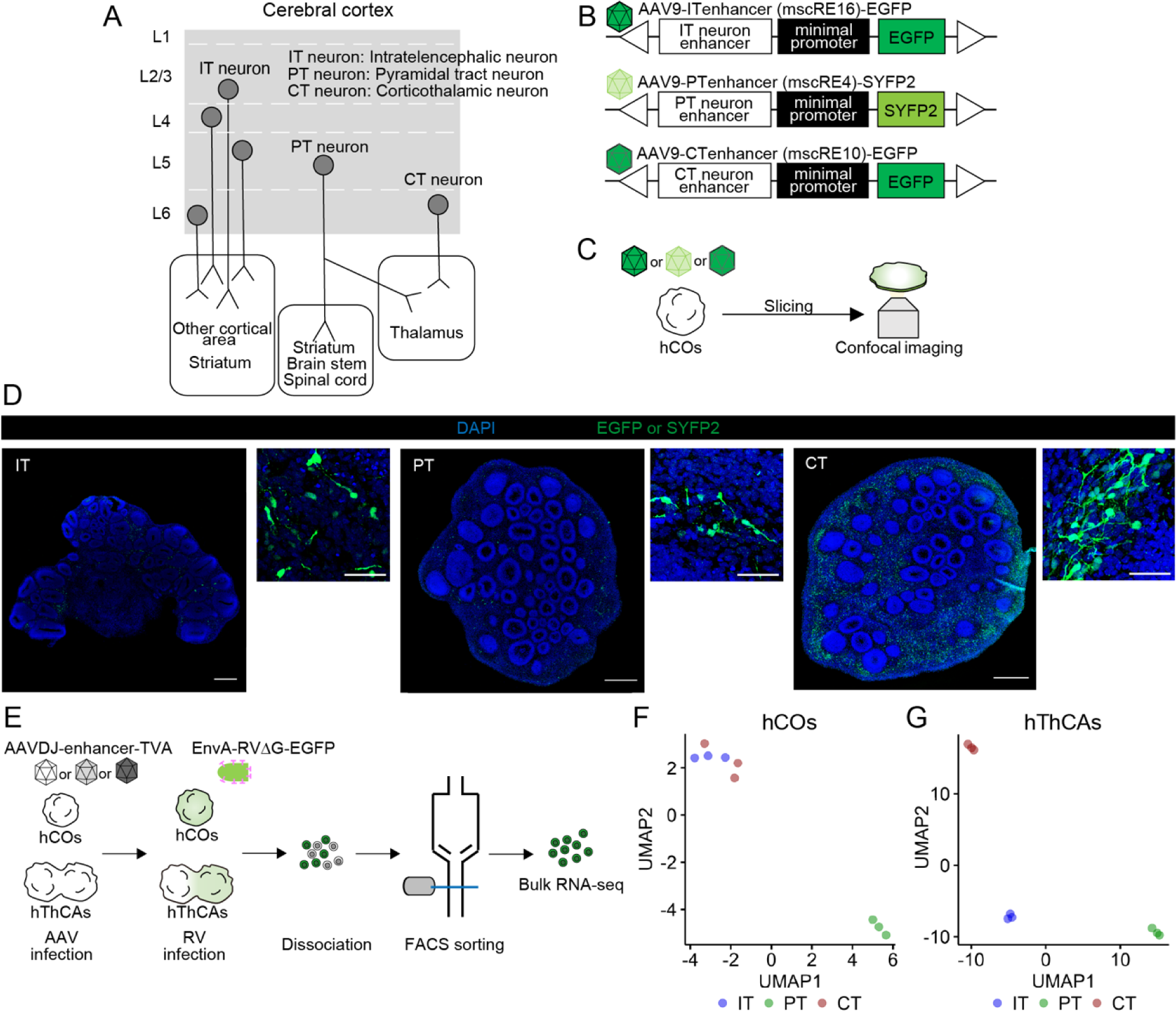
Validation of excitatory projection neuron subtypes in isolated hCOs and hThCA-derived hCOs. (A) Excitatory projection neuron subtypes in the cerebral cortex. (B) AAVs that encode a fluorescence protein under the cell type-specific enhancers mscRE16 (IT neurons: intratelencephalic neurons), mscRE4 (PT neurons: pyramidal tract neurons), and mscRE10 (CT neurons: corticothalamic neurons). (C) Schematic representation of viral labeling and confocal imaging in hCOs. (D) Immunostaining for GFP or SYFP2 in hCOs on day 52. hCOs were infected with AAV9- ITenhancer-EGFP (left), AAV9-PTenhancer-SYFP2 (middle), and AAV9-CTenhancer- EGFP (right). Scale bars represent 500 µm (left), 100 µm (right). (E) Bulk RNA sequencing method for each cortical subtype in isolated hCOs and hThCA- derived hCOs. (F) UMAP for the transcriptomes of each cortical subtype in isolated hCOs. n = 3 hCOs (IT, PT, and CT). (G) UMAP for the transcriptomes of each cortical subtype in hThCA-derived hCOs. n = 3 hThCA-derived hCOs (IT, PT, and CT).

**Figure S9.**
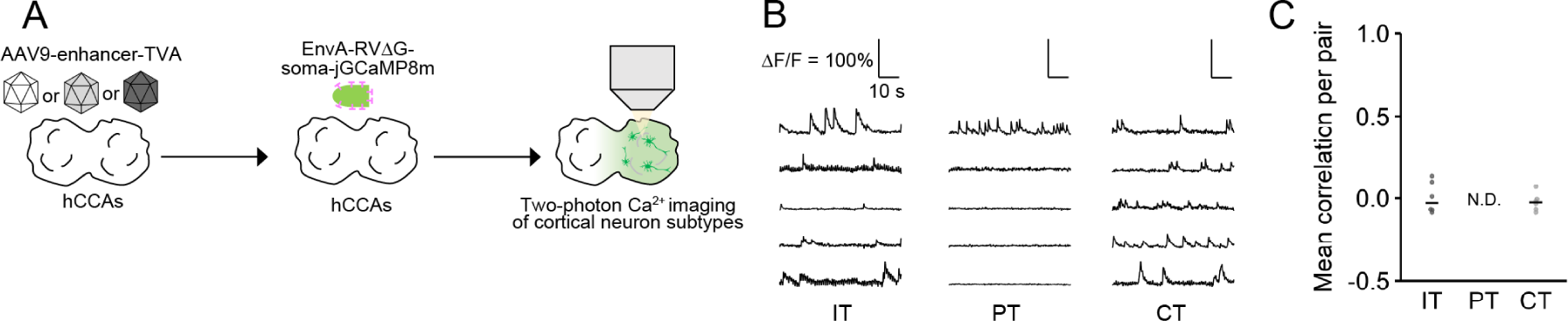
Cell type-specific two-photon Ca^2+^ imaging of hCCA-derived hCOs. (A) Method for cell type-specific two-photon Ca2+ imaging of hCCA-derived hCOs. (B) Representative time-series traces of jGCaMP8m signal changes of each cortical subtype in hCCA-derived hCOs. (C) Quantification of the mean correlation coefficients of the activity between all pairs of simultaneously recorded IT, PT, or CT neurons from hCCA-derived hCOs. n = 6 pairs (IT and CT).

**Table S1.**
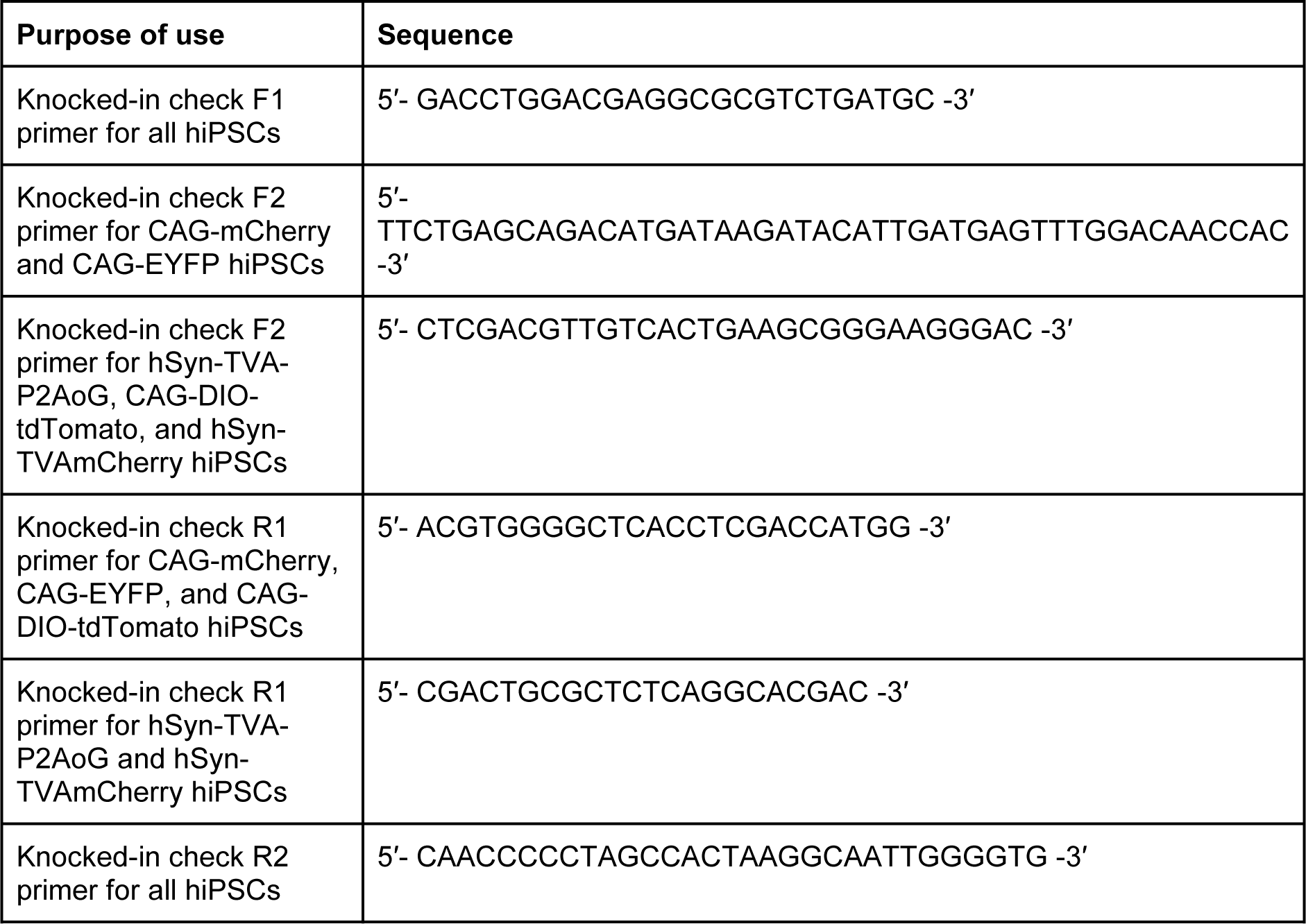
List of Knocked-in check primers.

**Table S2.**
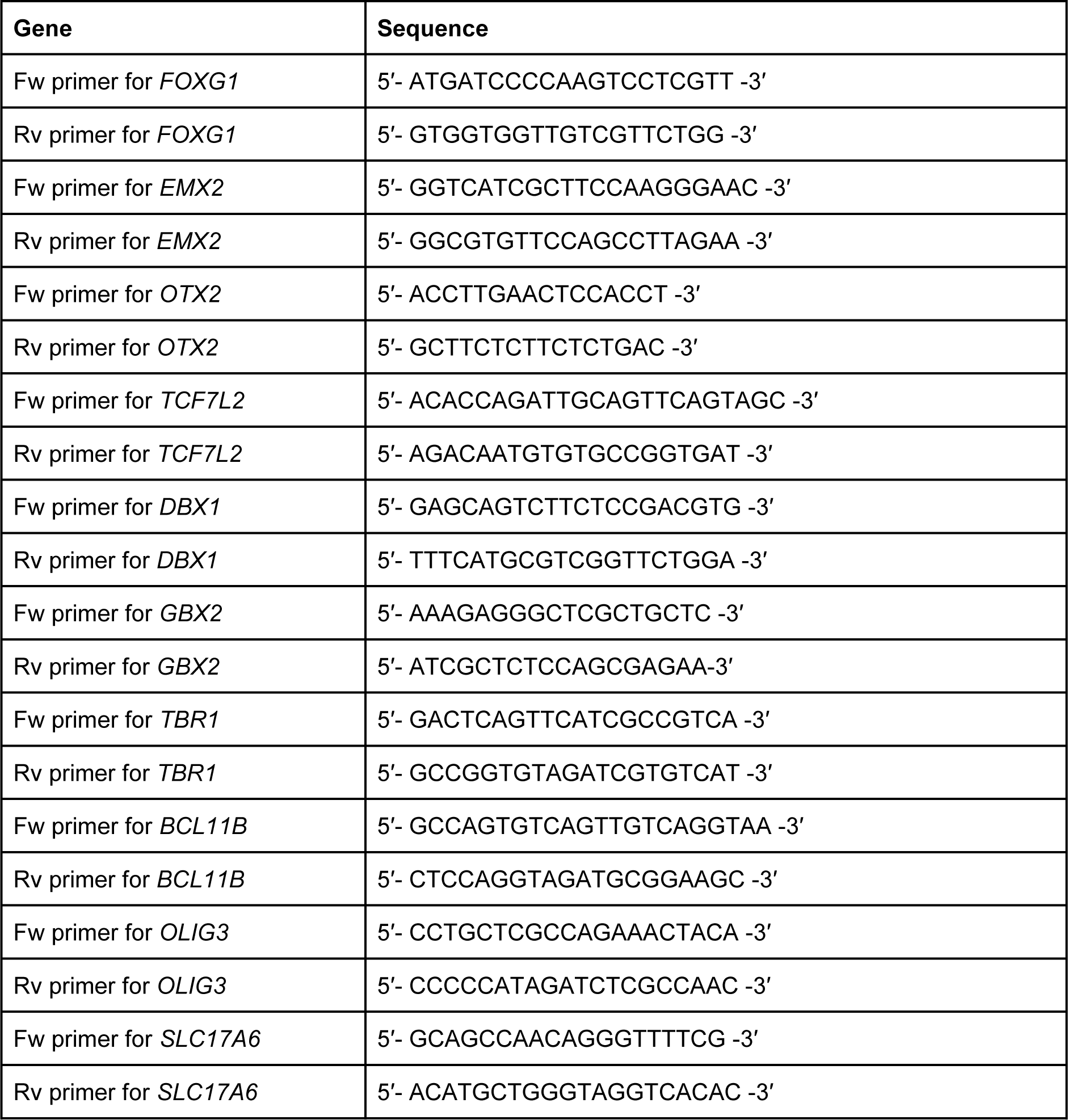
List of qPCR primers.

**Table S3.**
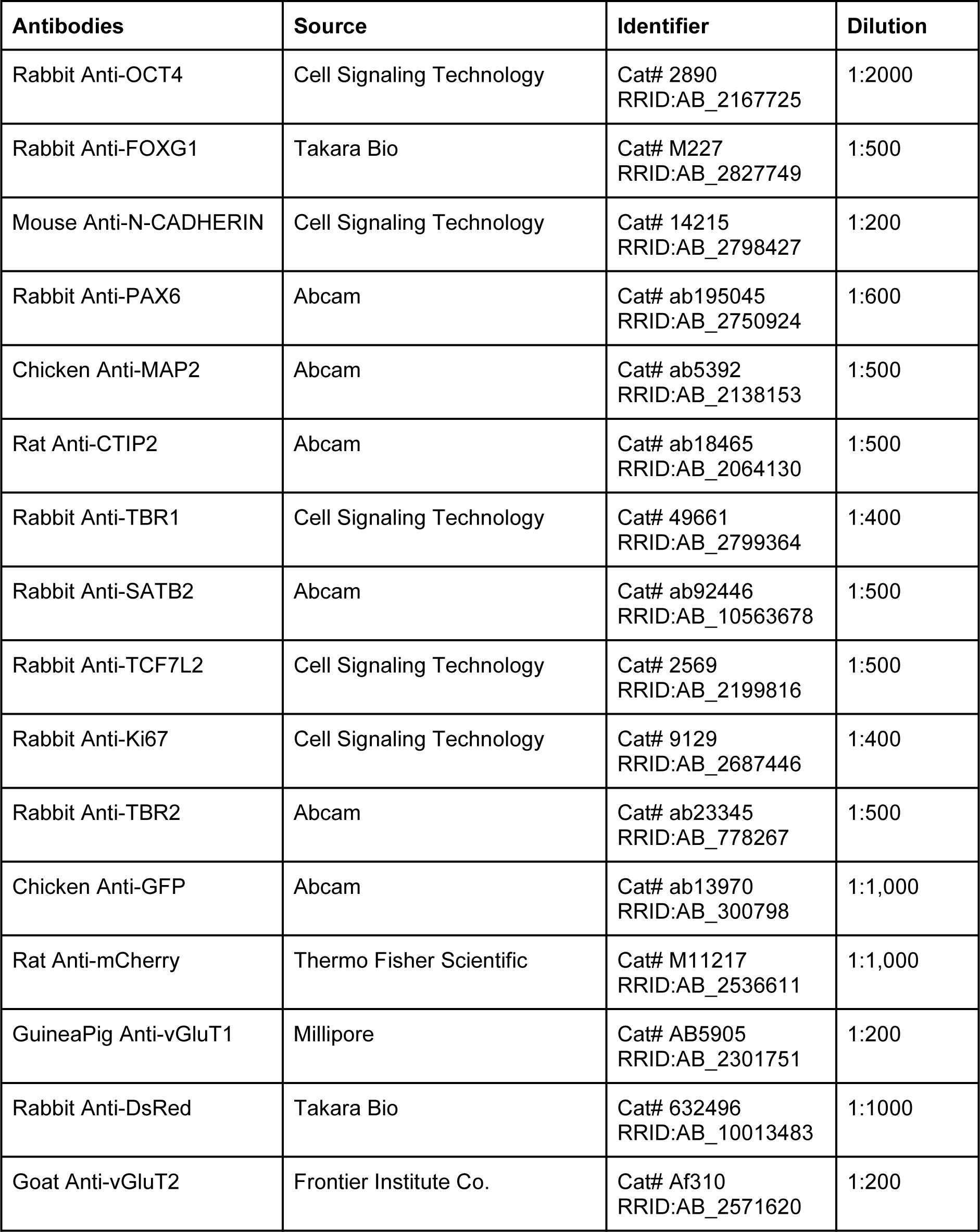
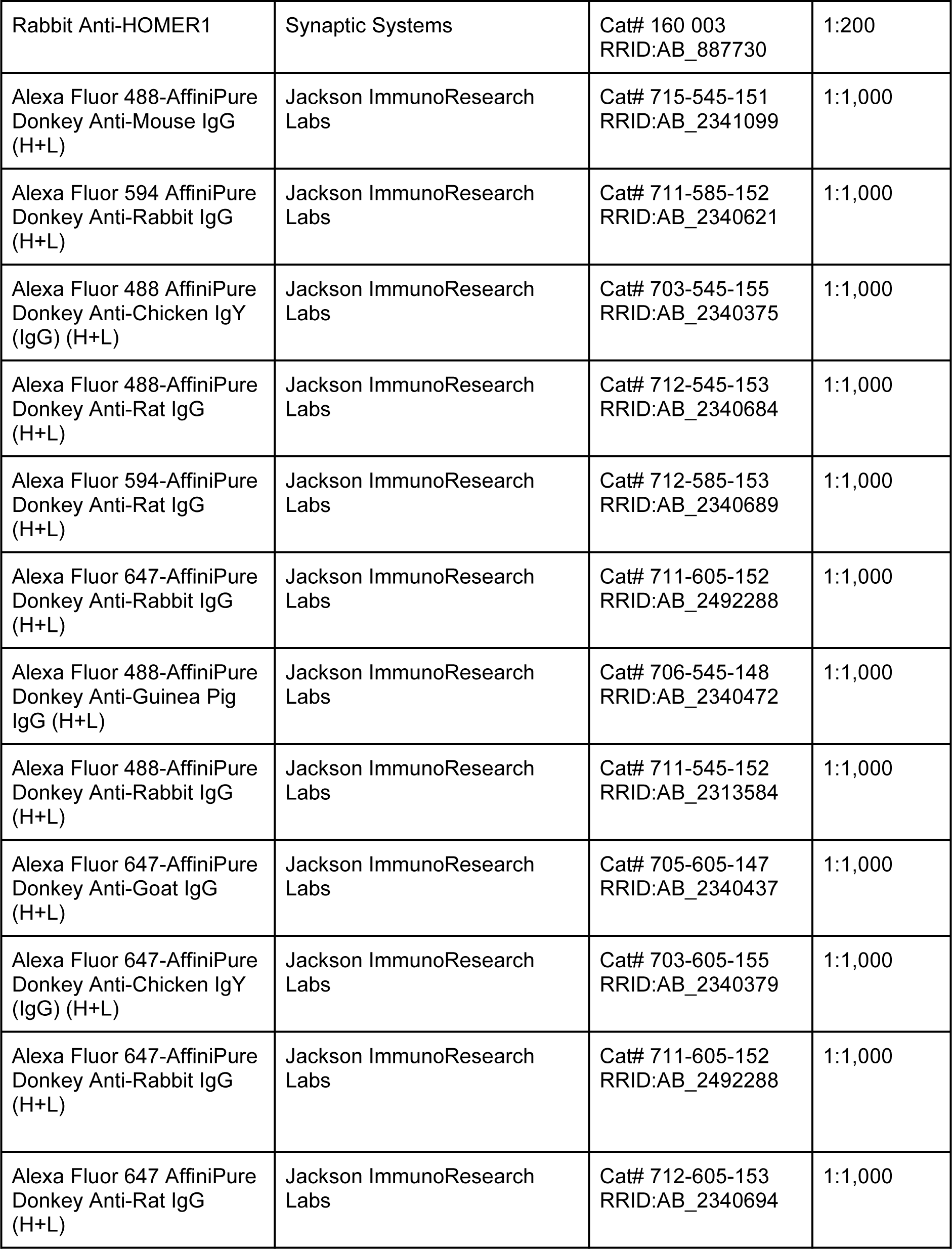
List of antibodies.

**Table S4.**
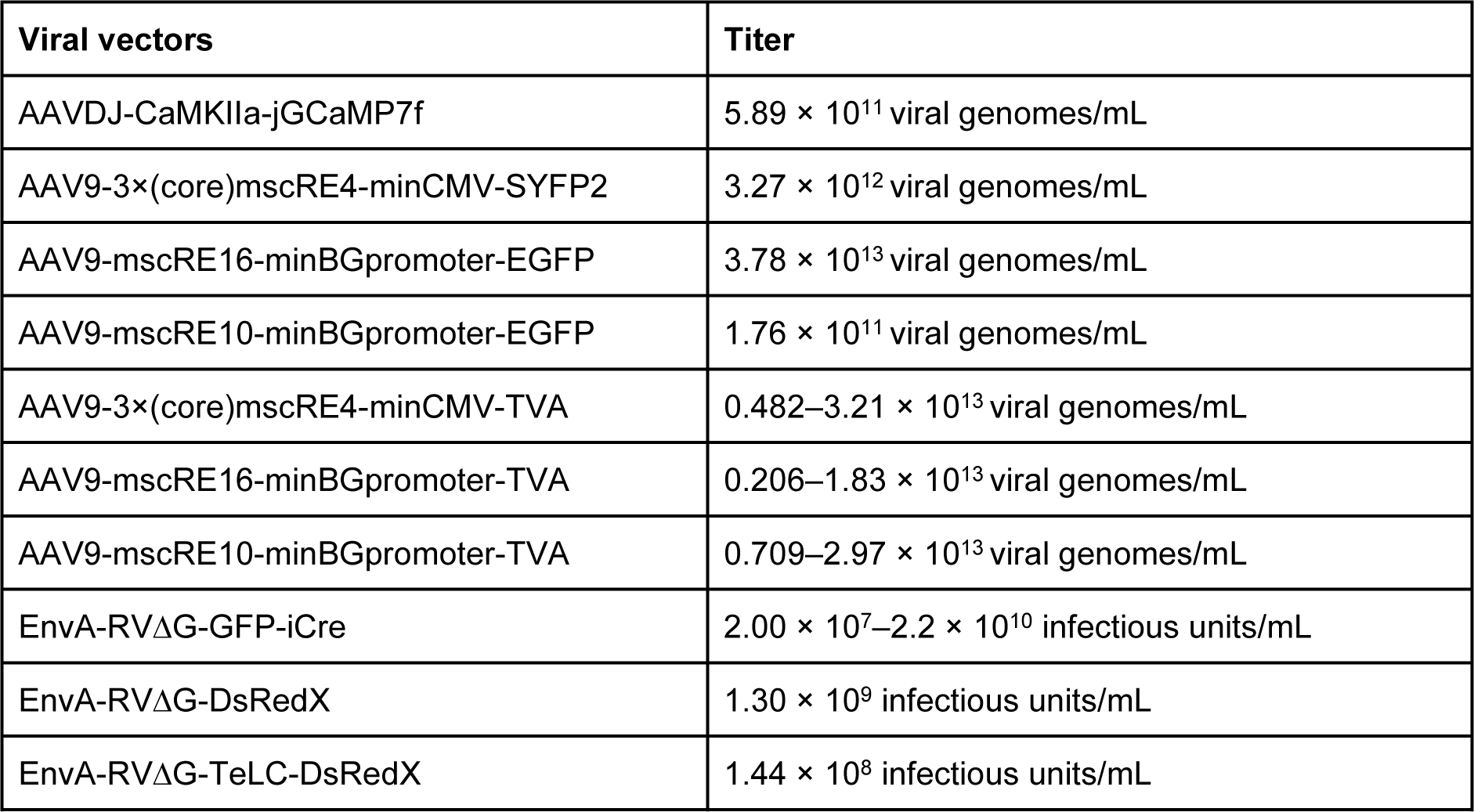
Titers of viral vectors.

| Viral vectors | Titer |
| --- | --- |
| AAVDJ-CaMKIIa-jGCaMP7f | $5.89 \times 10^{11}$ viral genomes/mL |
| AAV9-3×(core)mscRE4-minCMV-SYFP2 | $3.27 \times 10^{12}$ viral genomes/mL |
| AAV9-mscRE16-minBGpromoter-EGFP | $3.78 \times 10^{13}$ viral genomes/mL |
| AAV9-mscRE10-minBGpromoter-EGFP | $1.76 \times 10^{11}$ viral genomes/mL |
| AAV9-3×(core)mscRE4-minCMV-TVA | $0.482\text{--}3.21 \times 10^{13}$ viral genomes/mL |
| AAV9-mscRE16-minBGpromoter-TVA | $0.206\text{--}1.83 \times 10^{13}$ viral genomes/mL |
| AAV9-mscRE10-minBGpromoter-TVA | $0.709\text{--}2.97 \times 10^{13}$ viral genomes/mL |
| EnvA-RVΔG-GFP-iCre | $2.00 \times 10^7\text{--}2.2 \times 10^{10}$ infectious units/mL |
| EnvA-RVΔG-DsRedX | $1.30 \times 10^9$ infectious units/mL |
| EnvA-RVΔG-TeLC-DsRedX | $1.44 \times 10^8$ infectious units/mL |

**Table S5.**
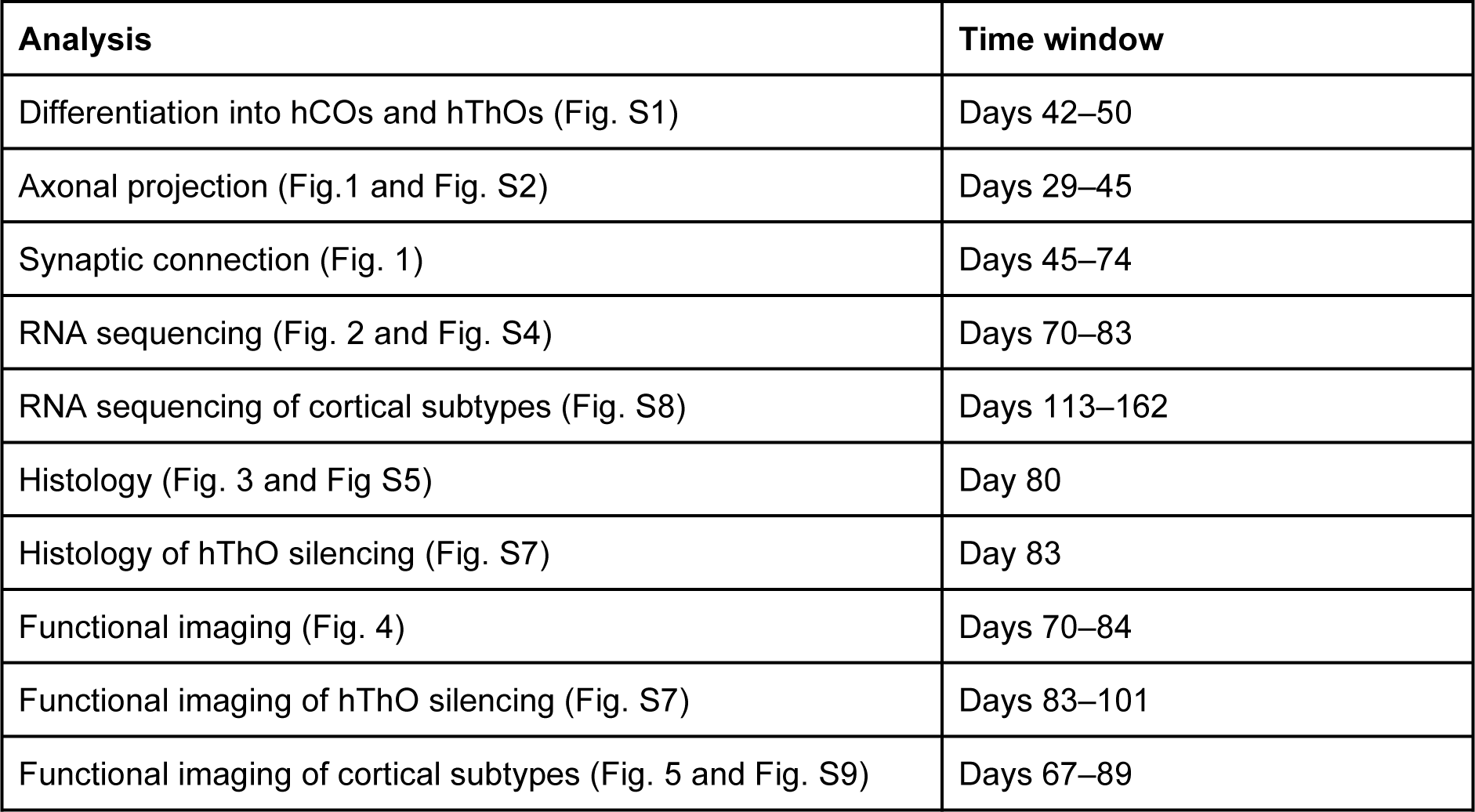
The time window of analysis.

## Notes

**Competing Interest Statement:** The authors have no competing interests to declare.

### Competing Interest Statement

The authors have declared no competing interest.

## References

1. D. C. Van Essen, et al., Cerebral cortical folding, parcellation, and connectivity in humans, nonhuman primates, and mice. Proc. Natl. Acad. Sci. U. S. A. 116, 26173–26180 (2019).

2. S. M. Kolk, P. Rakic, Development of prefrontal cortex. Neuropsychopharmacology 47, 41–57 (2022).

3. G. M. G. Shepherd, Corticostriatal connectivity and its role in disease. Nat. Rev. Neurosci. 14, 278– 291 (2013).

4. A. Baker, et al., Specialized Subpopulations of Deep-Layer Pyramidal Neurons in the Neocortex: Bridging Cellular Properties to Functional Consequences. J. Neurosci. 38, 5441–5455 (2018).

5. G. M. G. Shepherd, N. Yamawaki, Untangling the cortico-thalamo-cortical loop: cellular pieces of a knotty circuit puzzle. Nat. Rev. Neurosci. 22, 389–406 (2021).

6. S. M. Sherman, Thalamus plays a central role in ongoing cortical functioning. Nat. Neurosci. 19, 533–541 (2016).

7. M. M. Halassa, S. M. Sherman, Thalamocortical circuit motifs: A general framework. Neuron 103, 762–770 (2019).

8. A. Alzu’bi, J. Homman-Ludiye, J. A. Bourne, G. J. Clowry, Thalamocortical Afferents Innervate the Cortical Subplate much Earlier in Development in Primate than in Rodent. Cereb. Cortex 29, 1706– 1718 (2019).

9. G. López-Bendito, Z. Molnár, Thalamocortical development: how are we going to get there? Nat. Rev. Neurosci. 4, 276–289 (2003).

10. S.-J. Chou, et al., Geniculocortical input drives genetic distinctions between primary and higher-order visual areas. Science 340, 1239–1242 (2013).

11. G. Pouchelon, et al., Modality-specific thalamocortical inputs instruct the identity of postsynaptic L4 neurons. Nature 511, 471–474 (2014).

12. V. Moreno-Juan, et al., Prenatal thalamic waves regulate cortical area size prior to sensory processing. Nat. Commun. 8, 14172 (2017).

13. N. Antón-Bolaños, et al., Prenatal activity from thalamic neurons governs the emergence of functional cortical maps in mice. Science 364, 987–990 (2019).

14. Y. Tanaka, I.-H. Park, Regional specification and complementation with non-neuroectodermal cells in human brain organoids. J. Mol. Med. 99, 489–500 (2021).

15. S. P. Pașca, et al., A framework for neural organoids, assembloids and transplantation studies. Nature 639, 315–320 (2025).

16. S. P. Pașca, The rise of three-dimensional human brain cultures. Nature 553, 437–445 (2018).

17. M. A. Lancaster, J. A. Knoblich, Organogenesis in a dish: modeling development and disease using organoid technologies. Science 345, 1247125 (2014).

18. M. A. Lancaster, et al., Cerebral organoids model human brain development and microcephaly. Nature 501, 373–379 (2013).

19. M. Eiraku, et al., Self-organized formation of polarized cortical tissues from ESCs and its active manipulation by extrinsic signals. Cell Stem Cell 3, 519–532 (2008).

20. J. Mariani, et al., Modeling human cortical development in vitro using induced pluripotent stem cells. Proc. Natl. Acad. Sci. U. S. A. 109, 12770–12775 (2012).

21. T. Kadoshima, et al., Self-organization of axial polarity, inside-out layer pattern, and species-specific progenitor dynamics in human ES cell-derived neocortex. Proc. Natl. Acad. Sci. U. S. A. 110, 20284– 20289 (2013).

22. A. M. Paşca, et al., Functional cortical neurons and astrocytes from human pluripotent stem cells in 3D culture. Nat. Methods 12, 671–678 (2015).

23. M. Nishimura, et al., Conversion of silent synapses to AMPA receptor-mediated functional synapses in human cortical organoids. Neurosci. Res. 212, 20–30 (2025).

24. F. Birey, et al., Assembly of functionally integrated human forebrain spheroids. Nature 545, 54–59 (2017).

25. J. A. Bagley, D. Reumann, S. Bian, J. Lévi-Strauss, J. A. Knoblich, Fused cerebral organoids model interactions between brain regions. Nat. Methods 14, 743–751 (2017).

26. Y. Xiang, et al., Fusion of Regionally Specified hPSC-Derived Organoids Models Human Brain Development and Interneuron Migration. Cell Stem Cell 21, 383–398.e7 (2017).

27. Y. Miura, et al., Generation of human striatal organoids and cortico-striatal assembloids from human pluripotent stem cells. Nat. Biotechnol. 38, 1421–1430 (2020).

28. Y. Xiang, et al., hESC-Derived Thalamic Organoids Form Reciprocal Projections When Fused with Cortical Organoids. Cell Stem Cell 24, 487–497.e7 (2019).

29. J.-I. Kim, et al., Human assembloids reveal the consequences of CACNA1G gene variants in the thalamocortical pathway. Neuron 112, 4048–4059.e7 (2024).

30. M. H. Patton, et al., Synaptic plasticity in human thalamocortical assembloids. Cell Rep. 43, 114503 (2024).

31. H. Dana, et al., High-performance calcium sensors for imaging activity in neuronal populations and microcompartments. Nat. Methods 16, 649–657 (2019).

32. E. A. Susaki, et al., Whole-brain imaging with single-cell resolution using chemical cocktails and computational analysis. Cell 157, 726–739 (2014).

33. H. J. Kang, et al., Spatio-temporal transcriptome of the human brain. Nature 478, 483–489 (2011).

34. I. Kostović, M. Judas, The development of the subplate and thalamocortical connections in the human foetal brain: Human foetal cortical circuitry. Acta Paediatr. 99, 1119–1127 (2010).

35. P. Vanderhaeghen, F. Polleux, Developmental mechanisms underlying the evolution of human cortical circuits. Nat. Rev. Neurosci. 24, 213–232 (2023).

36. B. Ataman, et al., Evolution of Osteocrin as an activity-regulated factor in the primate brain. Nature 539, 242–247 (2016).

37. C. Dehay, P. Savatier, V. Cortay, H. Kennedy, Cell-cycle kinetics of neocortical precursors are influenced by embryonic thalamic axons. J. Neurosci. 21, 201–214 (2001).

38. F. Osakada, E. M. Callaway, Design and generation of recombinant rabies virus vectors. Nat. Protoc. 8, 1583–1601 (2013).

39. Y. Zhang, et al., Fast and sensitive GCaMP calcium indicators for imaging neural populations. bioRxiv 2021.11.08.467793 (2021).

40. F. Osakada, et al., New rabies virus variants for monitoring and manipulating activity and gene expression in defined neural circuits. Neuron 71, 617–631 (2011).

41. E. Link, et al., Tetanus toxin action: inhibition of neurotransmitter release linked to synaptobrevin proteolysis. Biochem. Biophys. Res. Commun. 189, 1017–1023 (1992).

42. M. Yamamoto, et al., Reversible suppression of glutamatergic neurotransmission of cerebellar granule cells in vivo by genetically manipulated expression of tetanus neurotoxin light chain. J. Neurosci. 23, 6759–6767 (2003).

43. L. T. Graybuck, et al., Enhancer viruses for combinatorial cell-subclass-specific labeling. Neuron 109, 1449–1464.e13 (2021).

44. N. Antón-Bolaños, A. Espinosa, G. López-Bendito, Developmental interactions between thalamus and cortex: a true love reciprocal story. Curr. Opin. Neurobiol. 52, 33–41 (2018).

45. Z. Molnár, S. Garel, G. López-Bendito, P. Maness, D. J. Price, Mechanisms controlling the guidance of thalamocortical axons through the embryonic forebrain. Eur. J. Neurosci. 35, 1573–1585 (2012).

46. G. López-Bendito, et al., Tangential neuronal migration controls axon guidance: a role for neuregulin-1 in thalamocortical axon navigation. Cell 125, 127–142 (2006).

47. Y. Shinmyo, et al., Draxin from neocortical neurons controls the guidance of thalamocortical projections into the neocortex. Nat. Commun. 6, 10232 (2015).

48. D. D. M. O’Leary, S.-J. Chou, S. Sahara, Area patterning of the mammalian cortex. Neuron 56, 252– 269 (2007).

49. C. R. Cadwell, A. Bhaduri, M. A. Mostajo-Radji, M. G. Keefe, T. J. Nowakowski, Development and Arealization of the Cerebral Cortex. Neuron 103, 980–1004 (2019).

50. E. M. Miyashita-Lin, R. Hevner, K. M. Wassarman, S. Martinez, J. L. Rubenstein, Early neocortical regionalization in the absence of thalamic innervation. Science 285, 906–909 (1999).

51. A. Bhaduri, et al., Cell stress in cortical organoids impairs molecular subtype specification. Nature 578, 142–148 (2020).

52. H. Sato, et al., Thalamus-derived molecules promote survival and dendritic growth of developing cortical neurons. J. Neurosci. 32, 15388–15402 (2012).

53. R. Llinás, F. J. Urbano, E. Leznik, R. R. Ramírez, H. J. F. van Marle, Rhythmic and dysrhythmic thalamocortical dynamics: GABA systems and the edge effect. Trends Neurosci. 28, 325–333 (2005).

54. M. M. Angulo Salavarria, C. Dell’Amico, A. D’Agostino, L. Conti, M. Onorati, Cortico-thalamic development and disease: From cells, to circuits, to schizophrenia. Front. Neuroanat. 17, 1130797 (2023).

55. L. C. Greig, M. B. Woodworth, M. J. Galazo, H. Padmanabhan, J. D. Macklis, Molecular logic of neocortical projection neuron specification, development and diversity. Nat. Rev. Neurosci. 14, 755– 769 (2013).

56. N. D. Woodward, M. Giraldo-Chica, B. Rogers, C. J. Cascio, Thalamocortical Dysconnectivity in Autism Spectrum Disorder: An Analysis of the Autism Brain Imaging Data Exchange. Biological Psychiatry: Cognitive Neuroscience and Neuroimaging 2, 76–84 (2017).

57. D. S. Roy, et al., Anterior thalamic dysfunction underlies cognitive deficits in a subset of neuropsychiatric disease models. Neuron 109, 2590–2603.e13 (2021).

58. R. Ayub, et al., Thalamocortical connectivity is associated with autism symptoms in high-functioning adults with autism and typically developing adults. Transl. Psychiatry 11, 93 (2021).

59. K. Ye, et al., Reproducible production and image-based quality evaluation of retinal pigment epithelium sheets from human induced pluripotent stem cells. Sci. Rep. 10, 14387 (2020).

60. S. A. Sloan, J. Andersen, A. M. Pașca, F. Birey, S. P. Pașca, Generation and assembly of human brain region-specific three-dimensional cultures. Nat. Protoc. 13, 2062–2085 (2018).

61. T. Kodera, et al., Modeling the marmoset brain using embryonic stem cell-derived cerebral assembloids. Biochem. Biophys. Res. Commun. 657, 119–127 (2023).

62. F. Osakada, et al., Toward the generation of rod and cone photoreceptors from mouse, monkey and human embryonic stem cells. Nat. Biotechnol. 26, 215–224 (2008).

63. Y. Masaki, M. Yamaguchi, R. F. Takeuchi, F. Osakada, Monosynaptic rabies virus tracing from projection-targeted single neurons. Neurosci. Res. 178, 20–32 (2022).

64. M. Guizar-Sicairos, S. T. Thurman, J. R. Fienup, Efficient subpixel image registration algorithms. Opt. Lett. 33, 156–158 (2008).

65. B. Scholl, D. E. Wilson, D. Fitzpatrick, Local order within global disorder: Synaptic architecture of visual space. Neuron 96, 1127–1138.e4 (2017).

66. R. F. Takeuchi, et al., Posteromedial cortical networks encode visuomotor prediction errors. bioRxiv 2022.08.16.504075 (2022).

67. E. A. Susaki, et al., Advanced CUBIC protocols for whole-brain and whole-body clearing and imaging. Nat. Protoc. 10, 1709–1727 (2015).

68. T. Suzuki, N. Morimoto, A. Akaike, F. Osakada, Multiplex neural circuit tracing with G-deleted rabies viral vectors. Front. Neural Circuits 13, 77 (2019).

69. S. Okigawa, et al., Cell type- and layer-specific convergence in core and shell neurons of the dorsal lateral geniculate nucleus. J. Comp. Neurol. 529, 2099–2124 (2021).

70. S. Melliou, P. Diamandis, Generation of cell-type-specific proteomes of neurodevelopment from human cerebral organoids. STAR Protoc 3, 101774 (2022).

71. R. Patro, G. Duggal, M. I. Love, R. A. Irizarry, C. Kingsford, Salmon provides fast and bias-aware quantification of transcript expression. Nat. Methods 14, 417–419 (2017).

