## Supplementary material for "Thalamus–cortex interactions drive cell type-specific cortical development in human pluripotent stem cell-derived assembloids": Figs S1-S9 and Tables S1-S5

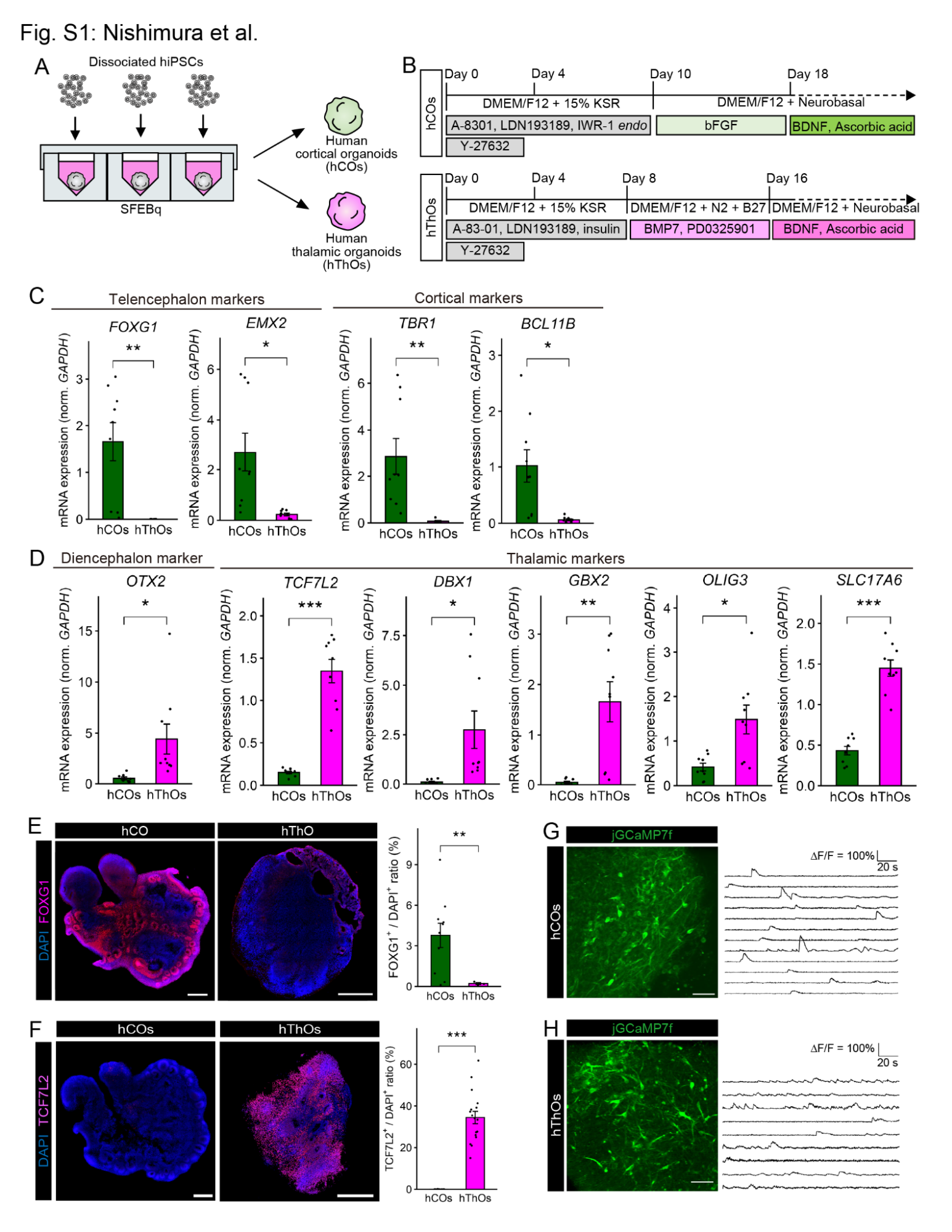

### Figure S1. Characterization of hCOs and hThOs.

(A) Schematic of generating region-specific organoids from hiPSCs.
(B) Protocol of the stepwise treatment for generating hCOs and hThOs from hiPSCs.
(C) qPCR analysis of telencephalic and cortical markers in hCOs and hThOs on day 42. Data are expressed as mean ± SEM. n = 9 hCOs, n = 9 hThOs, **p* < 0.05, ***p* < 0.01, ****p* < 0.001, Welch’s *t*-test.
(D) qPCR analysis of diencephalic and thalamic markers in hCOs and hThOs on day 42. Data are expressed as mean ± SEM. n = 9 hCOs and n = 9 hThOs, **p* < 0.05, ***p* < 0.01, ****p* < 0.001, Welch’s *t*-test.
(E) Immunostaining for FOXG1 (left) and quantification of FOXG1 expression in hCOs and hThOs (right) on day 42. Data are expressed as mean ± SEM: n = 13 hCOs, n = 3 hThOs. Scale bar represents 500 µm. ***p* < 0.01, Welch’s *t*-test.
(F) Immunostaining for TCF7L2 (left) and quantification of TCF7L2 expression in hCOs and hThOs (right) on day 42. Data are expressed as mean ± SEM: n = 11 hCOs, n = 17 hThOs. Scale bar represents 500 µm. ****p* < 0.001, Welch’s *t*-test.
(G) Representative images and time-series traces of jGCaMP7f signal changes in hCOs on days 50. Scale bar represents 50 µm.
(H) Representative images and time-series traces of jGCaMP7f signal changes in hThOs on days 50. Scale bar represents 50 µm.

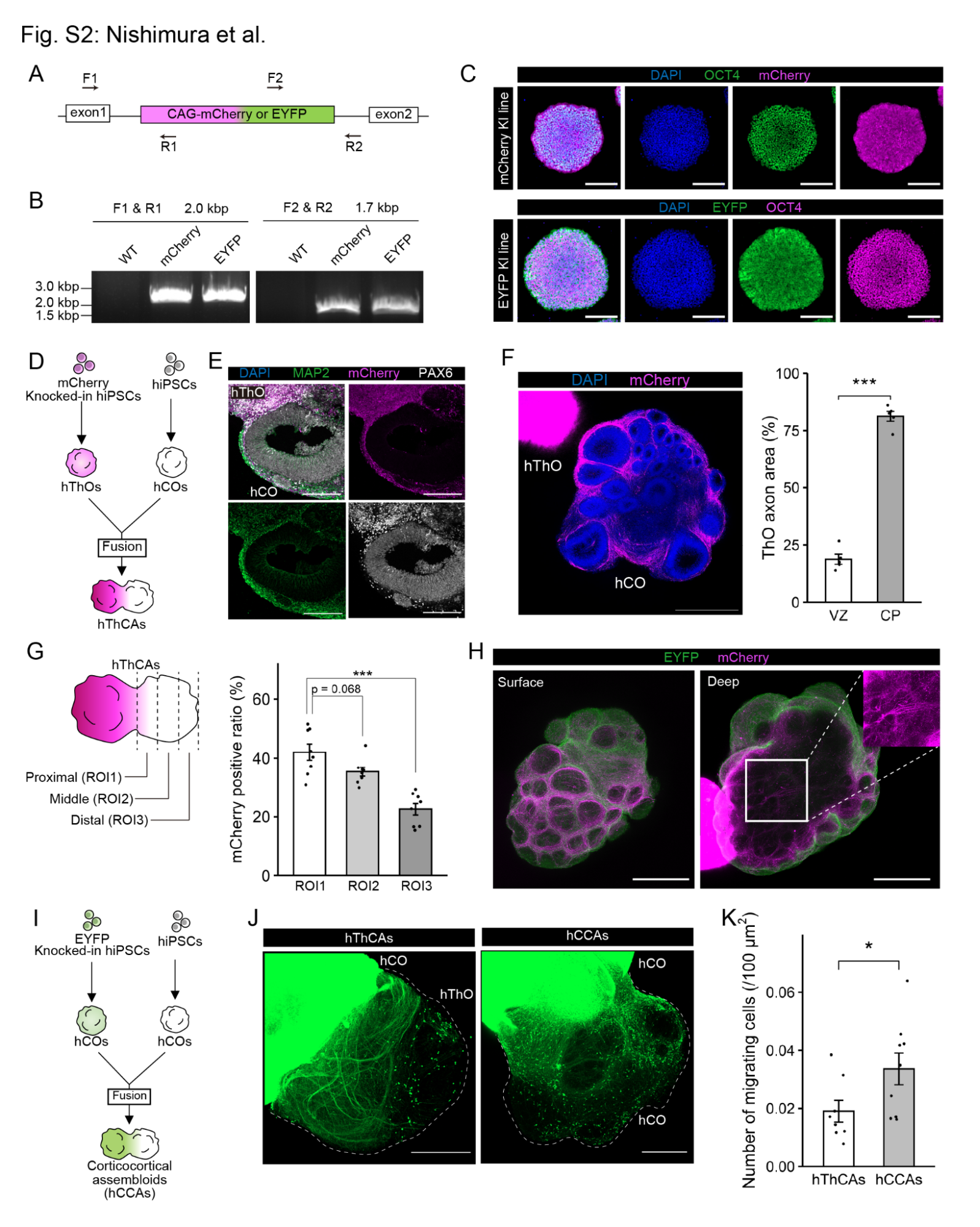

### Figure S2. Characterization of reciprocal projections within hThCAs.

(A) Primers for genotyping mCherry or EYFP knocked-in iPSC lines. Arrows, PCR primers.
(B) Electrophoresis images of the PCR products for mCherry-KI or EYFP-KI iPSC lines.
(C) Immunostaining for the pluripotent marker OCT4 and fluorescent proteins in mCherry-KI or EYFP-KI hiPSC lines. Scale bars represent 200 µm.
(D) Generation of hThCAs from mCherry-expressing hThOs and unlabeled hCOs.
(E) Immunostaining for mCherry, the neuronal marker MAP2, and the progenitor marker PAX6 in hThCAs at 14 dpf. Scale bars represent 200 µm.
(F) Three-dimensional immunostaining for mCherry (left) and quantification of the location of mCherry^+^ axons within the hCOs of hThCAs at 14 dpf (right). Scale bar represents 500 µm. Data are expressed as mean ± SEM: n = 5 organoids. ****p* < 0.001, Welch’s *t*-test.
(G) Schematic of each region and quantification of the proportion of mCherry^+^ thalamic axons in each region. Data are expressed as ± SEM: n = 8 hThCAs. ****p* < 0.001, Dunnett’s test.
(H) Representative image of hThCAs at 14 dpf stained for EYFP and mCherry. Scale bars represent 500 µm.
(I) Generation of human corticocortical assembloids (hCCAs) from EYFP-expressing and unlabeled hCOs.
(J) EYFP-expressing axonal projections from hCOs to the other side within hThCAs and hCCAs at 21 dpf. Scale bars represent 500 µm.
(K) Quantification of EYFP^+^ migrating cells from hCOs to the other side within hThCAs or hCCAs. Data are expressed as ± SEM: n = 8 hThCAs, n = 9 hCCAs. **p* < 0.05, Welch’s *t*-test.

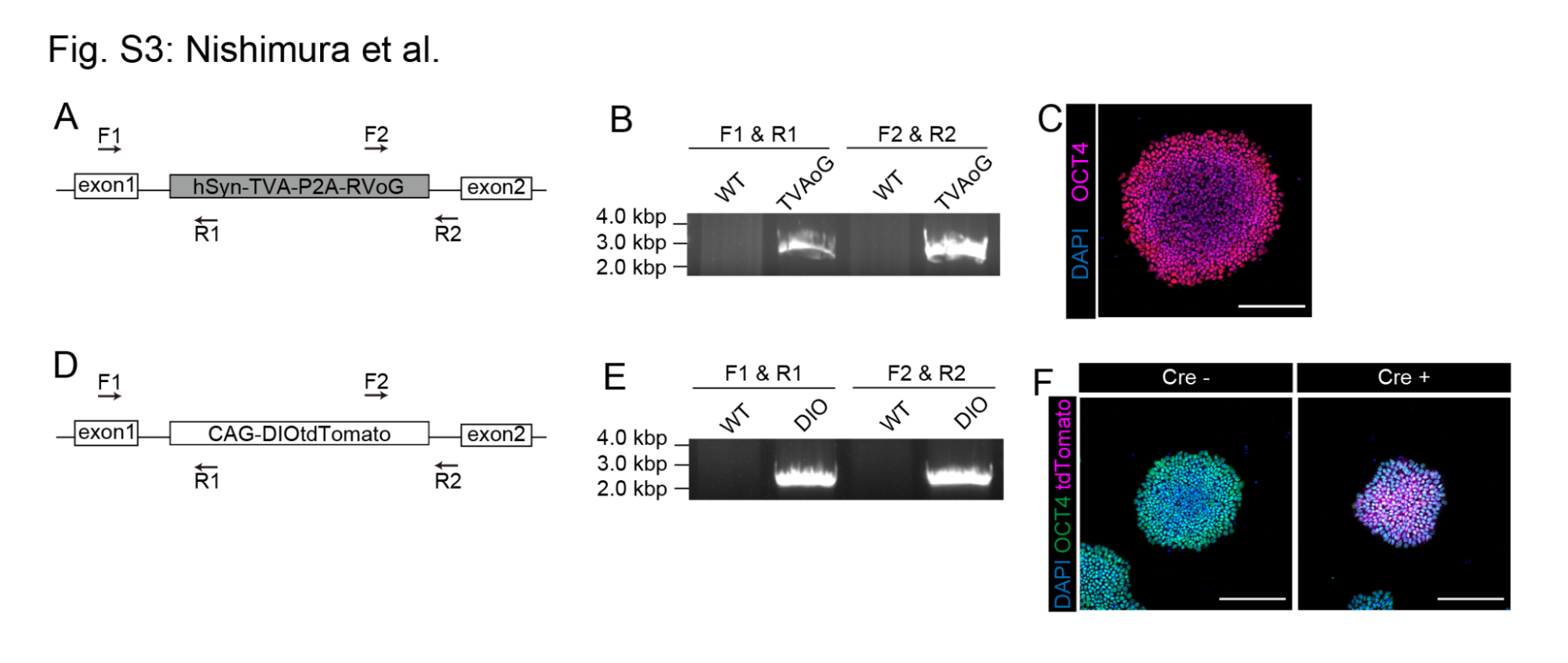

### Figure S3. Generation of hiPSCs-TVA-oG and hiPSCs-DIO-tdTomato knocked-in lines.

(A) Primers for genotyping the hiPSCs-TVA-P2A-oG line. Arrows, PCR primers.
(B) Electrophoresis images of the PCR products for the hiPSCs-TVA-P2A-oG line.
(C) Immunostaining for the pluripotent marker OCT4 in the hiPSCs-TVA-P2A-oG line. Scale bars represent 200 µm.
(D) Primers for genotyping the hiPSCs-DIO-tdTomato line. Arrows, PCR primers.
(E) Electrophoresis images of the PCR products for the knocked-in hiPSC lines.
(F) Immunostaining for the pluripotent marker OCT4 and tdTomato in the hiPSCs-DIO-tdTomato line with or without Cre. Scale bars represent 200 µm.

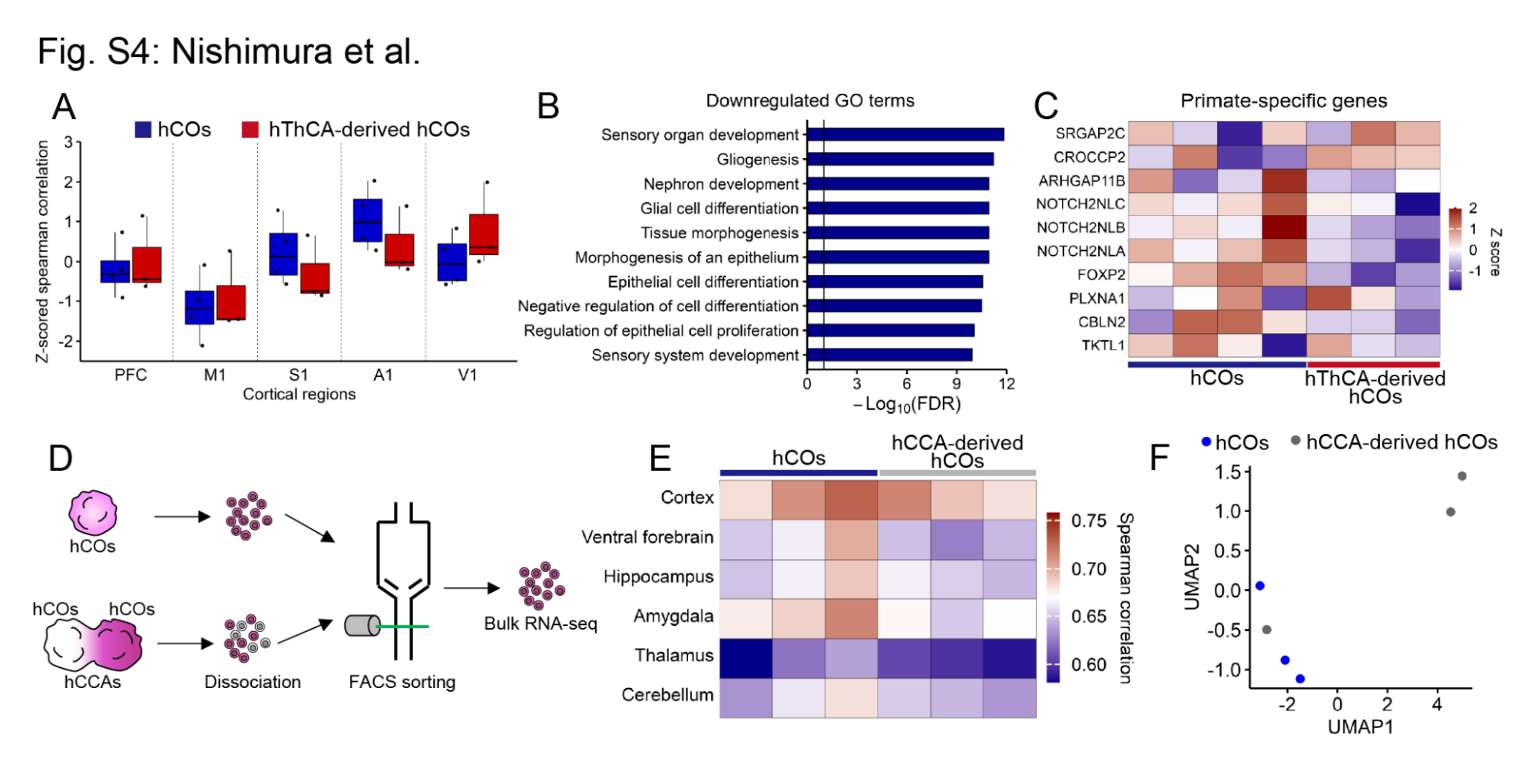

### Figure S4. Characterization of transcriptomes in hCOs and assembloid-derived hCOs.

(A) Z-scored spearman correlation between the transcriptomes of each region of human fetal cortex (12–17 pcw) and those of hCOs or hThCA-derived hCOs. n= 4 hCOs, n = 3 hThCA-derived hCOs.
(B) Top 10 gene ontology terms downregulated in the hThCA-derived hCOs. Vertical line represents −log_10_FDR = 1.
(C) Heatmap showing the expression of primate-specific genes in the hCOs and hThCA-derived hCOs. n= 4 hCOs, n = 3 hThCA-derived hCOs.
(D) Bulk RNA sequencing method for hCOs and human corticocortical assembloids (hCCA)-derived hCOs.
(E) Heatmap showing the Spearman correlations between the transcriptomes of region-specific human fetal brains and those of hCOs or hCCA-derived hCOs. n= 3 hCOs, n = 3 hCCA-derived hCOs.
(F) UMAP for the transcriptomes of hCOs and hCCA-derived hCOs. n= 3 hCOs, n = 3 hCCA-derived hCOs.

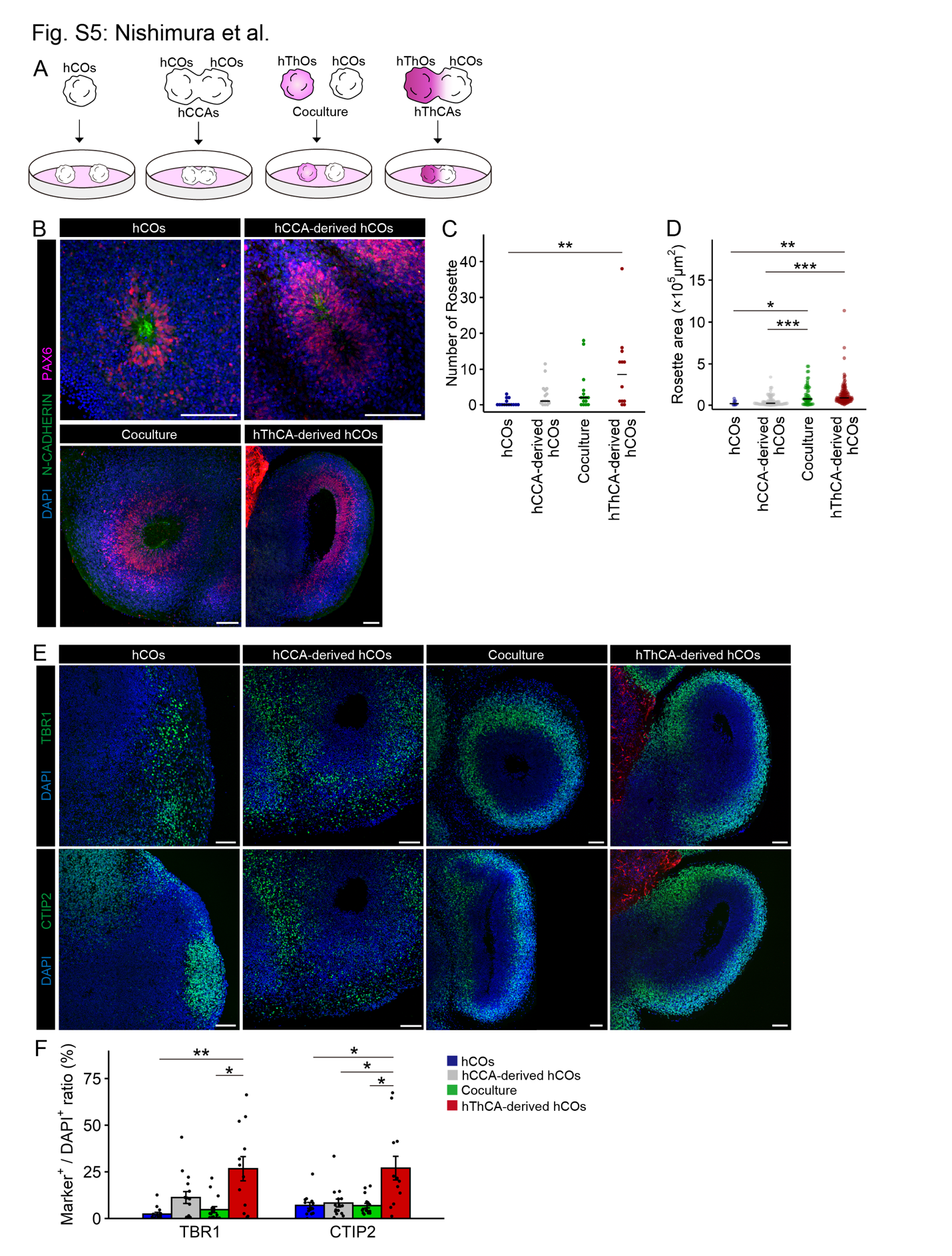

### Figure S5. Cytoarchitecture in isolated hCOs, hCCA-derived hCOs, cocultured hCOs, and hThCA-derived hCOs.

(A) Schematic of each culture condition.
(B) Immunostaining for N-CADHERIN and PAX6 in each hCOs on day 80. Scale bars represent 100 µm.
(C) Quantification of the number of rosettes in each hCOs. n = 14 hCOs, n = 15 hCCA-derived hCOs, n = 15 Coculture, and n = 12 hThCAs-derived hCOs. **p* < 0.05, Dunn test.
(D) Quantification of the area of each rosette in each hCOs. n = 8 rosettes (hCOs), n = 87 rosettes (hCCA-derived hCOs), n = 59 rosettes (Coculture), and n = 113 rosettes (hThCAs-derived hCOs). **p* < 0.05, ***p* < 0.01, ****p* < 0.001, Dunn test.
(E) Immunostaining for TBR1 and CTIP2 in each hCOs on day 80. Scale bars represent 100 µm.
(F) Quantification of TBR1 and CTIP2 expression in each hCOs. Data are expressed as mean ± SEM. n = 14 hCOs, n = 15 hCCA-derived hCOs, n = 16 Coculture, and n = 12 hThCAs-derived hCOs. **p* < 0.05, ***p* < 0.01, Dunn test.

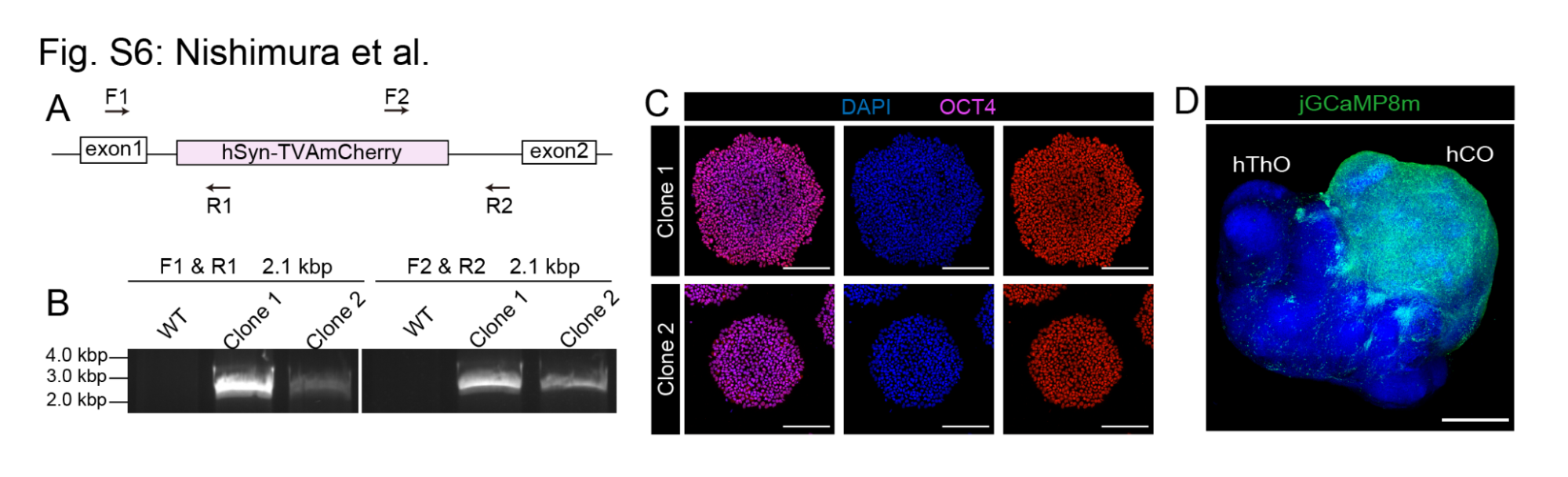

### Figure S6. Generation of the hiPSCs-TVAmCherry line.

(A) Primers for genotyping the hiPSCs-TVAmCherry line. Arrows, PCR primers.
(B) Electrophoresis images of the PCR products for the knocked-in hiPSC lines.
(C) Immunostaining for the pluripotent marker OCT4 in the hiPSCs-TVAmCherry line. Scale bars represent 200 µm.
(D) Restricted jGCaMP8m expression in cortical neurons of the hThCAs on day 54. Scale bar represents 1,000 µm.

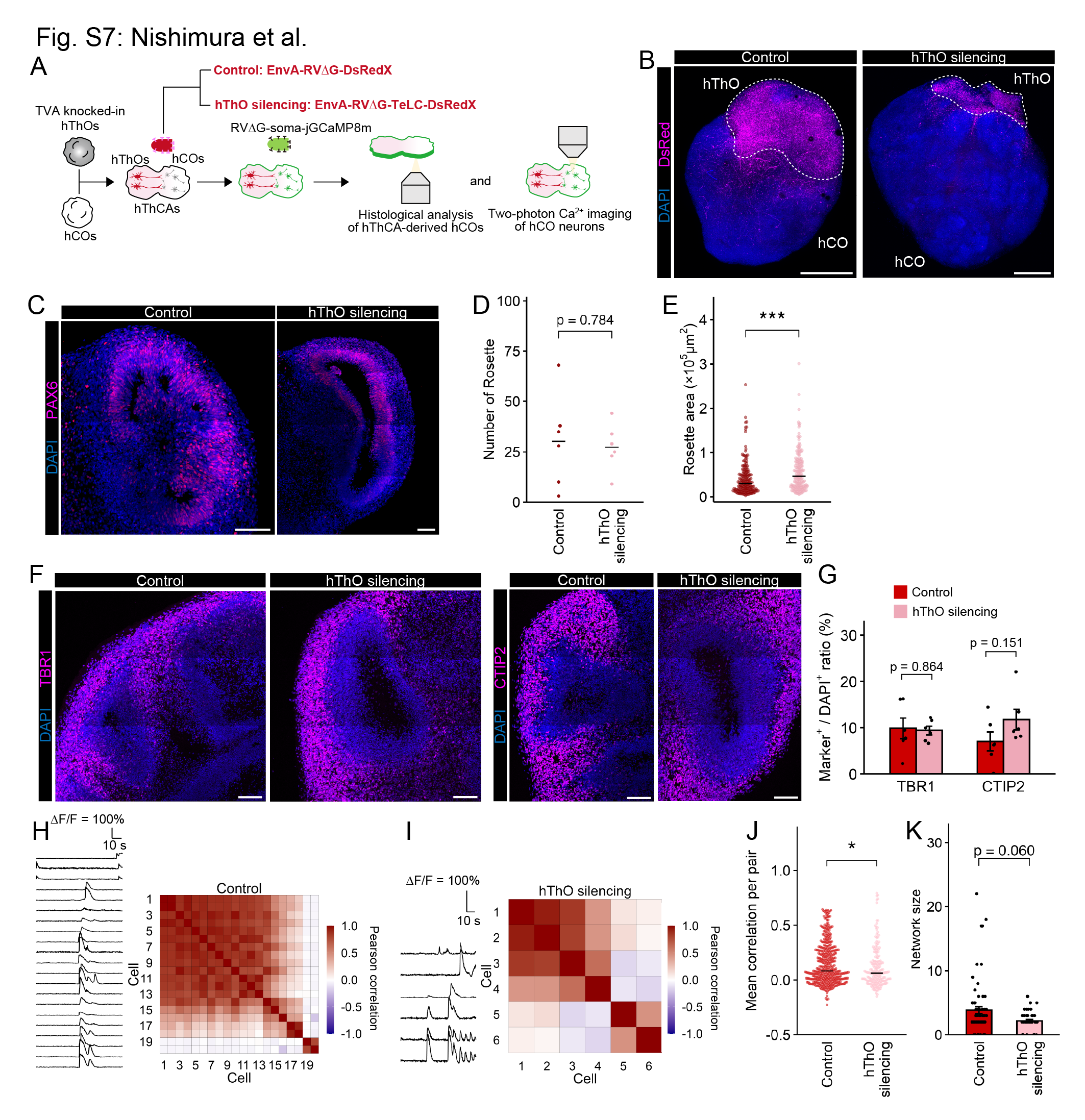

### Figure S7. Silencing of hThCA-derived hThOs activity.

(A) Experimental design for silencing of hThCA-derived hThOs activity.
(B) Restricted DsRed expression in thalamic neurons of the hThCAs on day 101. Scale bars represent 1,000 µm.
(C) Immunostaining for PAX6 in hThCAs in control and hThO silencing conditions on day 83. Scale bars represent 100 µm.
(D) Quantification of the number of rosettes in hThCA-derived hCOs in each condition. n = 6 hThCA-derived hCOs (Control and hThO silencing). Welch’s *t*-test.
(E) Quantification of the area of each rosette in hThCA-derived hCOs in each condition. n = 183 rosettes (Control), n = 164 rosettes (hThO silencing). ****p* < 0.001, Mann–Whitney U test.
(F) Immunostaining for TBR1 and CTIP2 in hThCAs in each condition on day 83. Scale bars represent 100 µm.
(G) Quantification of TBR1 and CTIP2 expression hThCA-derived hCOs in each condition. Data are expressed as mean ± SEM. n = 6 hThCA-derived hCOs (Control and hThO silencing). Welch’s *t*-test.
(H) Representative time-series traces of jGCaMP8m signal changes (left) and heatmap showing correlation coefficients of activity between all pairs of simultaneously recorded neurons (right) in hThCA-derived hCOs in the control condition.
(I) Representative time-series traces of jGCaMP8m signal changes (left) and heatmap showing correlation coefficients of activity between all pairs of simultaneously recorded neurons (right) in hThCA-derived hCOs in the hThO silencing condition.
(J) Quantification of mean correlation coefficients of activity between all pairs of simultaneously recorded neurons from hThCA-derived hCOs in each condition. Red dots, correlation of each cell pair; black line, median. n = 389 pairs (Control), n = 179 pairs (hThO silencing), **p* < 0.05, Mann–Whitney U test.
(K) Quantification of the network synchronization size in hThCA-derived hCOs. Mann–Whitney U test. Data are expressed as mean ± SEM. n = 69 network (Control), n = 41 network (hThO silencing). Mann–Whitney U test.

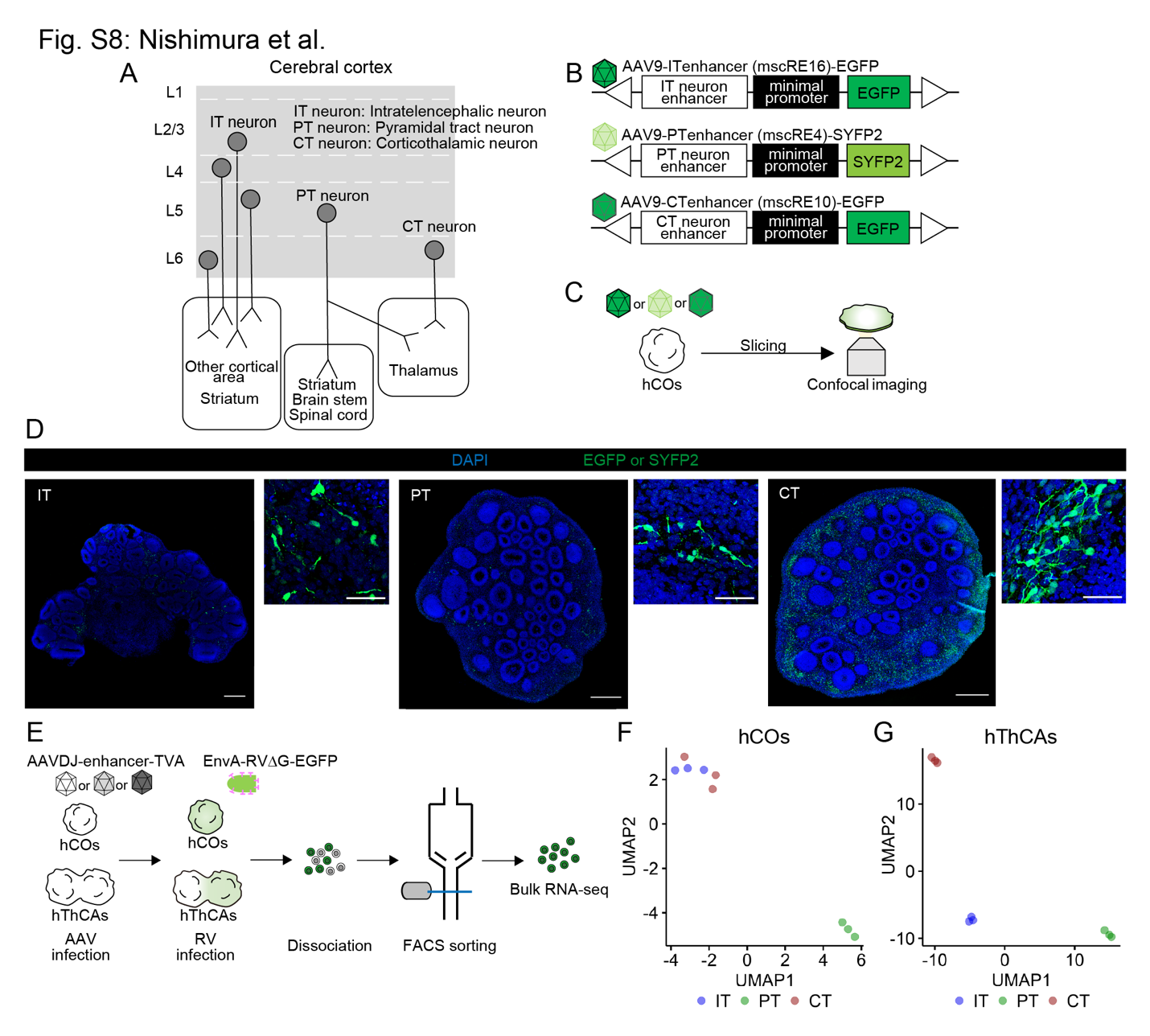

### Figure S8. Validation of excitatory projection neuron subtypes in isolated hCOs and hThCA-derived hCOs.

(A) Excitatory projection neuron subtypes in the cerebral cortex.
(B) AAVs that encode a fluorescence protein under the cell type-specific enhancers mscRE16 (IT neurons: intratelencephalic neurons), mscRE4 (PT neurons: pyramidal tract neurons), and mscRE10 (CT neurons: corticothalamic neurons).
(C) Schematic representation of viral labeling and confocal imaging in hCOs.
(D) Immunostaining for GFP or SYFP2 in hCOs on day 52. hCOs were infected with AAV9-ITenhancer-EGFP (left), AAV9-PTenhancer-SYFP2 (middle), and AAV9-CTenhancer-EGFP (right). Scale bars represent 500 µm (left), 100 µm (right).
(E) Bulk RNA sequencing method for each cortical subtype in isolated hCOs and hThCA-derived hCOs.
(F) UMAP for the transcriptomes of each cortical subtype in isolated hCOs. n = 3 hCOs (IT, PT, and CT).
(G) UMAP for the transcriptomes of each cortical subtype in hThCA-derived hCOs. n = 3 hThCA-derived hCOs (IT, PT, and CT).

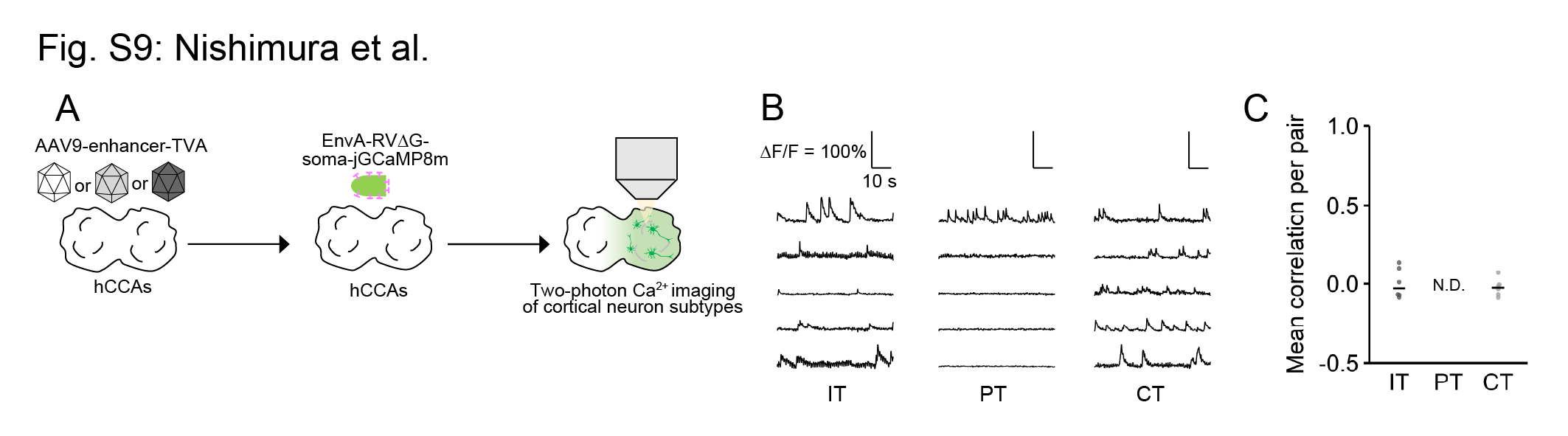

### Figure S9. Cell type-specific two-photon Ca^2+^ imaging of hCCA-derived hCOs.

(A) Method for cell type-specific two-photon Ca^2+^ imaging of hCCA-derived hCOs.
(B) Representative time-series traces of jGCaMP8m signal changes of each cortical subtype in hCCA-derived hCOs.
(C) Quantification of the mean correlation coefficients of the activity between all pairs of simultaneously recorded IT, PT, or CT neurons from hCCA-derived hCOs. n = 6 pairs (IT and CT).

### Table S1. List of Knocked-in check primers.

| **Purpose of use** | **Sequence** |
| --- | --- |
| Knocked-in check F1 primer for all hiPSCs | 5′- GACCTGGACGAGGCGCGTCTGATGC -3′ |
| Knocked-in check F2 primer for CAG-mCherry and CAG-EYFP hiPSCs | 5′- TTCTGAGCAGACATGATAAGATACATTGATGAGTTTGGACAACCAC -3′ |
| Knocked-in check F2 primer for hSyn-TVA-P2AoG, CAG-DIO-tdTomato, and hSyn-TVAmCherry hiPSCs | 5′- CTCGACGTTGTCACTGAAGCGGGAAGGGAC -3′ |
| Knocked-in check R1 primer for CAG-mCherry, CAG-EYFP, and CAG-DIO-tdTomato hiPSCs | 5′- ACGTGGGGCTCACCTCGACCATGG -3′ |
| Knocked-in check R1 primer for hSyn-TVA-P2AoG and hSyn-TVAmCherry hiPSCs | 5′- CGACTGCGCTCTCAGGCACGAC -3′ |
| Knocked-in check R2 primer for all hiPSCs | 5′- CAACCCCCTAGCCACTAAGGCAATTGGGGTG -3′ |

### Table S2. List of qPCR primers.

| **Gene** | **Sequence** |
| --- | --- |
| Fw primer for *FOXG1* | 5′- ATGATCCCCAAGTCCTCGTT -3′ |
| Rv primer for *FOXG1* | 5′- GTGGTGGTTGTCGTTCTGG -3′ |
| Fw primer for *EMX2* | 5′- GGTCATCGCTTCCAAGGGAAC -3′ |
| Rv primer for *EMX2* | 5′- GGCGTGTTCCAGCCTTAGAA -3′ |
| Fw primer for *OTX2* | 5′- ACCTTGAACTCCACCT -3′ |
| Rv primer for *OTX2* | 5′- GCTTCTCTTCTCTGAC -3′ |
| Fw primer for *TCF7L2* | 5′- ACACCAGATTGCAGTTCAGTAGC -3′ |
| Rv primer for *TCF7L2* | 5′- AGACAATGTGTGCCGGTGAT -3′ |
| Fw primer for *DBX1* | 5′- GAGCAGTCTTCTCCGACGTG -3′ |
| Rv primer for *DBX1* | 5′- TTTCATGCGTCGGTTCTGGA -3′ |
| Fw primer for *GBX2* | 5′- AAAGAGGGCTCGCTGCTC -3′ |
| Rv primer for *GBX2* | 5′- ATCGCTCTCCAGCGAGAA-3′ |
| Fw primer for *TBR1* | 5′- GACTCAGTTCATCGCCGTCA -3′ |
| Rv primer for *TBR1* | 5′- GCCGGTGTAGATCGTGTCAT -3′ |
| Fw primer for *BCL11B* | 5′- GCCAGTGTCAGTTGTCAGGTAA -3′ |
| Rv primer for *BCL11B* | 5′- CTCCAGGTAGATGCGGAAGC -3′ |
| Fw primer for *OLIG3* | 5′- CCTGCTCGCCAGAAACTACA -3′ |
| Rv primer for *OLIG3* | 5′- CCCCCATAGATCTCGCCAAC -3′ |
| Fw primer for *SLC17A6* | 5′- GCAGCCAACAGGGTTTTCG -3′ |
| Rv primer for *SLC17A6* | 5′- ACATGCTGGGTAGGTCACAC -3′ |

### Table S3. List of antibodies.

| **Antibodies** | **Source** | **Identifier** | **Dilution** |
| --- | --- | --- | --- |
| Rabbit Anti-OCT4 | Cell Signaling Technology | Cat# 2890  RRID:AB_2167725 | 1:2000 |
| Rabbit Anti-FOXG1 | Takara Bio | Cat# M227  RRID:AB_2827749 | 1:500 |
| Mouse Anti-N-CADHERIN | Cell Signaling Technology | Cat# 14215  RRID:AB_2798427 | 1:200 |
| Rabbit Anti-PAX6 | Abcam | Cat# ab195045  RRID:AB_2750924 | 1:600 |
| Chicken Anti-MAP2 | Abcam | Cat# ab5392  RRID:AB_2138153 | 1:500 |
| Rat Anti-CTIP2 | Abcam | Cat# ab18465  RRID:AB_2064130 | 1:500 |
| Rabbit Anti-TBR1 | Cell Signaling Technology | Cat# 49661  RRID:AB_2799364 | 1:400 |
| Rabbit Anti-SATB2 | Abcam | Cat# ab92446  RRID:AB_10563678 | 1:500 |
| Rabbit Anti-TCF7L2 | Cell Signaling Technology | Cat# 2569  RRID:AB_2199816 | 1:500 |
| Rabbit Anti-Ki67 | Cell Signaling Technology | Cat# 9129  RRID:AB_2687446 | 1:400 |
| Rabbit Anti-TBR2 | Abcam | Cat# ab23345  RRID:AB_778267 | 1:500 |
| Chicken Anti-GFP | Abcam | Cat# ab13970  RRID:AB_300798 | 1:1,000 |
| Rat Anti-mCherry | Thermo Fisher Scientific | Cat# M11217  RRID:AB_2536611 | 1:1,000 |
| GuineaPig Anti-vGluT1 | Millipore | Cat# AB5905  RRID:AB_2301751 | 1:200 |
| Rabbit Anti-DsRed | Takara Bio | Cat# 632496  RRID:AB_10013483 | 1:1000 |
| Goat Anti-vGluT2 | Frontier Institute Co. | Cat# Af310  RRID:AB_2571620 | 1:200 |
| Rabbit Anti-HOMER1 | Synaptic Systems | Cat# 160 003  RRID:AB_887730 | 1:200 |
| Alexa Fluor 488-AffiniPure Donkey Anti-Mouse IgG (H+L) | Jackson ImmunoResearch Labs | Cat# 715-545-151  RRID:AB_2341099 | 1:1,000 |
| Alexa Fluor 594 AffiniPure Donkey Anti-Rabbit IgG (H+L) | Jackson ImmunoResearch Labs | Cat# 711-585-152  RRID:AB_2340621 | 1:1,000 |
| Alexa Fluor 488 AffiniPure Donkey Anti-Chicken IgY (IgG) (H+L) | Jackson ImmunoResearch Labs | Cat# 703-545-155  RRID:AB_2340375 | 1:1,000 |
| Alexa Fluor 488-AffiniPure Donkey Anti-Rat IgG (H+L) | Jackson ImmunoResearch Labs | Cat# 712-545-153  RRID:AB_2340684 | 1:1,000 |
| Alexa Fluor 594-AffiniPure Donkey Anti-Rat IgG (H+L) | Jackson ImmunoResearch Labs | Cat# 712-585-153  RRID:AB_2340689 | 1:1,000 |
| Alexa Fluor 647-AffiniPure Donkey Anti-Rabbit IgG (H+L) | Jackson ImmunoResearch Labs | Cat# 711-605-152  RRID:AB_2492288 | 1:1,000 |
| Alexa Fluor 488-AffiniPure Donkey Anti-Guinea Pig IgG (H+L) | Jackson ImmunoResearch Labs | Cat# 706-545-148  RRID:AB_2340472 | 1:1,000 |
| Alexa Fluor 488-AffiniPure Donkey Anti-Rabbit IgG (H+L) | Jackson ImmunoResearch Labs | Cat# 711-545-152  RRID:AB_2313584 | 1:1,000 |
| Alexa Fluor 647-AffiniPure Donkey Anti-Goat IgG (H+L) | Jackson ImmunoResearch Labs | Cat# 705-605-147  RRID:AB_2340437 | 1:1,000 |
| Alexa Fluor 647-AffiniPure Donkey Anti-Chicken IgY (IgG) (H+L) | Jackson ImmunoResearch Labs | Cat# 703-605-155  RRID:AB_2340379 | 1:1,000 |
| Alexa Fluor 647-AffiniPure Donkey Anti-Rabbit IgG (H+L) | Jackson ImmunoResearch Labs | Cat# 711-605-152  RRID:AB_2492288 | 1:1,000 |
| Alexa Fluor 647 AffiniPure Donkey Anti-Rat IgG (H+L) | Jackson ImmunoResearch Labs | Cat# 712-605-153  RRID:AB_2340694 | 1:1,000 |

### Table S4. Titers of viral vectors.

| **Viral vectors** | **Titer** |
| --- | --- |
| AAVDJ-CaMKIIa-jGCaMP7f | 5.89 × 10^11^ viral genomes/mL |
| AAV9-3×(core)mscRE4-minCMV-SYFP2 | 3.27 × 10^12^ viral genomes/mL |
| AAV9-mscRE16-minBGpromoter-EGFP | 3.78 × 10^13^ viral genomes/mL |
| AAV9-mscRE10-minBGpromoter-EGFP | 1.76 × 10^11^ viral genomes/mL |
| AAV9-3×(core)mscRE4-minCMV-TVA | 0.482–3.21 × 10^13^ viral genomes/mL |
| AAV9-mscRE16-minBGpromoter-TVA | 0.206–1.83 × 10^13^ viral genomes/mL |
| AAV9-mscRE10-minBGpromoter-TVA | 0.709–2.97 × 10^13^ viral genomes/mL |
| EnvA-RV∆G-GFP-iCre | 2.00 × 10^7^–2.2 × 10^10^ infectious units/mL |
| EnvA-RV∆G-DsRedX | 1.30 × 10^9^ infectious units/mL |
| EnvA-RV∆G-TeLC-DsRedX | 1.44 × 10^8^ infectious units/mL |

### Table S5. The time window of analysis.

| **Analysis** | **Time window** |
| --- | --- |
| Differentiation into hCOs and hThOs (Fig. S1) | Days 42–50 |
| Axonal projection (Fig.1 and Fig. S2) | Days 29–45 |
| Synaptic connection (Fig. 1) | Days 45–74 |
| RNA sequencing (Fig. 2 and Fig. S4) | Days 70–83 |
| RNA sequencing of cortical subtypes (Fig. S8) | Days 113–162 |
| Histology (Fig. 3 and Fig S5) | Day 80 |
| Histology of hThO silencing (Fig. S7) | Day 83 |
| Functional imaging (Fig. 4) | Days 70–84 |
| Functional imaging of hThO silencing (Fig. S7) | Days 83–101 |
| Functional imaging of cortical subtypes (Fig. 5 and Fig. S9) | Days 67–89 |
